# PCR data accurately predict infectious virus: a characterization of SARS-CoV-2 in non-human primates

**DOI:** 10.1101/2023.06.23.546114

**Authors:** Celine E. Snedden, James O. Lloyd-Smith

## Abstract

Researchers and clinicians often rely on molecular assays like PCR to identify and monitor viral infections instead of the resource-prohibitive gold standard of viral culture. However, it remains unclear when (if ever) PCR measurements of viral load are reliable indicators of replicating or infectious virus. Here, we compare total RNA, subgenomic RNA, and viral culture results from 24 studies of SARS-CoV-2 in non-human primates using bespoke statistical models. On out-of-sample data, our best models predict subgenomic RNA from total RNA with 91% accuracy, and they predict culture positivity with 85% accuracy. Total RNA and subgenomic RNA showed equivalent performance as predictors of culture positivity. Multiple cofactors, including exposure conditions and host traits, influence culture predictions for total RNA quantities spanning twelve orders of magnitude. Our model framework can be adapted to compare any assays, in any host species, and for any virus, to support laboratory analyses, medical decisions, and public health guidelines.

## Introduction

Assays that detect and quantify the presence of viral genetic material are invaluable tools for clinicians, virologists, and epidemiologists, since they are used to identify infections, monitor individual infection trajectories, and track population-wide disease trends. The global reliance on quantitative reverse transcription-polymerase chain reaction (RT-qPCR) during the COVID-19 pandemic underscores its importance as a fast, sensitive, and relatively inexpensive mainstay of research and public health. Yet positive RT-qPCR results do not necessarily indicate active infection or viral shedding because these assays only target and quantify viral genomic material.^1, 2^ Viral culture is the gold-standard method to detect infectious virus, but it is slow, labor-intensive, and requires niche resources like permissive cells and biosafety facilities. This precludes its use as a primary diagnostic in public health crises or even in standard clinical and research practices where speed and accessibility matter. The development of alternate methods to accurately characterize infectiousness is an active priority.

Seeking a culture-free method to identify replicating virus, many studies on SARS-CoV-2 developed alternative RT-qPCR assays based on coronavirus transcription mechanisms. Within host cells, coronaviruses transcribe not only full-length genomic RNA (gRNA) but also multiple subgenomic RNAs (sgRNA), which are a nested set of RNA segments that function as mRNA for translation of some structural and accessory proteins.^3^ Standard RT-qPCR protocols^4^ typically amplify both gRNA and sgRNA simultaneously (henceforth termed a total RNA assay and abbreviated to ‘totRNA’). Since sgRNAs are only transcribed after cellular entry and are generally not packaged into mature virions,^5^ sgRNA-specific assays for SARS-CoV-2 were developed as a proxy for replicating virus,^6^ and they have been used widely, including to distinguish between replicating virus and residual inoculum in animal challenge experiments.^7, 8^

Despite the popularity of sgRNA assays, their diagnostic utility relative to totRNA or gRNA assays is debated. Based on findings that sgRNA degrades faster than gRNA^8^, is not found in virions^5^, and correlates better with viral culture results,^9–12^ some consider sgRNA a better indicator of recent replication and infectiousness.^6, 13^ Others dispute these claims based on contrary findings, including evidence of similar degradation rates between sgRNA and gRNA,^14–16^ the discovery of membrane-associated and nuclease-resistant sgRNA,^14^ and analyses showing that sgRNA does not correlate better with culture outcomes.^17^ Studies finding that sgRNA quantities scale linearly with totRNA prompted further claims that sgRNA quantification offers no additional value relative to totRNA,^15–17^ and skeptics have argued that any improved correlation between sgRNA and culture likely reflects the assay’s lower sensitivity rather than true biological signal.^14–16^ Meanwhile, samples with large quantities of totRNA but undetectable sgRNA or unculturable virus are widely evident in the literature, especially in animal challenge experiments, but they go largely unexplained.^8, 18^ These patterns highlight the complexity of the relationships among PCR assays and viral culture, and they underscore that our understanding of their relative trajectories during infection remains incomplete. Given their foundational importance for research and healthcare, many studies have called for better methods to interpret these assays and their interrelationships.^19–22^

Data limitations are central to these unresolved debates on how well PCR predicts culture and whether that varies by RNA type since the generalizability of observed patterns remains unclear. Each study’s sample size is typically quite small (e.g., often less than 100 RNA-positive samples), protocols differ between studies (e.g., PCR target genes, cell lines), patient demographics vary (e.g., hospitalized patients versus routine screening of university students), and analytical methods differ (e.g., descriptive statistics, logistic or linear regressions). Further unexplained variation may depend on patients’ age, sex, and comorbidities, which can affect infection outcomes^23–25^ but are often unaccounted for in assay comparisons. Exposure route and dose are also unknown for clinical infections, and because the true infection time is unknown, analyses of clinical data must rely on metrics like time since symptom onset,^6, 15, 20, 26, 27^ for which individual heterogeneity and recall bias can introduce substantial noise. Despite substantial effort to correlate RNA presence with culture outcomes, no study has yet jointly evaluated these various cofactors to identify and quantify their effects, and thus no best practices exist to predict an individual’s infectiousness.

In this study, we compiled and jointly analyzed a database of non-human primate (NHP) experiments, including 24 articles that reported per-sample measurements of at least two of the following assays: totRNA, sgRNA, and viral culture. This meta-analytic design enabled larger sample sizes and knowledge of variables that are unknowable with clinical data (i.e., exposure time and conditions), all for a gold-standard animal model of human disease.^28^ We developed a Bayesian hurdle model to predict the results from these disparate assays and to evaluate the effects of NHP species, demographic characteristics (age, sex), exposure conditions (dose and route), time since infection, and study protocols (sample type, target gene, cell line, culture assay) on the relationships among assay outcomes. We first applied this method to predict sgRNA results from totRNA results, which enabled us to reconstruct their relative trajectories for all included individuals. Then, we tested the ability of both PCR assays to predict viral culture results. We characterized model performance on withheld data to evaluate predictive accuracy and generalizability. Our results have many applications, including: (i) to improve PCR-based clinical decisions with more accurate prediction of infectious status, (ii) to support literature syntheses and further model analyses by enabling quantitative comparison of SARS-CoV-2 infection dynamics across studies, and (iii) to offer a standardized, adaptable framework for quantitative comparison of any assays, in any host species, and for any virus.

## Methods

### Database compilation

Following the PRISMA guidelines for systematic literature searches,^29^ we constructed a comprehensive database of SARS-CoV-2 viral load and infectious virus data from non-human primate experiments (**Figure S1**). To be included, articles were required to: (i) experimentally infect rhesus macaques (*Macaca mulatta*), cynomolgus macaques (*Macaca fascicularis*), or African green monkeys (*Chlorocebus sabaeus*) with SARS-CoV-2 (restricted to basal strains, excluding the D614G mutation or other named variant), and (ii) report quantitative or qualitative measurements of viral load (measured by RT-qPCR) or infectious virus (measured by plaque assay or endpoint titration) from at least one biological specimen for at least one individual and at least one sample time post infection. Only individuals receiving no or placebo treatments were recorded.

Of 86 studies meeting these criteria, we used the 24 articles that reported at least two of the following assays: totRNA PCR, sgRNA PCR, or viral culture (**Table S1, Figure S1**).^7, 8, 18, 30–50^ Raw data were used when available (published or obtained via email correspondence); otherwise, one author (CES) extracted data from published figures using the package ‘digitize’^51^ in R.^52^ Additional details of data acquisition and standardization are described in the **Supplementary Methods.**

### Bayesian hurdle model framework

To compare disparate assays, we developed a Bayesian hurdle model with two components: (i) a logistic regression that predicts whether assay Y will fall above the limit of detection (Y_>LOD_) based on assay X, and (ii) a linear regression that describes the quantitative relationship between X and Y when both are measurable (Y_value_) (**Figure S2**). Each component may include distinct sets of additional predictor variables (A_i_ and B_j_, respectively). For the linear component, we incorporated hierarchical errors such that the model estimates article-specific error distributions (σ_a_) based on distributions of population average errors 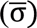 and error standard deviations (σ_sd_). This captures potential differences in experimental noise among studies and protocols. The basic form of this model is as follows, where δ and β are regression coefficients associated with the predictors noted in the subscript:

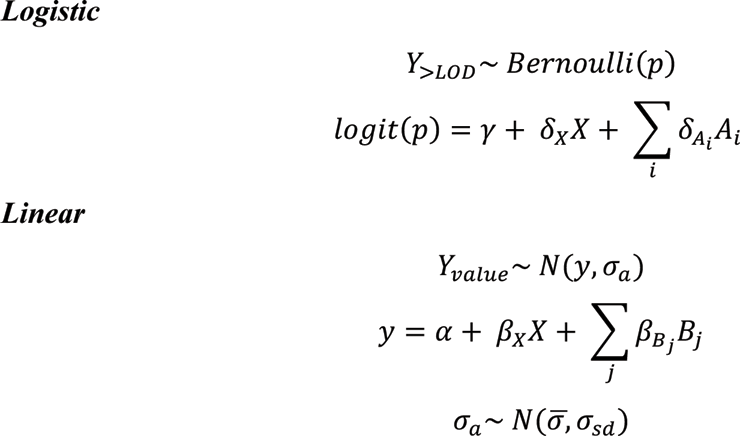

We evaluated the predictive performance of multiple models with different combinations of candidate predictors, and so the ∑δ_Ai_A_i_ and ∑β_Bj_B_j_ terms varied for each considered model. Categorical predictors with more than two classifications were treated as unordered index variables, while binary predictors were treated as indicator variables. For instances of unknown age or sex, we marginalized over all possibilities.

We first applied this framework to predict sgRNA from totRNA results (termed the ‘sgRNA model’). All totRNA-negative samples are predicted to be sgRNA-negative, by definition. We then predicted viral culture results from PCR data using a parallel framework (termed the ‘culture model’), with the following minor modifications: (i) we considered models depending on totRNA, sgRNA, or both as predictors, and (ii) we restricted analyses to the logistic component, given scarcity of quantitative culture results. The model predicts all RNA-negative samples are culture negative.

### Candidate predictor selection and prior sensitivity analyses

All candidate predictors were included because of hypothesized effects on the relationships among assay results, as summarized below. We chose informative priors to rule out implausible parameter values and to reflect existing knowledge on the expected direction of individual effects (outlined in the **Supplementary Methods**), where appropriate. Notably, prior predictive simulations confirmed variable but reasonable *a priori* expectations for these informative priors, with substantial improvement over non-informative priors that do not reflect existing knowledge (**Figure S12**). Parameter estimates for the best models were qualitatively similar between informative and non-informative priors (**Figure S12**).

All considered models included totRNA, sgRNA, or both as the primary predictor(s). For all models, we considered multiple demographic factors including age class, sex, and non-human primate species, given hypothesized effects on SARS-CoV-2 infection.^23–25, 38, 53, 54^ Because exposure conditions can affect initial virion and totRNA quantities, we included inoculation dose (in log10 pfu) and day post infection as candidate predictors. For day post infection, we distinguished between inoculated tissues sampled on the first day versus all other days post infection, and non-inoculated tissues on any day post infection (see **Table S9** for tissue-specific categorization). Because sample content and processing may vary between non-invasive (e.g., swabs) and invasive samples (e.g., whole tissues obtained at necropsy), we considered sample type as a binary predictor. We also included predictors to account for assay-specific variation. For sgRNA models, we derived a target gene predictor based on the expected number of transcripts available for amplification during each PCR protocol, given that sgRNA abundance varies by gene^55^ and totRNA assays can amplify both genomic and subgenomic RNA. We distinguished between totRNA assays that amplify most (‘totRNA-high’; targeting the N gene) or few sgRNA species (‘totRNA-low’; E gene) and sgRNA assays that target highly expressed (‘sgRNA-high’; sgN) or less expressed sgRNA species (‘sgRNA-low’; sgE, sg7), resulting in four possible protocol combinations. For culture models, we used the totRNA target gene as the predictor, except for the models including only sgRNA as the primary predictor. Since viral infectivity varies among cell lines^19, 56, 57^ and culture sensitivity differs between endpoint dilution and plaque assays,^58^ we included cell line and culture assay as additional predictors for culture.

### Evaluating and comparing model performance

To find the highest performing model for each investigation, we first used a forward search to identify the model with the best performance for each possible number of predictors. We used 10-fold cross-validation to evaluate each model’s predictive performance on withheld data, and for each stage we selected the predictor that most increased the expected log pointwise predictive density (ELPD).^59^ Following convention, we considered an ELPD difference of less than 4 to be small when comparing two models.^59^ Of those models identified by the forward search, we selected the ‘best model’ as the one with fewest predictors that achieved similar or better performance compared to the ‘full model’ (containing all predictors) on out-of-sample (test) data for three relevant statistics: (i) ELPD, (ii) prediction accuracy (i.e., the percent of correctly classified samples for the logistic component, or the percent of samples where the observed value fell within the 50% prediction interval for the linear component), and (iii) Matthew’s correlation coefficient^60^ (MCC; logistic components) or median absolute error on the posterior predictive medians (MAE; linear component). Comprehensive descriptions of model evaluation and selection are provided in the **Supplementary Methods**.

### Computational methods and software

All data preparation, analysis, and plotting were completed with R version 4.2.0.^52^ Posterior sampling of the Bayesian model was performed with No-U-Turn Sampling (NUTS) via the probabilistic programming language Stan^61^ using the interface CmdStanR version 0.5.2. All model fits were generated by running six replicate chains with 4000 iterations each, of which the first 2000 iterations were treated as the warmup period and the final 2000 iterations were used to generate parameter estimates. Model convergence was assessed by the sampling software using *R̂*, effective sample sizes, and other diagnostic measures employed by CmdStan by default. No issues were detected.

## Results

### The compiled dataset includes 2167 samples from 174 individual non-human primates

A comprehensive literature search for studies that challenged non-human primates with SARS-CoV-2 identified 24 articles that reported per-sample measurements of at least two of the following assays: totRNA RT-qPCR, sgRNA RT-qPCR, and viral culture (**Figure S1**; Tables 1, **S1**). Of those, 14 articles reported totRNA and sgRNA for 116 individuals and 1194 samples, and 15 articles reported viral culture and either RNA type for 90 individuals and 1315 samples. Five articles reported results for all three assays, totaling 342 such samples.

The dataset includes various demographic groups, including both sexes, ages ranging from 1 to 22 years old, and three non-human primate species (rhesus macaque, cynomolgus macaque, African green monkey) (Tables 1, **S1**). The included articles span multiple study protocols, including different target genes, cell lines, exposure conditions, sample types, and sampling times. Only studies using early SARS-CoV-2 strains (i.e., excluding those reporting the D614G mutation or named variants) were included to minimize underlying strain-specific variation. Sampling locations include the upper and lower respiratory tracts, gastrointestinal tract, and other regions.

**Table 1.**
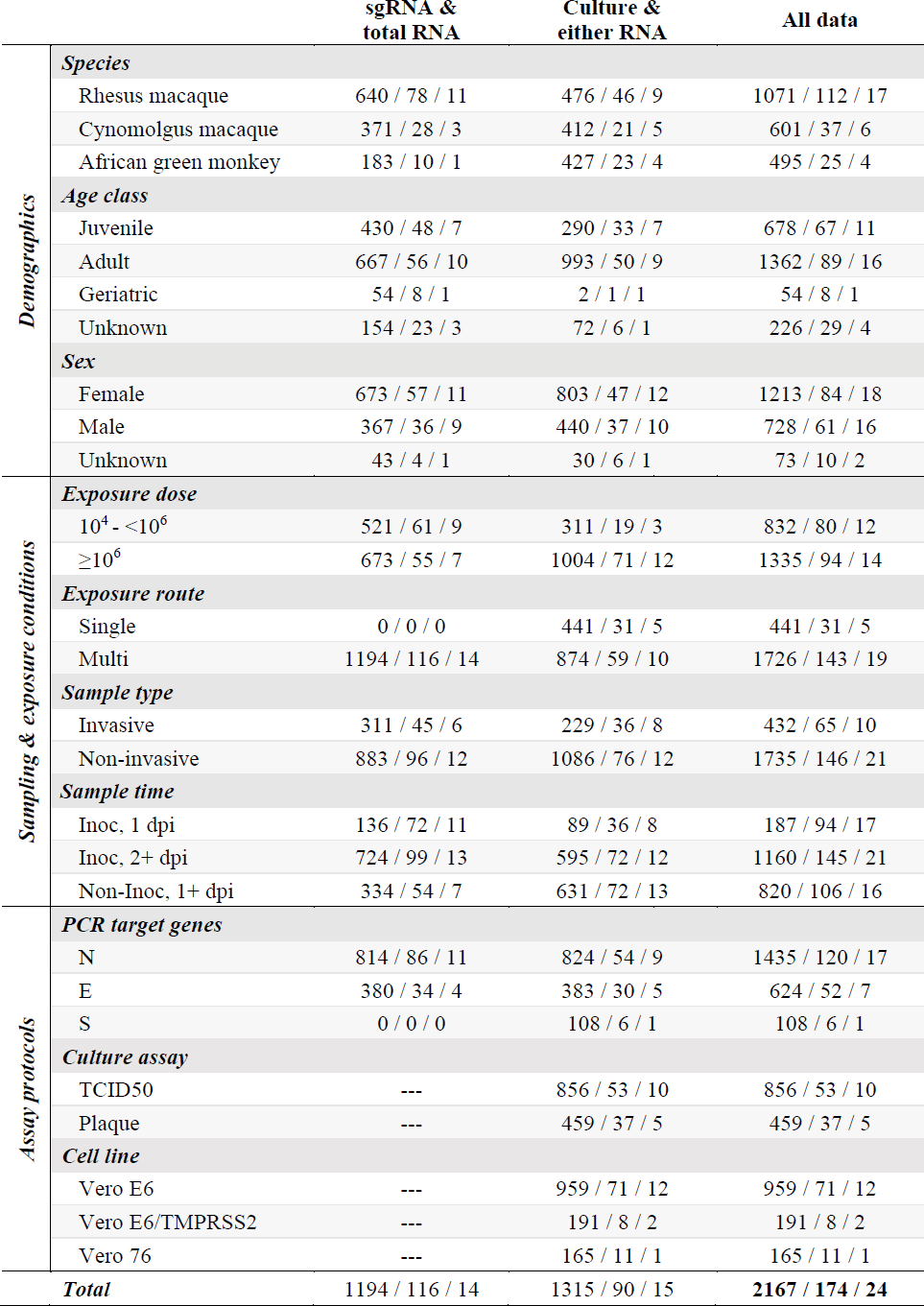
Dataset summary. Columns stratify by assay availability, including samples with results for sgRNA and totRNA, culture and either RNA type, and any combination of two or more included assays. Entries indicate sample sizes for the corresponding cofactor, formatted as: the number of samples/individuals/articles. Doses are grouped by total plaque forming units (though they are analyzed as a continuous variable). Target gene corresponds with the totRNA assay when available, otherwise the sgRNA assay. The full article-specific data distribution is shown in **Table S1**.

### Total RNA quantity does not solely explain sgRNA and culture results

Across individuals and samples in the database, totRNA, sgRNA, and culture trajectories exhibit patterns and challenges consistent with previous reports, including unexpected instances of sgRNA negativity and culture positivity (Figures 1A, **S3-10**). Comparing PCR results, totRNA copy numbers are larger than sgRNA copy numbers when both are detectable (median difference: 1.45 log10 units) (**Figure S11A**), and totRNA becomes undetectable simultaneously or later in infection than sgRNA **(Figure S11D**), with rare exceptions for both patterns likely due to assay noise or processing errors. When both totRNA and sgRNA are detectable for a given individual, their trajectories are typically highly correlated (median Pearson correlation coefficient: 0.92; **Figure S11C**). However, as is particularly common in animal challenge experiments but also reported in clinical data, totRNA-positive samples in this database are often sgRNA-negative (30.0%), and totRNA quantities for these samples can be curiously large, ranging from 0.15 up to 6.38 log10 copy numbers (**Figure S11B**).

**Figure 1.**
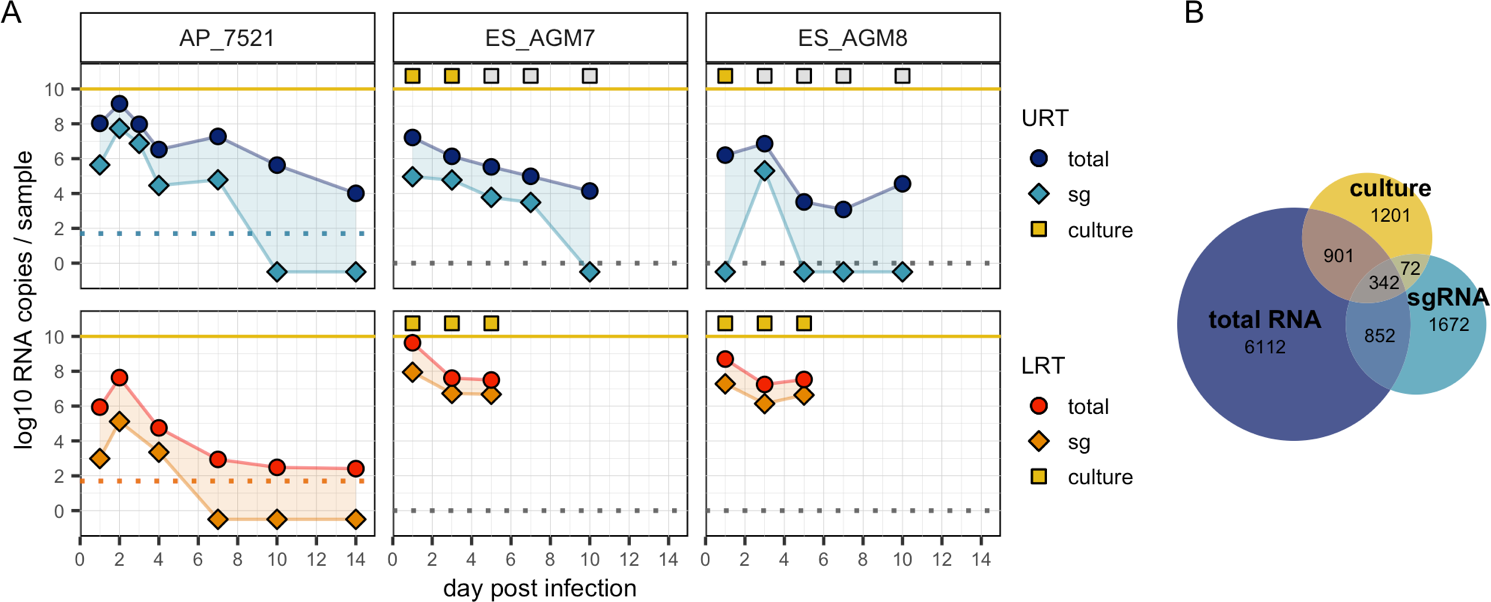
Example trajectories and distribution of samples across assay types. (A) Each column presents the totRNA (circle) and sgRNA (diamond) trajectories for the labelled individual. When available, culture results (square) are plotted above the yellow line, with yellow and grey fill indicating positive or negative culture, respectively. Samples from the upper respiratory tract (URT) are plotted above the lower respiratory tract (LRT). Dashed lines indicate reported limits of detection (plotted at 0 when unavailable). Samples with undetectable RNA are plotted below 0. Representative individuals were chosen from the full dataset. All individual trajectories are shown in **Figures S3-10**. (B) Number of samples available in our database for the corresponding assay(s).

TotRNA and culture positivity results are also often discordant, disagreeing for 39.3% of all available samples and 61.3% of all totRNA-positive samples. Up to 11.02 log10 totRNA copy numbers were quantified in a culture-negative sample, which is only 1 log10 smaller than the maximum copy numbers observed in a culture-positive sample (12.09 log10) (**Figure S11EF**). As few as 2.06 log10 totRNA copy numbers (when detectable) were noted in a culture-positive sample. As expected, totRNA typically becomes detectable earlier and remains detectable later than infectious virus, although for six individuals culture positivity preceded RNA positivity and one culture-positive individual was never totRNA-positive (**Figure S11GH**). Considerably fewer samples with culture data also had sgRNA results (Figure 1B), so comparisons are limited, but patterns broadly parallel those for totRNA. Together, these patterns highlight that totRNA quantity cannot entirely explain sgRNA and culture outcomes. Statistical models may uncover cofactors underlying the discrepancies among these essential assays.

### Predictive performance on withheld data clearly identifies the best statistical models

To compare disparate assays, we developed a Bayesian hurdle model that predicts whether an assay of interest will fall above the limit of detection (the ‘logistic component’) and, if so, predicts a quantitative value for that assay (the ‘linear component’) (**Figure S2**). We used stepwise forward regression with 10-fold cross-validation to evaluate predictive performance on withheld data for variable numbers of predictors. This allowed us to identify the most parsimonious model with similar or better performance on three key metrics compared to the model containing all predictors (the ‘best’ and ‘full’ models, respectively). To benchmark our analysis against prior work, we also evaluated the ‘simple model’ containing PCR results as the sole predictor.

We first applied this method to predict sgRNA from totRNA assays (the ‘sgRNA model’), for which we considered species, age class, sex, exposure dose, day post infection, PCR target gene, and sample type (invasive vs. non-invasive) as candidate predictors. We then applied the logistic model framework to relate PCR results to culture positivity (the ‘culture model’), including cell line and culture assay as additional candidate predictors (see **Methods** for justifications).

For both model types, the selection procedure clearly identified the best models (Figure 2), where each component included a unique set of predictors. These results were robust to alternate cross-validation procedures, predictor scaling, and prior distributions. Each selected model is generalizable, as shown by comparable prediction accuracy between training and test sets. See the **Supplementary Methods** for further details on model evaluation and selection.

**Figure 2.**
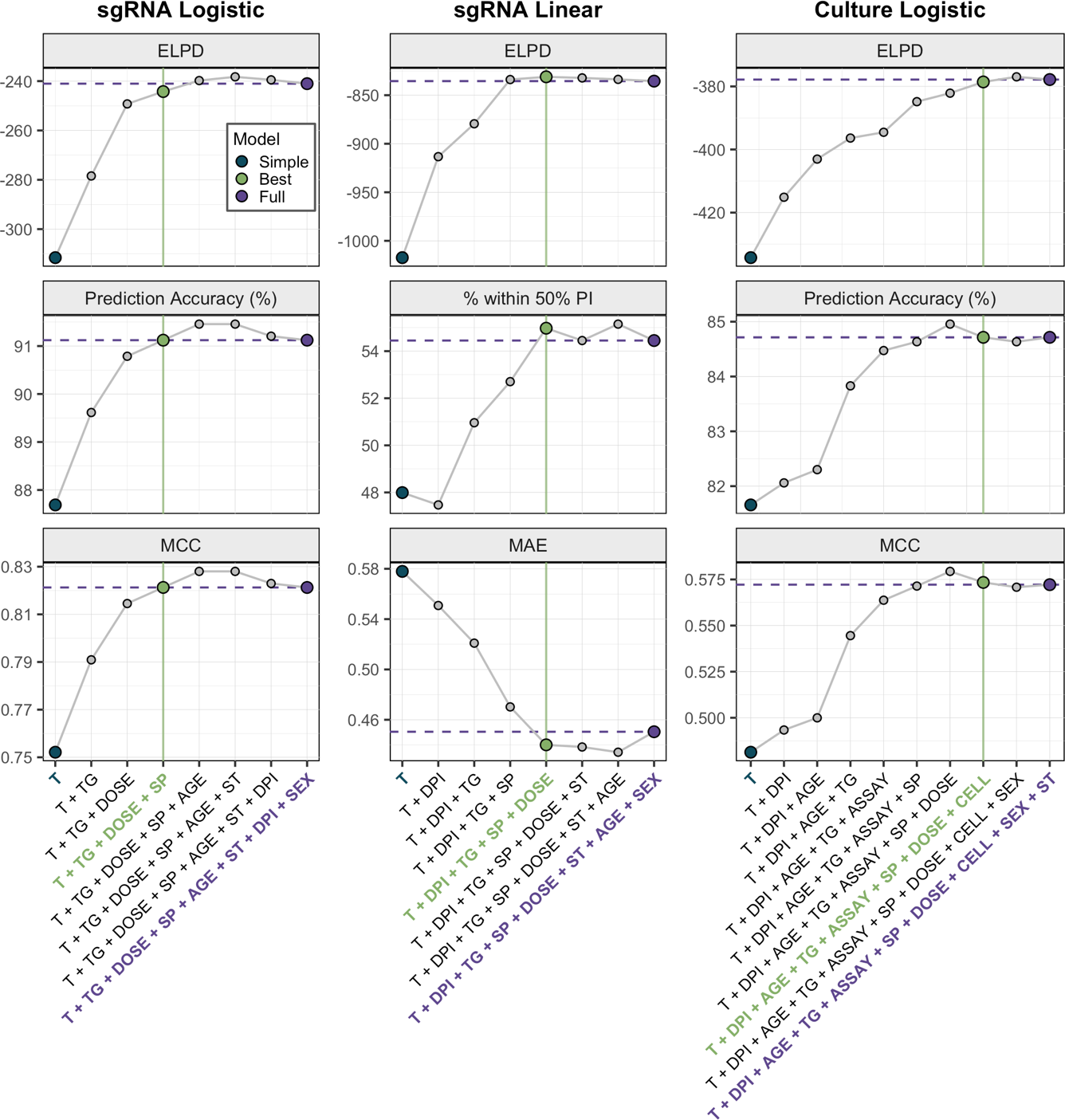
Model selection criteria identify the best models. The highest performing models for each predictor number and modeling component are shown, ordered by increasing predictor numbers. Purple horizontal lines depict performance of the full model. Green vertical lines indicate the best model, chosen according to the displayed metrics. These include estimated log pointwise predictive density (ELPD), prediction accuracy, percent of samples within the 50% prediction interval, Matthews correlation coefficient (MCC), and median absolute error around the median (MAE). These were generated using test data during 10-fold cross validation. Acronyms are: T, totRNA; DPI, day post infection; SP, species; TG, target gene; ST, sample type; CELL, cell line; ASSAY, culture assay. All tested models are shown in **Tables S2-5**.

### Exposure dose, species, and PCR target gene improve predictions of sgRNA positivity

TotRNA levels clearly correlate with sgRNA positivity, but the substantial overlap in totRNA quantities measured for both sgRNA-positive and sgRNA-negative samples emphasize that other factors must influence sgRNA outcomes (Figure 3A). The best sgRNA logistic model identified exposure dose, species, and PCR target gene as key additional predictors of sgRNA positivity (Figure 2, **Table S2**). This model is highly accurate, correctly classifying 91.1% of withheld samples. It outperforms the simple model both by increasing prediction accuracy and by assigning higher probabilities to correct classifications for more samples (Figure 3B). For intermediate quantities of totRNA (2-6 log10 copies), sgRNA positivity predictions differ between the simple and best models (Figure 3C), emphasizing the particular importance of accounting for cofactors in this range. The best and full models perform similarly (Figure 2).

**Figure 3.**
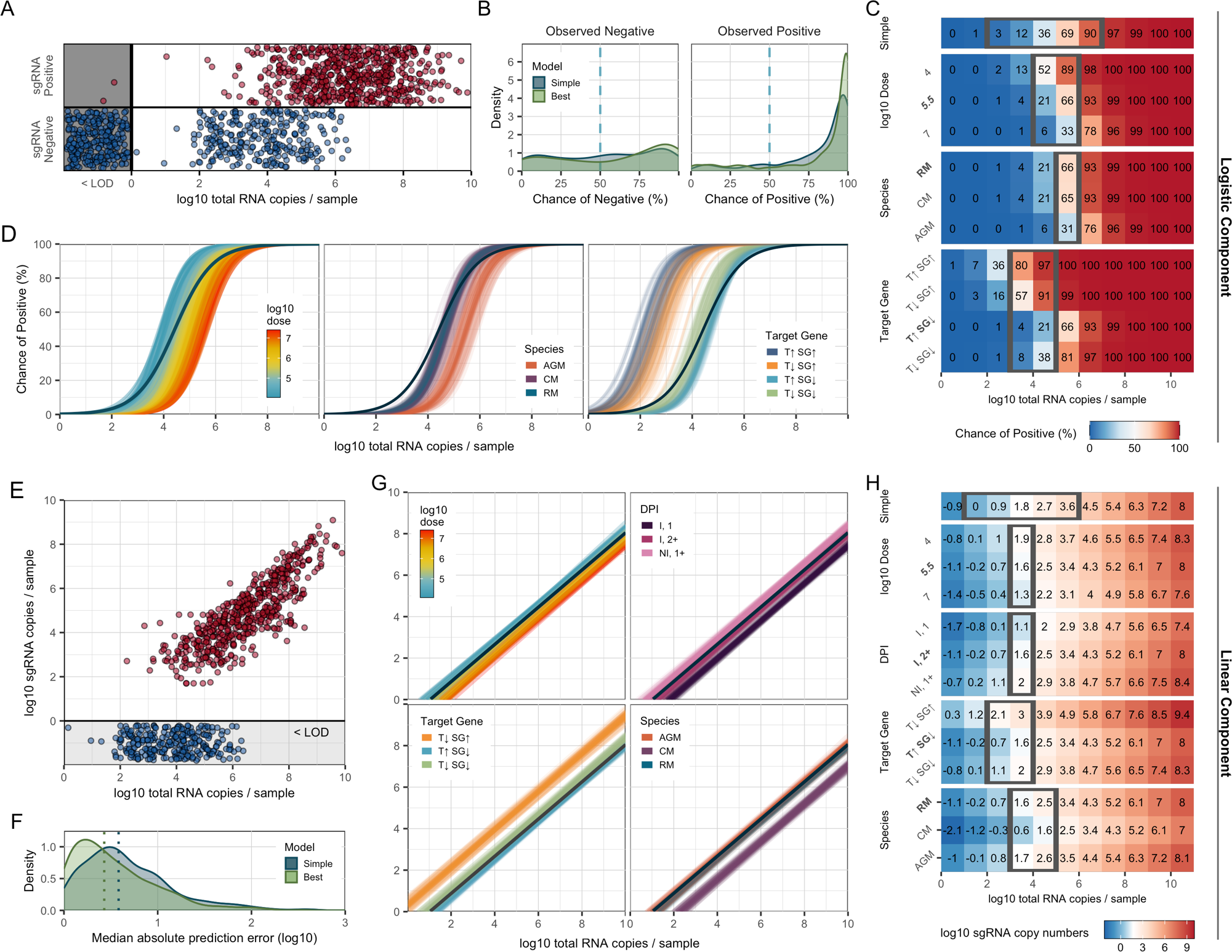
The best sgRNA model captures key sources of underlying variation in PCR outcomes. (A) All available sgRNA data plotted against totRNA results (with vertical jitter), with all totRNA-negative samples plotted in the grey region (with horizontal and vertical jitter). One totRNA- and sgRNA-positive sample with −1.18 log10 totRNA copies is not visible. (B) Distribution of median model-predicted chances of sgRNA detection for all available totRNA-positive samples, stratified by model type and observed outcomes. Samples right of the dashed line are correct predictions. (C) Median predicted chances of sgRNA detection for the simple model (top row) and all cofactor groups for the best model (other rows), evaluated for specific totRNA levels. Predictions were generated using the following ‘standardized cofactor set’ (which are highlighted in bold text): rhesus macaques inoculated with 5.5 log10 pfu and sampled at least two days post infection from inoculated tissues, which were processed with a totRNA-high/sgRNA-low assay. For the simple model, the grey box encloses totRNA copies where classifications differ among the simple model and any possible combination of cofactors. For all other rows, grey boxes enclose regions where classifications differ within the cofactor group for the standardized cofactor set. (D) 300 posterior draws from the best logistic model for the standardized cofactor set, with colored lines as indicated in panel-specific legends. The dark blue line presents the simple model’s mean fit. (E) All available sgRNA data for totRNA-positive samples, where sgRNA-negative samples are plotted below 0 (with vertical jitter). (F) Distribution of median absolute errors for all sgRNA-positive samples, stratified by model type. (G) As in (D) but for the best linear component. (H) As in (C) but reporting median sgRNA copy number predictions. Grey boxes enclose regions where predicted sample quantities within the cofactor group fall both above and below a common limit of detection (1.69 log10). Acronyms are as follows: ‘RM’, rhesus macaque; ‘CM’: cynomolgus macaque; ‘AGM’: African green monkey; ‘Non-Inv’: non-invasive; ‘Inv.’: invasive; ‘DPI’: day post infection; ‘I, 1’: inoculated tissues sampled on day 1 post infection; ‘I, 2+’: inoculated tissues sampled any other day post infection; ‘NI, 1+’; non-inoculated tissues on any day post infection; “T↑SG↑”: totRNA-high/sgRNA-high; “T↓SG↑”: totRNA-low/sgRNA-high; “T↑SG↓”: totRNA-high/sgRNA-low; “T↓SG↓”: totRNA-high/sgRNA-low.

Our best model reveals insights into the three additional predictors of sgRNA outcomes: exposure dose, species, and PCR targe gene. The following trends hold for model predictions across any cofactor combination when holding totRNA quantity constant: (i) individuals inoculated with larger doses have smaller chances of detecting sgRNA, (ii) African green monkeys have the highest chance of sgRNA detection, while rhesus and cynomolgus macaques have similar predictions, and (iii) assays targeting highly-expressed sgRNA species (‘sgRNA-high’ assays) have higher chances of sgRNA detection than those targeting less-expressed sgRNA species (‘sgRNA-low’). We refer the reader to Figure 3C for quantitative median predicted chances of sgRNA detection for a select cofactor combination, Figure 3D for qualitative variability in those predictions, and **Table S6** for the associated 90% prediction intervals. Columns in Figure 3C with a strong color gradient indicate substantial impacts of the associated cofactor on sgRNA predictions, and grey boxes highlight totRNA ranges where final classifications differ within that cofactor group (for the standardized cofactor set).

### Exposure conditions, species, and PCR target gene impact expected RNA ratios

sgRNA quantities scale positively with totRNA quantities, but with considerable unexplained variation (Figure 3E). Our best sgRNA linear model identified exposure dose, species, PCR target gene, and day post infection as key predictors of sgRNA quantity (note these are the same predictors as for the sgRNA logistic model, but with day post infection also included). This model performs well on withheld data, with 55.0% of observed sample values falling within the model-generated 50% prediction interval (Figure 2**, Table S3**). The best model clearly outperforms the simple model, decreasing the median absolute prediction error from 0.58 to 0.43 log10 copies (Figure 3F) and increasing the correlation between observed and median predicted values (from an adjusted R^2^ of 0.68 to 0.77). The best model performs marginally better than the full model, with small improvements in prediction accuracy (Figure 2).

Below, we explore the effects of each selected cofactor on predicted sgRNA copy numbers. We report qualitative trends that hold across all cofactor combinations, and we refer the reader to Figure 3H for median (quantitative) predicted sgRNA copy numbers for a select cofactor combination (our ‘standardized cofactor set’, see figure legend). Variability in these predictions are presented qualitatively in Figure 3G and quantitatively (as 90% prediction intervals) in **Table S6.**

The best model predicts that exposure conditions and sampling time impact RNA ratios. Samples obtained from individuals inoculated with larger doses must have higher total RNA copy numbers to expect the same sgRNA quantity. Results for day post infection parallel these exposure-dependent patterns. To expect a given sgRNA quantity, totRNA copies must be highest for inoculated tissues on the first day post infection, intermediate for inoculated tissues on all later days post infection, and lowest for non-inoculated tissues on any day post infection.

PCR target genes and species also affect predictions. Conditional on totRNA quantity, totRNA-low/sgRNA-high assays have the largest predicted median sgRNA quantities, followed by totRNA-low/sgRNA-low and totRNA-high/sgRNA-low assays. Quantitative sgRNA outcomes were unavailable for totRNA-high/sgRNA-high assays, so estimates were not possible for those protocols. Predictions also varied by species, where African green monkeys and cynomolgus macaques have the highest and lowest predicted median sgRNA quantities for any given totRNA quantity, respectively. Rhesus macaques and African green monkeys had highly similar predictions.

### The sgRNA model accurately reconstructs individual viral load trajectories

To further analyze performance, we reconstructed individual viral load trajectories using the best sgRNA model (Figures 4**, S3-5**). The model correctly predicted the timing of the first and last observed sgRNA positive for 90.1% (n=219/243) and 72.8% (n=177/243) of all individual- and (non-invasive) sample-specific trajectories with at least two sampling times, respectively (**Figure S13**). Notably, 70.0% (n=170/243) of those trajectories were predicted without a single misclassification. The distribution of predicted sgRNA quantities was highly similar to the distribution of observed sgRNA quantities (median differences of estimated means: −0.04 log10 units; 90% Credible Interval [CI]: −0.18, 0.08; **Supplementary Methods**) but highly dissimilar to observed totRNA values (−0.79; 90%CrI: −0.92, −0.66), offering further confidence in the model’s excellent performance.

**Figure 4.**
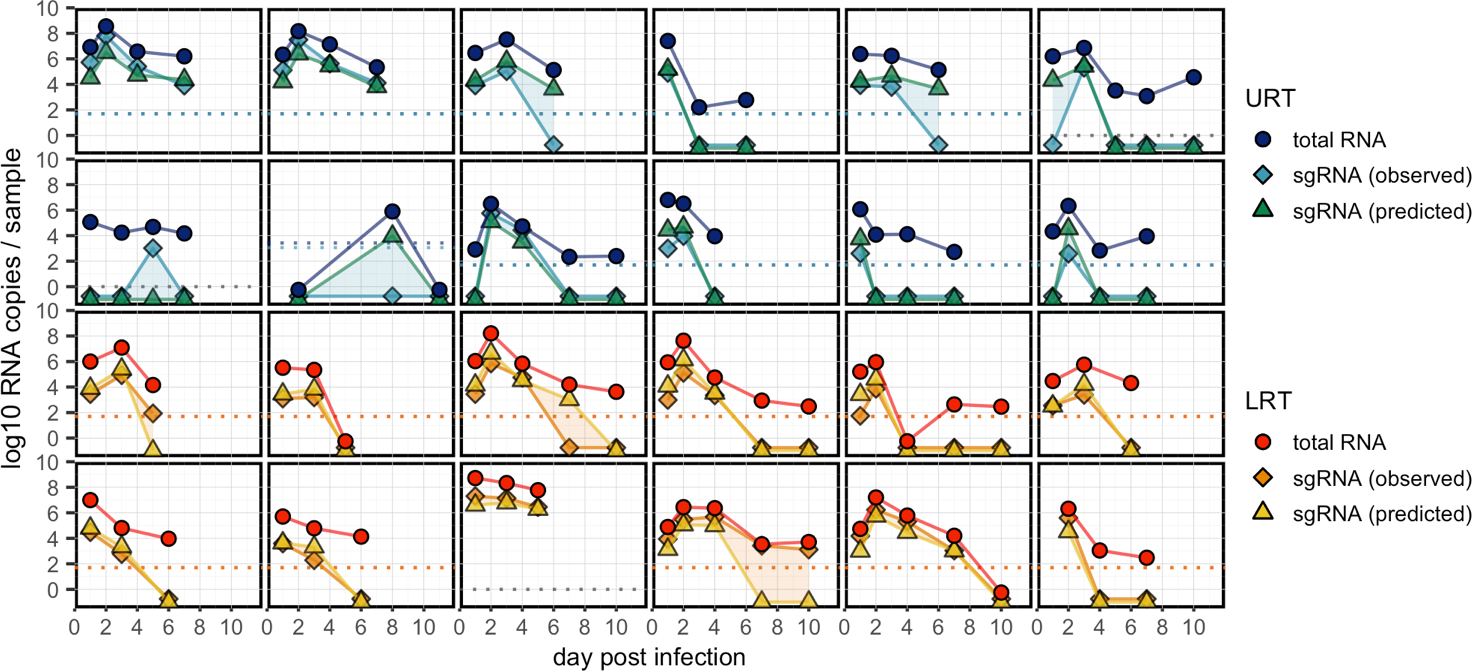
The best sgRNA model reconstructs individual trajectories with high accuracy. Each panel includes the data for one randomly selected individual sampled from either the upper respiratory (URT) or lower respiratory tract (LRT), including observed totRNA (circle), observed sgRNA (diamond), and median predicted sgRNA (triangle) quantities. Detection limits are plotted as dashed lines in the corresponding color when available, otherwise grey lines are plotted at zero. All undetectable samples are plotted below zero. See **Figures S3-5** for all individuals.

### Total RNA and sgRNA are both suitable predictors of viral culture

To determine which PCR assay best predicts viral culture, we compared models including totRNA, sgRNA, or both as predictors of culture positivity. We first evaluated performance only on data with quantitative results for both assays and for models with no additional cofactors, for which totRNA, sgRNA, and both had similar prediction accuracy (**Table S7**). Because few samples had both sgRNA and culture outcomes (Figure 1B), we imputed median sgRNA predictions where needed, using the best performing sgRNA model. On this full dataset, all three models also performed similarly well, though totRNA showed some evidence of better predicting culture positive samples. We then ran our model selection procedure on totRNA and sgRNA separately for all available data, which resulted in highly similar prediction accuracy for both best models, though the model using totRNA was more parsimonious, with two fewer predictors (**Tables S4, S5**). Given this parsimony and the lack of reliance on imputed sgRNA values, plus the lack of evidence that sgRNA is a superior predictor, we based further analyses solely on totRNA.

### Demography, exposure conditions, and assay protocols resolve disparities in culture results

We next sought to predict culture positivity from totRNA results using the logistic model framework. The best model contained day post infection, inoculation dose, age class, species, culture assay, cell line, and PCR target gene as predictors, and it correctly classifies 84.7% of withheld data (Figure 2; **Tables S4, S7**). It outperforms the simple model by correctly predicting an additional 7.0% of culture positive samples and by assigning higher probabilities for true classifications (Figure 5B). Strikingly, culture predictions can differ between the simple and best models for all considered quantities of totRNA (0-12 log10 copies) (Figure 5C), highlighting the necessity of cofactor-mediated inference for culture outcomes. The best model performs similarly to the full model (Figure 2).

**Figure 5.**
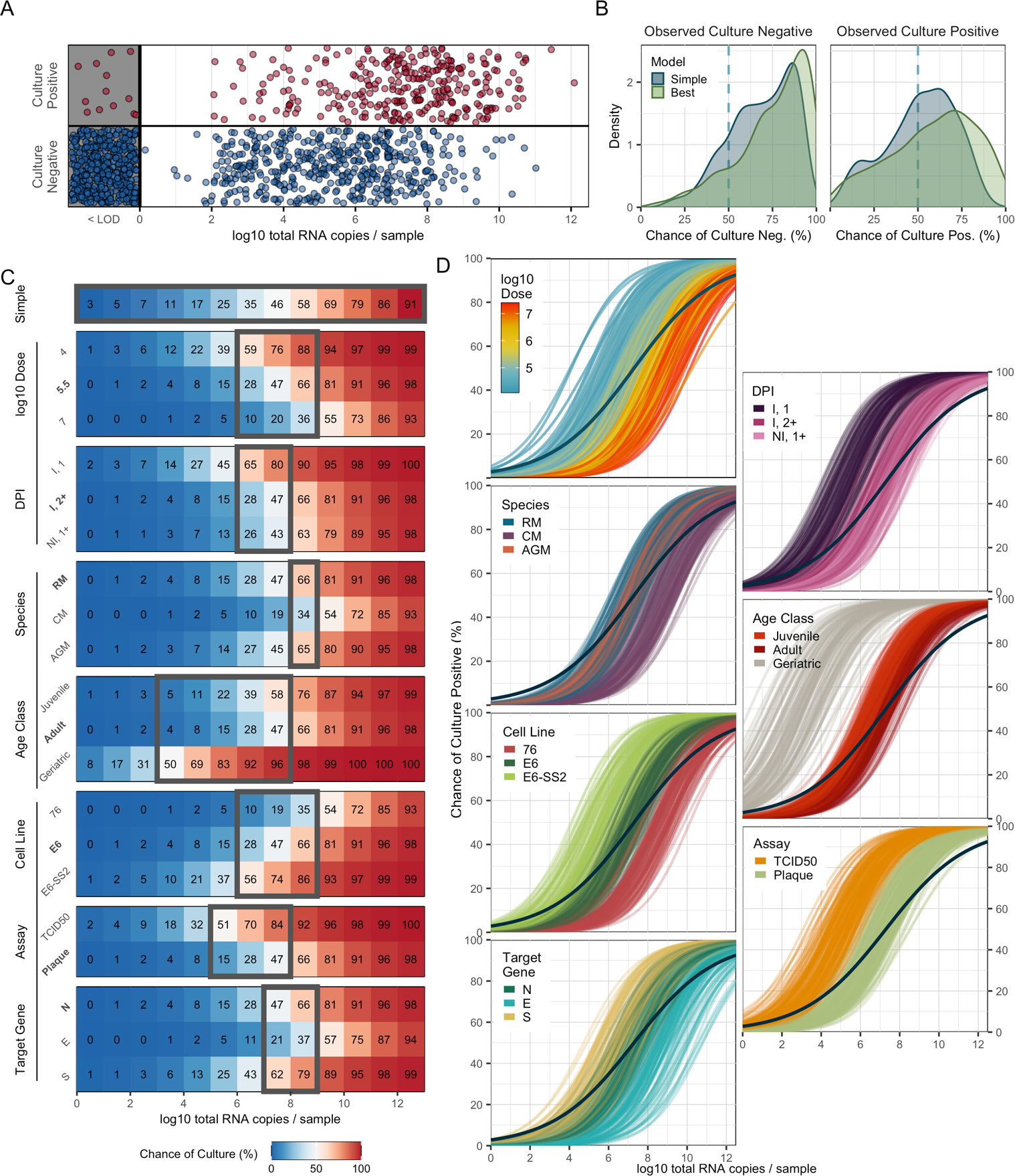
The best culture model captures key sources of underlying variation in culture outcomes. (A) All available culture data plotted against totRNA results (with vertical jitter), with all totRNA-negative samples plotted in the grey region (with horizontal and vertical jitter). (B) Distribution of median model-predicted chances of positive culture for all totRNA-positive samples, stratified by model type and observed outcomes. Samples right of the dashed vertical line are correct predictions. (C) Median predicted chance of positive culture for the simple model (top row) and all cofactor groups included in the best model (other rows) for totRNA copies (evaluated at integer values, starting at 0). Predictions were generated using the following ‘standardized cofactor set’ (which are highlighted in bold text): adult rhesus macaques inoculated with 5.5 log10 pfu and sampled at least two days post infection from inoculated tissues, where PCR targets the Nucleocapsid gene and culture uses plaque assays with VeroE6 cells. Grey boxes enclose regions where classifications differ within the cofactor group for the standardized cofactor set. For the simple model, it encloses regions where classifications differ between the simple model and any possible combination of cofactors. (D) 300 posterior draws from the best model for the standardized cofactor set, with colored lines as indicated in panel-specific legends. The dark blue line presents the simple model’s mean fit. Acronyms are as described in Figure 3, plus the following: E6, VeroE6; E6-SS2, VeroE6-TMPRSS2; and 76, Vero76 cells.

In the text below, we explore the effects of each selected cofactor on culture outcomes. Given the high dimensionality of these predictions, we report qualitative trends that hold for any cofactor combination, and we refer the reader to Figure 5C for median predicted chances of positive culture for a select combination of cofactors (i.e., our ‘standardized cofactor set’, see figure legend). Columns in Figure 5C with a strong color gradient indicate dramatic impacts of the associated cofactor on culture predictions, and grey boxes highlight totRNA ranges where final classifications differ within that cofactor group (for the standardized cofactor set). These ranges differ for other cofactor combinations. We present the variability for our select results qualitatively in Figure 5D and quantitatively (as 90% prediction intervals) in **Table S8**.

Exposure conditions had substantial impacts on culture predictions. Individuals inoculated with larger doses have smaller probabilities of obtaining successful culture for given totRNA quantity. Interestingly, in contrast with results predicting lower sgRNA (per totRNA quantity) in inoculated tissues (Figure 3GH), the culture model predicts that inoculated tissues sampled on the first day post infection have the highest probabilities of being culture positive per totRNA quantity. Inoculated tissues on later days post infection and all non-inoculated tissues are much less likely to be culture positive, with substantial overlap in the predicted probabilities of those two groups.

Multiple demographic factors also affect culture outcomes. Predictions for juvenile and adult age classes largely overlap, but geriatric individuals have substantially higher predicted chances of successful culture for the same viral load. However, few samples from geriatric individuals were available (**Table 2**), and so these results should be interpreted cautiously. Predictions also vary based on species: the chances of successful culture for some viral load are smallest for cynomolgus macaques compared to rhesus macaques and African green monkeys, where the latter two species have highly similar predictions.

Assay conditions also influence culture outcomes, as expected. The model predicts that VeroE6-TMPRSS2 cells have the highest chance of positive culture, followed by VeroE6 and Vero76 cells. TCID50 assays are predicted to have higher sensitivity than plaque assays, and the chances of culture positivity (for a given viral load) are higher for PCR protocols targeting Spike (S) than for those targeting the Nucleocapsid (N) or Envelope (E) genes.

### Individual trajectories uncover sources of culture prediction errors

Although our culture model exhibits remarkable 84.7% accuracy on withheld data, we analyzed our predictions further to identify potential causes and implications of existing errors. 64.1% (n=116/181) of incorrect predictions were false negatives, of which a curious 11.2% (n=13/116) were PCR negative. Even excluding these totRNA-negative samples, totRNA copies for false negative samples were substantially smaller than for true positives (median difference of estimated population means: −2.83 log10 units; 90%CrI: −3.13, −2.53) but more similar to true negatives (median difference: 0.57; 90%CrI: 0.27, 0.87). These RNA-low but culture-positive samples could be explained by PCR or sample processing issues resulting in the amplification of less RNA (e.g., sample degradation), or by culture contamination. Similarly, totRNA copy numbers for false positive predictions were substantially larger than for true negatives (median difference: 3.05; 90%CrI: 2.74, 3.36) but were similar to true positives (median difference: −0.35, 90%CrI: −0.66, −0.04). Culture insensitivity could explain these RNA-high but culture-negative samples.

We further characterized errors by analyzing performance in the context of individual trajectories for (non-invasive) samples with at least two sampling times (Figure 6**, S7-9, S14**). Overall, the model correctly predicted 58.3% (n=120/206) of these trajectories without a single culture misclassification. We categorized errors into four types: (i) samples obtained on the first or last sampling day (termed an ‘edge’), (ii) samples obtained as culture results transition between positive and negative states (‘transition’), (iii) samples where observed culture results change for one sampling time despite surrounding instances of the opposite classification (‘data blip’), and (iv) samples where culture predictions change for one sampling time despite surrounding instances of the opposite classification (‘prediction blip’). Edge and transition errors are the most common, constituting 38.9% (n=51/131) and 33.6% (n=44/131) of the errors, respectively. Edge errors are difficult to analyze, given limited information from surrounding time points, but transitions may reflect sample quality and assay sensitivity interacting to drive noisy outcomes for samples with intermediate RNA or virion quantities. Data blips are less common (n=19/131; 14.5%), for which all except one are observed culture positives surrounded by culture negatives (leading to false-negative prediction errors) (Figures 6B**, S14A**). Eight of these samples co-occur with increases in totRNA quantities from the previous sampling time, suggesting they may reflect true local replication (e.g., as in rebound cases). The remaining instances accompany decreases in totRNA quantities, where sample contamination could drive spurious culture positivity or PCR processing issues could result in RNA underestimates. Prediction blips are the least common (n=17/131; 13.0%), of which 70.6% (n=12/17) are false negatives that often have lower totRNA quantities than the previous sampling time (Figures 6B**, S14B**). These could be explained by sample quality or PCR processing issues resulting in RNA underestimates, which is particularly plausible for instances where totRNA levels increase in the next sampling time. In contrast, false positive prediction blips primarily occur after sharp increases and high quantities of totRNA, and all occur for plaque assays. Given our model predicts lower sensitivity for plaque assays, these errors could reflect failed culture, though RNA overestimates could also explain this pattern.

**Figure 6.**
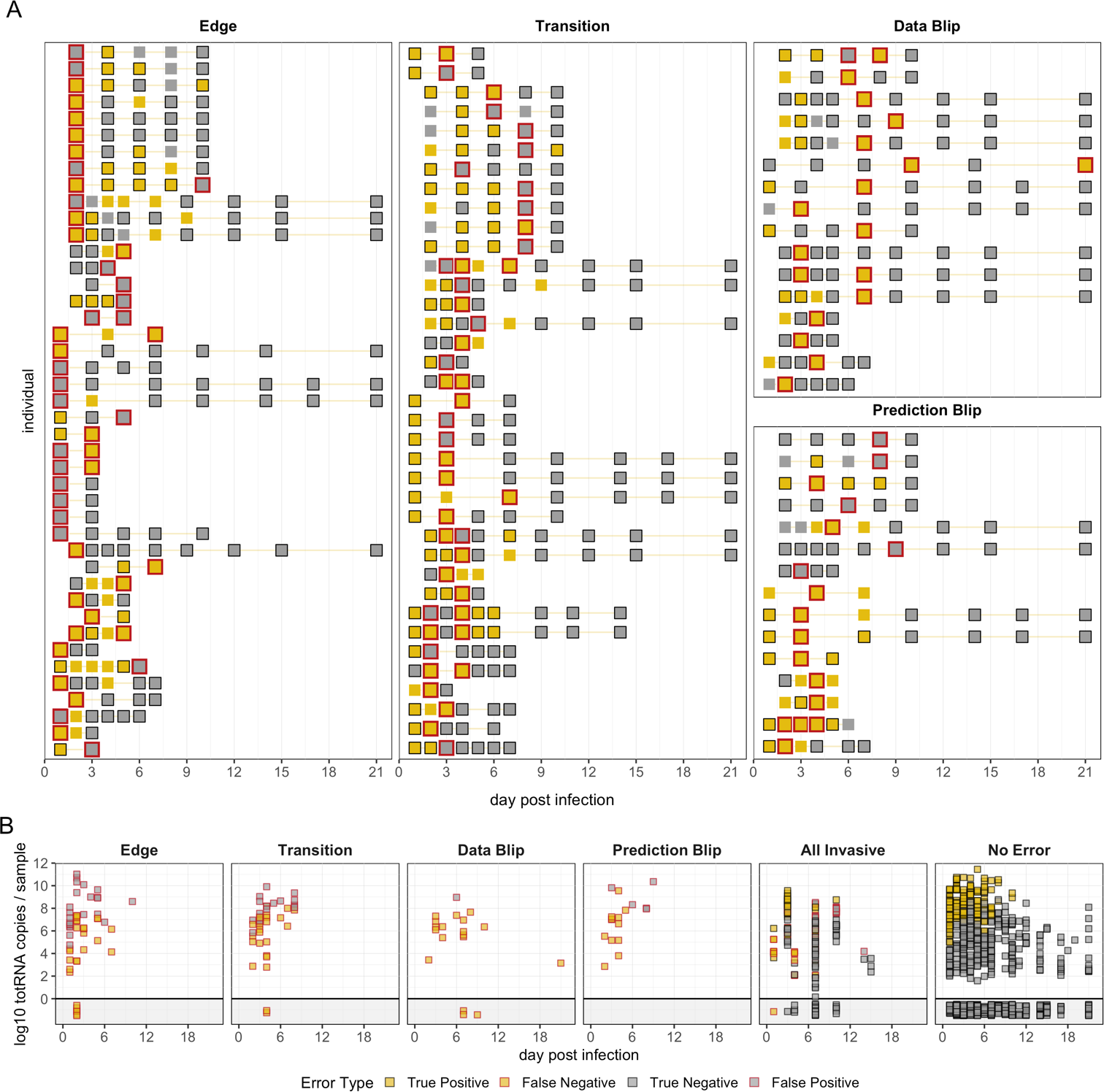
Error analysis reveals potential causes of culture prediction errors. (A) Each row shows culture results for one individual-sample trajectory that contains at least one instance of the panel-specific error type. Trajectories may appear in multiple panels if they contain multiple error types, though trajectory ordering is inconsistent. Red outlines highlight samples with the denoted error type. (B) TotRNA values over time for each error type, all invasive samples, and all correctly classified non-invasive samples (‘no error’). In (A) and (B), yellow squares indicate culture positives and grey indicates culture negatives. Squares with black outline are correctly classified, while those with no or red outline are incorrectly classified. The data blip individual on day 21 post infection has another sample available at a later timepoint, so it is not considered an ‘edge.’

## Discussion

In this study, we developed a generalizable model to infer the results of one virological assay from another. By applying this framework to our compiled database of non-human primate experiments on SARS-CoV-2, we generated highly accurate predictions of sgRNA and culture results from standard PCR protocols. These analyses allowed us to answer foundational questions about whether totRNA and sgRNA assays are fundamentally interchangeable and what factors drive the complicated relationships between PCR and culture outcomes. Our best models identify key sources of biological variation (including exposure conditions, demographics, and assay protocols), across which predictions varied widely. We showed that because standard, simple regression models ignore this variation, they could incorrectly infer culture outcomes for samples with totRNA copy numbers spanning twelve orders of magnitude; our biologically-informed models showed substantial gains in accuracy and precision. Our findings highlight the importance of cofactor-mediated interpretations of viral load and culture positivity – no single threshold value applies across study designs.

We addressed the unresolved debate about the relative merit of sgRNA to predict culture outcomes by conducting the first comprehensive analysis of a large dataset of controlled exposures. We found no clear evidence that sgRNA outperforms totRNA, and instead found that both infer culture outcomes with high accuracy when accounting for key biological covariates. Given these results and that we can reconstruct sgRNA trajectories from totRNA outcomes with high accuracy, underlying cofactors may explain previously observed differences in the relative predictive capacity of totRNA and sgRNA.^9, 11^ Future studies could prospectively measure all three assays (ideally with quantitative culture) to confirm and extend our findings, though notably our model achieved a remarkable 85% accuracy in predicting culture outcomes and our error analysis showed that many prediction errors may have arisen from upstream data issues (see below).

Our models characterize many biological patterns hypothesized (or known) based on previous experimental work, including the effects of exposure conditions on sgRNA and culture outcomes. In particular, we find that larger exposure doses increase the totRNA copy numbers associated with predicting culture positivity and detectable sgRNA. This suggests that the amplification of residual (inoculum-derived) genomic RNA may explain curious instances of sgRNA- or culture-negative samples with large totRNA copies, substantiating concerns in the animal challenge literature that inoculation procedures likely directly influence viral detection and quantification.^7^ The amplification of residual inoculum may also explain differences predicted between inoculated and non-inoculated tissues, where exposed tissues tend to have larger totRNA quantities than non-exposed tissues for any given sgRNA value, particularly on the first day post infection. Inoculum effects on totRNA quantity likely linger throughout infection, given that sgRNA predictions for exposed tissues on later days post infection fall between predictions for exposed tissues on the first day and non-exposed tissues on all days. Interestingly, the chance of positive culture (for a given totRNA value) is highest for exposed tissues sampled on the first day post infection, which is consistent with detection of lingering inoculum-derived virions. In contrast to sgRNA, culture predictions for exposed tissues on all later days post infection are highly similar to non-exposed tissues. These patterns are consistent with most inoculated virions having infected cells, dispersed to other tissues, or been cleared by the immune system within the first two days of infection, whereas the high stability of RNA^62^ likely enables its prolonged detection.

Our work showed that the relationships between virological assays were also shaped by host demographic factors. Primate species affected all relationships we considered, where cynomolgus macaques were predicted to have the lowest sgRNA:totRNA ratio and the smallest chance of positive culture per totRNA quantity. African green monkeys and rhesus macaques have highly similar predictions for sgRNA:totRNA ratios and chances of positive culture. Curiously, African green monkeys also have the smallest chance of sgRNA detection per totRNA quantity, but only one study^8^ reported totRNA and sgRNA outcomes for this species. Our models did not identify age-mediated effects on sgRNA outcomes but did predict geriatrics have the highest chances of positive culture per totRNA quantity. Sex did not influence either sgRNA or culture outcomes. While these results may reflect differing susceptibility, disease severity, or infection kinetics among non-human primate species and age classes, as has been previously suggested,^24, 38, 53, 54, 63^ sample sizes were limited for African green monkeys and geriatrics, so these patterns should be interpreted cautiously. Also, given the complexity of viral fitness, cellular processes, and immune responses, inference on the cause of demographic-specific differences is difficult without mechanistic theory. Mathematical models of the cellular life cycle^64^ may uncover processes that explain the stoichiometric differences we observed among RNA types and virions.

Assay protocols had clear impacts on model predictions. PCR target gene was a consistent factor in our best models, with effects aligned with known differences in RNA quantities. We find that totRNA protocols targeting the Spike (S) gene must amplify less totRNA than those targeting the Envelope (E) or Nucleocapsid (N) genes to predict the same chance of positive culture. This likely reflects that totRNA assays targeting S will amplify only sgS and no other sgRNA species (because it is the most upstream sgRNA), whereas the others amplify multiple sgRNA species and thus will have inherently higher per-sample totRNA copy numbers. Similar reasoning can explain observed differences in sgRNA outcomes, where sgRNA protocols amplifying the highly-expressed sgN have higher chances of detecting sgRNA (per totRNA quantity) and also larger sgRNA:totRNA ratios than protocols amplifying the less-expressed sgE and sg7 species. For viral culture, our model predicts VeroE6-TMPRSS2 cells have the highest chance of detecting infectious virus (per totRNA quantity), which is concordant with the importance of TMPRSS2 for SARS-CoV-2 cellular entry^57^ and agrees with experiments showing VeroE6-TMPRSS2 cells are more permissive to infection than VeroE6 cells.^19, 56^ In accordance with our results, prior work has also shown that VeroE6 cells are more sensitive than Vero76 cells, which is likely related to increased TMPRSS2 expression in VeroE6 cells.^65^ Our model also predicts that TCID50 assays are more likely to detect infectious SARS-CoV-2 than plaque assays, agreeing with standard assay conversions^66^ and prior experimental work.^58^

By analyzing our culture predictions for individual trajectories, we identified potential causes of prediction errors. Many occurred during transition periods when viral replication slows or begins; during these periods, assay readouts will depend strongly on sample quality and assay sensitivity, so additional caution in interpreting culture outcomes is warranted. Beyond this, while we expect some errors due to complex and non-stationary biological effects, many errors are also consistent with PCR or culture processing issues. Sample quality, preservation methods, and storage conditions can substantially impact the quantification of RNA copy numbers and the detection of infectious virus.^67, 68^ PCR issues resulting in the amplification of less RNA may explain curious culture-positive samples with low or no detectable RNA (generating false negative predictions), while culture insensitivity may explain some culture-negative samples with especially large RNA quantities (i.e., false positives). Alternatively, sample contamination or sample swapping could cause elevated RNA levels or spurious culture positivity, where the latter is particularly plausible for ‘data blips’ of a single culture positive surrounded by a series of culture negatives, although these could reflect brief, intermittent replication. In any case, if we assume our model predictions were correct for at least some of these suspect samples (or else if we exclude them from accuracy calculations entirely), our culture model’s true accuracy would be higher than 85%.

With this study, we demonstrated the utility and feasibility of meta-analyses and Bayesian statistical techniques for virological studies, which will become increasingly important tools under new data sharing mandates.^69^ Multiple factors enabled us to rigorously analyze our aggregate database: (i) PCR results were reported as RNA copy numbers, which are internally standardized (as opposed to unstandardized Ct values),^67^ (ii) processing techniques and viral concentrations per reported sample volume are consistent within each study, (iii) many articles reported results for multiple cofactors, and (iv) we accounted for any additional between-study variation by including article-level hierarchical error rates when possible. Under typical analytical approaches, our investigations would have required one study to generate the data for all protocols, samples, and demographics of interest, which would be time and resource prohibitive. Crucially, our approach did not require the generation of new data, which is especially important for non-human primate models where ethical principles^70, 71^ and constrained supply^72, 73^ demand principled data reuse whenever possible.

Although the concordances noted between prior work and our results offer confidence in our models’ performance, our study has limitations. Multiple source articles did not report age class or sex, requiring our model fits to marginalize over all possibilities. Consequently, parameter estimates for age and sex may underestimate their effects. This underscores the importance of comprehensive reporting, especially for animal challenge experiments where using previously collected data would increase adherence to the 3R principles.^70^ Also, few articles reported results for both sgRNA and culture, so some of our investigations relied on sgRNA predictions. Prospective data on all three assays and more comprehensive data panels across cofactors would enable deeper exploration of the predictive capacity of totRNA and sgRNA for viral culture. Finally, while some cofactors were not selected for inclusion in our best models, we cannot exclude the possibility that their effects exist but were not evident or were masked by other predictors. Because covariate coverage relied on different studies in different labs, it remains possible that lab or study effects impacted our results despite our use of article-level error terms, or that some covariate effects were absorbed into these terms. Despite these limitations, our analysis (and similar analyses) can help prioritize resource allocation, so future experiments can more easily adopt the gold-standard approach of testing model-based findings in head-to-head comparisons under fixed conditions.

While we developed this model to analyze PCR and viral culture data from non-human primate experiments, the framework can be adapted easily to other animal models or other viral assays. For example, our model could robustly compare the relationships among antigen tests, PCR, and viral culture, which has recently gained interest^9, 10, 74, 75^ and would benefit from the increased sample size and cofactor coverage possible with definitive meta-analytical treatment. The framework also shows particular potential for clinical diagnostics, where it offers a straightforward, standardized pipeline to predict whether an individual is infectious based on PCR results, which is a clear need.^12, 15, 17, 19–22^ With such a tool, public health officials and clinicians could weigh transmission risk with medical resource availability and economic burden to designate evidence-based hospital discharge criteria and public health guidelines. This would initially require careful calibration on high-quality data and some model modifications, including adjusting for infection timing by accounting for days post symptom onset instead of days post infection. Prediction accuracy would likely further increase if the models capitalized on individual-specific viral load trajectories when available (e.g., for hospitalized patients undergoing regular screening) or other proxies like rapid antigen tests. These model adjustments could range in complexity from incorporating a mechanistic modeling component^76^ to simply including an additional predictor for temporal changes in RNA copy numbers.

By assembling and analyzing a large database that captures infection patterns described in the clinical and animal challenge literature, we demonstrated that highly accurate RNA-based culture predictions are possible with our statistical framework. By using non-human primate data, we were able to identify underlying effects of exposure conditions, which would be impossible for humans without experimental challenge trials (of which only one exists for SARS-CoV-2, to date^77^). Consequently, our model offers the first set of explicit quantitative guidelines on interpreting assay outcomes in light of exposure conditions, which has direct implications for analyzing animal challenge experiments. We propose our method as a standardized framework to conduct assay comparisons, whether for individual virology experiments, clinical diagnostic settings, qualitative literature syntheses, or quantitative meta-analyses. Such approaches for data aggregation and (meta-)analysis are vital and powerful tools for an era of increasing data-sharing, with untapped potential to develop translational applications and to guide further research into fundamental mechanisms.

## Data and Code Availability

All data and code used to produce the results and figures in this manuscript (including some additional analyses not included because of space constraints) will be made available on GitHub at: https://github.com/celinesnedden/sars2-assay-comp.

## Acknowledgements

We thank the authors of all articles included in this study, many of whom kindly corresponded with us to provide clarifications and raw data. We also thank the members of the Lloyd-Smith lab for valuable discussions of this work. J.O.L.-S. and C.E.S were both supported by the Defense Advanced Research Projects Agency DARPA PREEMPT #D18AC00031. C.E.S was also supported by the National Institutes of Health (grant 5T32 GM008185-33) and the UCLA Office of the Vice Chancellor for Research 3R Grant. J.O.L.-S. was also supported by the National Science Foundation (DEB-1557022 and DEB-2245631) and the UCLA AIDS Institute and Charity Treks. The content of the article does not necessarily reflect the position or the policy of the US government, and no official endorsement should be inferred.

## Author contributions

Conceptualization, J.O.L.-S and C.E.S; Methodology, J.O.L.-S and C.E.S; Formal Analysis, C.E.S.; Writing – Original Draft, C.E.S.; Writing – Review & Editing, J.O.L.-S and C.E.S; Visualization, C.E.S; Funding Acquisition, J.O.L.-S and C.E.S.

## References

1. Kralik, P., and Ricchi, M. (2017). A Basic Guide to Real Time PCR in Microbial Diagnostics: Definitions, Parameters, and Everything. Front. Microbiol. 8, 108. 10.3389/fmicb.2017.00108.

2. Yang, S., and Rothman, R.E. (2004). PCR-based diagnostics for infectious diseases: uses, limitations, and future applications in acute-care settings. Lancet Infect. Dis. 4, 337–348. 10.1016/S1473-3099(04)01044-8.

3. Fehr, A.R., and Perlman, S. (2015). Coronaviruses: An Overview of Their Replication and Pathogenesis. In Coronaviruses: Methods and Protocols Methods in Molecular Biology., H. J. Maier, E. Bickerton, and P. Britton, eds. (Springer), pp. 1–23. 10.1007/978-1-4939-2438-7_1.

4. Lu, X., Wang, L., Sakthivel, S.K., Whitaker, B., Murray, J., Kamili, S., Lynch, B., Malapati, L., Burke, S.A., Harcourt, J., et al. (2020). US CDC Real-Time Reverse Transcription PCR Panel for Detection of Severe Acute Respiratory Syndrome Coronavirus 2. Emerg Infect Dis. 26, 1654–1665. 10.3201/eid2608.201246.

5. Escors, D., Izeta, A., Capiscol, C., and Enjuanes, L. (2003). Transmissible Gastroenteritis Coronavirus Packaging Signal Is Located at the 5′ End of the Virus Genome. J. Virol. 77, 7890– 7902. 10.1128/JVI.77.14.7890-7902.2003.

6. Wölfel, R., Corman, V.M., Guggemos, W., Seilmaier, M., Zange, S., Müller, M.A., Niemeyer, D., Jones, T.C., Vollmar, P., Rothe, C., et al. (2020). Virological assessment of hospitalized patients with COVID-2019. Nature 581, 465–469. 10.1038/s41586-020-2196-x.

7. Dagotto, G., Mercado, N.B., Martinez, D.R., Hou, Y.J., Nkolola, J.P., Carnahan, R.H., Crowe, J.E., Baric, R.S., and Barouch, D.H. (2021). Comparison of Subgenomic and Total RNA in SARS-CoV-2-Challenged Rhesus Macaques. J. Virol. 95. 10.1128/JVI.02370-20.

8. Speranza, E., Williamson, B.N., Feldmann, F., Sturdevant, G.L., Pérez, L.P.-, Meade-White, K., Smith, B.J., Lovaglio, J., Martens, C., Munster, V.J., et al. (2021). Single-cell RNA sequencing reveals SARS-CoV-2 infection dynamics in lungs of African green monkeys. Sci. Transl. Med. 13. 10.1126/scitranslmed.abe8146.

9. Ford, L., Lee, C., Pray, I.W., Cole, D., Bigouette, J.P., Abedi, G.R., Bushman, D., Delahoy, M.J., Currie, D.W., Cherney, B., et al. (2021). Epidemiologic Characteristics Associated With Severe Acute Respiratory Syndrome Coronavirus 2 (SARS-CoV-2) Antigen-Based Test Results, Real-Time Reverse Transcription Polymerase Chain Reaction (rRT-PCR) Cycle Threshold Values, Subgenomic RNA, and Viral Culture Results From University Testing. Clin. Infect. Dis. 73, e1348–e1355. 10.1093/cid/ciab303.

10. Bonenfant, G., Deyoe, J.E., Wong, T., Grijalva, C.G., Cui, D., Talbot, H.K., Hassell, N., Halasa, N., Chappell, J., Thornburg, N.J., et al. (2022). Surveillance and Correlation of Severe Acute Respiratory Syndrome Coronavirus 2 Viral RNA, Antigen, Virus Isolation, and Self-Reported Symptoms in a Longitudinal Study With Daily Sampling. Clin. Infect. Dis. 75, 1698–1705. 10.1093/cid/ciac282.

11. Perera, R.A.P.M., Tso, E., Tsang, O.T.Y., Tsang, D.N.C., Fung, K., Leung, Y.W.Y., Chin, A.W.H., Chu, D.K.W., Cheng, S.M.S., Poon, L.L.M., et al. (2020). SARS-CoV-2 Virus Culture and Subgenomic RNA for Respiratory Specimens from Patients with Mild Coronavirus Disease - Volume 26, Number 11—November 2020 - Emerging Infectious Diseases journal - CDC. 26, 2701–2704. 10.3201/eid2611.203219.

12. Bravo, M.S., Berengua, C., Marín, P., Esteban, M., Rodriguez, C., del Cuerpo, M., Miró, E., Cuesta, G., Mosquera, M., Sánchez-Palomino, S., et al. (2022). Viral Culture Confirmed SARS-CoV-2 Subgenomic RNA Value as a Good Surrogate Marker of Infectivity. J. Clin. Microbiol. 60, e01609–21. 10.1128/JCM.01609-21.

13. Rodríguez-Grande, C., Adán-Jiménez, J., Catalán, P., Alcalá, L., Estévez, A., Muñoz, P., Pérez-Lago, L., García de Viedma, D., and on behalf of the Gregorio Marañón Microbiology-ID COVID-19 Study Group (2021). Inference of Active Viral Replication in Cases with Sustained Positive Reverse Transcription-PCR Results for SARS-CoV-2. J. Clin. Microbiol. 59, e02277-20. 10.1128/JCM.02277-20.

14. Alexandersen, S., Chamings, A., and Bhatta, T.R. (2020). SARS-CoV-2 genomic and subgenomic RNAs in diagnostic samples are not an indicator of active replication. Nat. Commun. 11, 6059. 10.1038/s41467-020-19883-7.

15. Dimcheff, D.E., Valesano, A.L., Rumfelt, K.E., Fitzsimmons, W.J., Blair, C., Mirabelli, C., Petrie, J.G., Martin, E.T., Bhambhani, C., Tewari, M., et al. (2021). Severe Acute Respiratory Syndrome Coronavirus 2 Total and Subgenomic RNA Viral Load in Hospitalized Patients. J. Infect. Dis. 224, 1287–1293. 10.1093/infdis/jiab215.

16. Verma, R., Kim, E., Martínez-Colón, G.J., Jagannathan, P., Rustagi, A., Parsonnet, J., Bonilla, H., Khosla, C., Holubar, M., Subramanian, A., et al. (2021). SARS-CoV-2 Subgenomic RNA Kinetics in Longitudinal Clinical Samples. Open Forum Infect. Dis. 8, ofab310. 10.1093/ofid/ofab310.

17. van Kampen, J.J.A., van de Vijver, D.A.M.C., Fraaij, P.L.A., Haagmans, B.L., Lamers, M.M., Okba, N., van den Akker, J.P.C., Endeman, H., Gommers, D.A.M.P.J., Cornelissen, J.J., et al. (2021). Duration and key determinants of infectious virus shedding in hospitalized patients with coronavirus disease-2019 (COVID-19). Nat. Commun. 12, 267. 10.1038/s41467-020-20568-4.

18. Salguero, F.J., White, A.D., Slack, G.S., Fotheringham, S.A., Bewley, K.R., Gooch, K.E., Longet, S., Humphries, H.E., Watson, R.J., Hunter, L., et al. (2021). Comparison of rhesus and cynomolgus macaques as an infection model for COVID-19. Nat. Commun. 12, 1260. 10.1038/s41467-021-21389-9.

19. Bruce, E.A., Mills, M.G., Sampoleo, R., Perchetti, G.A., Huang, M.-L., Despres, H.W., Schmidt, M.M., Roychoudhury, P., Shirley, D.J., Jerome, K.R., et al. (2022). Predicting infectivity: comparing four PCR-based assays to detect culturable SARS-CoV-2 in clinical samples. EMBO Mol. Med. 14, e15290. 10.15252/emmm.202115290.

20. Bullard, J., Dust, K., Funk, D., Strong, J.E., Alexander, D., Garnett, L., Boodman, C., Bello, A., Hedley, A., Schiffman, Z., et al. (2020). Predicting Infectious Severe Acute Respiratory Syndrome Coronavirus 2 From Diagnostic Samples. Clin. Infect. Dis. 71, 2663–2666. 10.1093/cid/ciaa638.

21. Gniazdowski, V., Paul Morris, C., Wohl, S., Mehoke, T., Ramakrishnan, S., Thielen, P., Powell, H., Smith, B., Armstrong, D.T., Herrera, M., et al. (2021). Repeated Coronavirus Disease 2019 Molecular Testing: Correlation of Severe Acute Respiratory Syndrome Coronavirus 2 Culture With Molecular Assays and Cycle Thresholds. Clin. Infect. Dis. 73, e860–e869. 10.1093/cid/ciaa1616.

22. La Scola, B., Le Bideau, M., Andreani, J., Hoang, V.T., Grimaldier, C., Colson, P., Gautret, P., and Raoult, D. (2020). Viral RNA load as determined by cell culture as a management tool for discharge of SARS-CoV-2 patients from infectious disease wards. Eur. J. Clin. Microbiol. Infect. Dis. 39, 1059–1061. 10.1007/s10096-020-03913-9.

23. Gadi, N., Wu, S.C., Spihlman, A.P., and Moulton, V.R. (2020). What’s Sex Got to Do With COVID-19? Gender-Based Differences in the Host Immune Response to Coronaviruses. Front. Immunol. 11.

24. Bajaj, V., Gadi, N., Spihlman, A.P., Wu, S.C., Choi, C.H., and Moulton, V.R. (2021). Aging, Immunity, and COVID-19: How Age Influences the Host Immune Response to Coronavirus Infections? Front. Physiol. 11.

25. Fajnzylber, J., Regan, J., Coxen, K., Corry, H., Wong, C., Rosenthal, A., Worrall, D., Giguel, F., Piechocka-Trocha, A., Atyeo, C., et al. (2020). SARS-CoV-2 viral load is associated with increased disease severity and mortality. Nat. Commun. 11, 5493. 10.1038/s41467-020-19057-5.

26. Kim, M.-C., Cui, C., Shin, K.-R., Bae, J.-Y., Kweon, O.-J., Lee, M.-K., Choi, S.-H., Jung, S.-Y., Park, M.-S., and Chung, J.-W. (2021). Duration of Culturable SARS-CoV-2 in Hospitalized Patients with Covid-19. N. Engl. J. Med. 384, 671–673. 10.1056/NEJMc2027040.

27. Salvatore, P.P., Dawson, P., Wadhwa, A., Rabold, E.M., Buono, S., Dietrich, E.A., Reses, H.E., Vuong, J., Pawloski, L., Dasu, T., et al. (2021). Epidemiological Correlates of Polymerase Chain Reaction Cycle Threshold Values in the Detection of Severe Acute Respiratory Syndrome Coronavirus 2 (SARS-CoV-2). Clin. Infect. Dis. 72, e761–e767. 10.1093/cid/ciaa1469.

28. Estes, J.D., Wong, S.W., and Brenchley, J.M. (2018). Nonhuman primate models of human viral infections. Nat. Rev. Immunol. 18, 390–404. 10.1038/s41577-018-0005-7.

29. Moher, D., Liberati, A., Tetzlaff, J., and Altman, D.G. (2009). Preferred Reporting Items for Systematic Reviews and Meta-Analyses: The PRISMA Statement. Ann. Intern. Med. 151, 264–269. 10.7326/0003-4819-151-4-200908180-00135.

30. Baum, A., Ajithdoss, D., Copin, R., Zhou, A., Lanza, K., Negron, N., Ni, M., Wei, Y., Mohammadi, K., Musser, B., et al. (2020). REGN-COV2 antibodies prevent and treat SARS-CoV-2 infection in rhesus macaques and hamsters. Science 370, 1110–1115. 10.1126/science.abe2402.

31. Chandrashekar, A., Liu, J., Martinot, A.J., McMahan, K., Mercado, N.B., Peter, L., Tostanoski, L.H., Yu, J., Maliga, Z., Nekorchuk, M., et al. (2020). SARS-CoV-2 infection protects against rechallenge in rhesus macaques. Science, eabc4776–eabc4776. 10.1126/science.abc4776.

32. Corbett, K.S., Flynn, B., Foulds, K.E., Francica, J.R., Boyoglu-Barnum, S., Werner, A.P., Flach, B., O’Connell, S., Bock, K.W., Minai, M., et al. (2020). Evaluation of the mRNA-1273 Vaccine against SARS-CoV-2 in Nonhuman Primates. N. Engl. J. Med., NEJMoa2024671–NEJMoa2024671. 10.1056/NEJMoa2024671.

33. Cross, R.W., Agans, K.N., Prasad, A.N., Borisevich, V., Woolsey, C., Deer, D.J., Dobias, N.S., Geisbert, J.B., Fenton, K.A., and Geisbert, T.W. (2020). Intranasal exposure of African green monkeys to SARS-CoV-2 results in acute phase pneumonia with shedding and lung injury still present in the early convalescence phase. Virol. J. 17, 125–125. 10.1186/s12985-020-01396-w.

34. Deng, W., Bao, L., Gao, H., Xiang, Z., Qu, Y., Song, Z., Gong, S., Liu, J., Liu, J., Yu, P., et al. (2020). Ocular conjunctival inoculation of SARS-CoV-2 can cause mild COVID-19 in rhesus macaques. Nat. Commun. 11, 4400–4400. 10.1038/s41467-020-18149-6.

35. Gabitzsch, E., Safrit, J.T., Verma, M., Rice, A., Sieling, P., Zakin, L., Shin, A., Morimoto, B., Adisetiyo, H., Wong, R., et al. (2021). Dual-Antigen COVID-19 Vaccine Subcutaneous Prime Delivery With Oral Boosts Protects NHP Against SARS-CoV-2 Challenge. Front. Immunol. 12.

36. Jiao, L., Li, H., Xu, J., Yang, M., Ma, C., Li, J., Zhao, S., Wang, H., Yang, Y., Yu, W., et al. (2021). The Gastrointestinal Tract Is an Alternative Route for SARS-CoV-2 Infection in a Nonhuman Primate Model. Gastroenterology 160, 1647–1661. 10.1053/j.gastro.2020.12.001.

37. Ishigaki, H., Nakayama, M., Kitagawa, Y., Nguyen, C.T., Hayashi, K., Shiohara, M., Gotoh, B., and Itoh, Y. (2021). Neutralizing antibody-dependent and -independent immune responses against SARS-CoV-2 in cynomolgus macaques. Virology 554, 97–105. 10.1016/j.virol.2020.12.013.

38. Johnston, S.C., Ricks, K.M., Jay, A., Raymond, J.L., Rossi, F., Zeng, X., Scruggs, J., Dyer, D., Frick, O., Koehler, J.W., et al. (2021). Development of a coronavirus disease 2019 nonhuman primate model using airborne exposure. PLOS ONE 16, e0246366. 10.1371/journal.pone.0246366.

39. Jones, B.E., Brown-Augsburger, P.L., Corbett, K.S., Westendorf, K., Davies, J., Cujec, T.P., Wiethoff, C.M., Blackbourne, J.L., Heinz, B.A., Foster, D., et al. (2021). The neutralizing antibody, LY-CoV555, protects against SARS-CoV-2 infection in nonhuman primates. Sci. Transl. Med. 13, eabf1906. 10.1126/scitranslmed.abf1906.

40. Kobiyama, K., Imai, M., Jounai, N., Nakayama, M., Hioki, K., Iwatsuki-Horimoto, K., Yamayoshi, S., Tsuchida, J., Niwa, T., Suzuki, T., et al. (2021). Optimization of an LNP-mRNA vaccine candidate targeting SARS-CoV-2 receptor-binding domain. 2021.03.04.433852. 10.1101/2021.03.04.433852.

41. Li, D., Edwards, R.J., Manne, K., Martinez, D.R., Schäfer, A., Alam, S.M., Wiehe, K., Lu, X., Parks, R., Sutherland, L.L., et al. (2021). In vitro and in vivo functions of SARS-CoV-2 infection-enhancing and neutralizing antibodies. Cell 184, 4203–4219.e32. 10.1016/j.cell.2021.06.021.

42. Munster, V.J., Feldmann, F., Williamson, B.N., van Doremalen, N., Pérez-Pérez, L., Schulz, J., Meade-White, K., Okumura, A., Callison, J., Brumbaugh, B., et al. (2020). Respiratory disease in rhesus macaques inoculated with SARS-CoV-2. Nature 585, 268–272. 10.1038/s41586-020-2324-7.

43. Nagata, N., Iwata-Yoshikawa, N., Sano, K., Ainai, A., Shiwa, N., Shirakura, M., Kishida, N., Arita, T., Suzuki, Y., Harada, T., et al. (2021). The peripheral T cell population is associated with pneumonia severity in cynomolgus monkeys experimentally infected with severe acute respiratory syndrome coronavirus 2. 2021.01.07.425698. 10.1101/2021.01.07.425698.

44. Patel, A., Walters, J.N., Reuschel, E.L., Schultheis, K., Parzych, E., Gary, E.N., Maricic, I., Purwar, M., Eblimit, Z., Walker, S.N., et al. (2021). Intradermal-delivered DNA vaccine induces durable immunity mediating a reduction in viral load in a rhesus macaque SARS-CoV-2 challenge model. Cell Rep. Med. 2, 100420. 10.1016/j.xcrm.2021.100420.

45. Shan, C., Yao, Y.-F., Yang, X.-L., Zhou, Y.-W., Gao, G., Peng, Y., Yang, L., Hu, X., Xiong, J., Jiang, R.-D., et al. (2020). Infection with novel coronavirus (SARS-CoV-2) causes pneumonia in Rhesus macaques. Cell Res., 1–8. 10.1038/s41422-020-0364-z.

46. Singh, D.K., Singh, B., Ganatra, S.R., Gazi, M., Cole, J., Thippeshappa, R., Alfson, K.J., Clemmons, E., Gonzalez, O., Escobedo, R., et al. (2021). Responses to acute infection with SARS-CoV-2 in the lungs of rhesus macaques, baboons and marmosets. Nat. Microbiol. 6, 73–86. 10.1038/s41564-020-00841-4.

47. Williamson, B.N., Feldmann, F., Schwarz, B., Meade-White, K., Porter, D.P., Schulz, J., van Doremalen, N., Leighton, I., Yinda, C.K., Pérez-Pérez, L., et al. (2020). Clinical benefit of remdesivir in rhesus macaques infected with SARS-CoV-2. Nature, 1–7. 10.1038/s41586-020-2423-5.

48. Yu, J., Tostanoski, L.H., Peter, L., Mercado, N.B., McMahan, K., Mahrokhian, S.H., Nkolola, J.P., Liu, J., Li, Z., Chandrashekar, A., et al. (2020). DNA vaccine protection against SARS-CoV-2 in rhesus macaques. Science, eabc6284–eabc6284. 10.1126/science.abc6284.

49. Woolsey, C., Borisevich, V., Prasad, A.N., Agans, K.N., Deer, D.J., Dobias, N.S., Heymann, J.C., Foster, S.L., Levine, C.B., Medina, L., et al. (2021). Establishment of an African green monkey model for COVID-19 and protection against re-infection. Nat. Immunol. 22, 86–98. 10.1038/s41590-020-00835-8.

50. van Doremalen, N., Lambe, T., Spencer, A., Belij-Rammerstorfer, S., Purushotham, J.N., Port, J.R., Avanzato, V.A., Bushmaker, T., Flaxman, A., Ulaszewska, M., et al. (2020). ChAdOx1 nCoV-19 vaccine prevents SARS-CoV-2 pneumonia in rhesus macaques. Nature, 1–8. 10.1038/s41586-020-2608-y.

51. Poisot, T. (2011). The digitize package: extracting numerical data from scatterplots.

52. R Core Team (2022). R: A Language and Environment for Statistical Computing.

53. Lu, S., Zhao, Y., Yu, W., Yang, Y., Gao, J., Wang, J., Kuang, D., Yang, M., Yang, J., Ma, C., et al. (2020). Comparison of nonhuman primates identified the suitable model for COVID-19. Signal Transduct. Target. Ther. 5, 157–157. 10.1038/s41392-020-00269-6.

54. Blair, R.V., Vaccari, M., Doyle-Meyers, L.A., Roy, C.J., Russell-Lodrigue, K., Fahlberg, M., Monjure, C.J., Beddingfield, B., Plante, K.S., Plante, J.A., et al. (2021). Acute Respiratory Distress in Aged, SARS-CoV-2–Infected African Green Monkeys but Not Rhesus Macaques. Am. J. Pathol. 191, 274–282. 10.1016/j.ajpath.2020.10.016.

55. Kim, D., Lee, J.-Y., Yang, J.-S., Kim, J.W., Kim, V.N., and Chang, H. (2020). The Architecture of SARS-CoV-2 Transcriptome. Cell 181, 914–921.e10. 10.1016/j.cell.2020.04.011.

56. Matsuyama, S., Nao, N., Shirato, K., Kawase, M., Saito, S., Takayama, I., Nagata, N., Sekizuka, T., Katoh, H., Kato, F., et al. (2020). Enhanced isolation of SARS-CoV-2 by TMPRSS2-expressing cells. Proc. Natl. Acad. Sci. 117, 7001–7003. 10.1073/pnas.2002589117.

57. Hoffmann, M., Kleine-Weber, H., Schroeder, S., Krüger, N., Herrler, T., Erichsen, S., Schiergens, T.S., Herrler, G., Wu, N.H., Nitsche, A., et al. (2020). SARS-CoV-2 Cell Entry Depends on ACE2 and TMPRSS2 and Is Blocked by a Clinically Proven Protease Inhibitor. Cell 181, 271–280. 10.1016/j.cell.2020.02.052.

58. Smither, S.J., Lear-Rooney, C., Biggins, J., Pettitt, J., Lever, M.S., and Olinger, G.G. (2013). Comparison of the plaque assay and 50% tissue culture infectious dose assay as methods for measuring filovirus infectivity. J. Virol. Methods 193, 565–571. 10.1016/j.jviromet.2013.05.015.

59. Sivula, T., Magnusson, M., Matamoros, A.A., and Vehtari, A. (2022). Uncertainty in Bayesian Leave-One-Out Cross-Validation Based Model Comparison. 10.48550/arXiv.2008.10296.

60. Chicco, D., and Jurman, G. (2020). The advantages of the Matthews correlation coefficient (MCC) over F1 score and accuracy in binary classification evaluation. BMC Genomics 21, 6. 10.1186/s12864-019-6413-7.

61. Stan Development Team (2022). Stan Modeling Language Users Guide and Reference Manual.

62. Matson, M.J., Yinda, C.K., Seifert, S.N., Bushmaker, T., Fischer, R.J., Doremalen, N. van, Lloyd-Smith, J.O., and Munster, V.J. Effect of Environmental Conditions on SARS-CoV-2 Stability in Human Nasal Mucus and Sputum. Emerg. Infect. Dis. 26, 2276–2278. 10.3201/eid2609.202267.

63. Yuan, L., Tang, Q., Zhu, H., Guan, Y., Cheng, T., and Xia, N. (2021). SARS-CoV-2 infection and disease outcomes in non-human primate models: advances and implications. Emerg. Microbes Infect. 10, 1881–1889. 10.1080/22221751.2021.1976598.

64. Grebennikov, D., Kholodareva, E., Sazonov, I., Karsonova, A., Meyerhans, A., and Bocharov, G. (2021). Intracellular Life Cycle Kinetics of SARS-CoV-2 Predicted Using Mathematical Modelling. Viruses 13, 1735. 10.3390/v13091735.

65. Amarilla, A.A., Modhiran, N., Setoh, Y.X., Peng, N.Y.G., Sng, J.D.J., Liang, B., McMillan, C.L.D., Freney, M.E., Cheung, S.T.M., Chappell, K.J., et al. (2021). An Optimized High-Throughput Immuno-Plaque Assay for SARS-CoV-2. Front. Microbiol. 12.

66. Carter, J., and Saunders, V.A. (2007). Virology: Principles and Applications (John Wiley & Sons).

67. Puhach, O., Meyer, B., and Eckerle, I. (2023). SARS-CoV-2 viral load and shedding kinetics. Nat. Rev. Microbiol. 21, 147–161. 10.1038/s41579-022-00822-w.

68. Ørpetveit, I., Mikalsen, A.B., Sindre, H., Evensen, Ø., Dannevig, B.H., and Midtlyng, P.J. (2010). Detection of Infectious Pancreatic Necrosis Virus in Subclinically Infected Atlantic Salmon by Virus Isolation in Cell Culture or Real-Time Reverse Transcription Polymerase Chain Reaction: Influence of Sample Preservation and Storage. J. Vet. Diagn. Invest. 22, 886–895. 10.1177/104063871002200606.

69. Kozlov, M. (2022). NIH issues a seismic mandate: share data publicly. Nature 602, 558–559. 10.1038/d41586-022-00402-1.

70. Russell, W.M.S., and Burch, R.L. (1959). The principles of humane experimental technique (Methuen).

71. Prescott, M.J. (2010). Ethics of primate use. Adv. Sci. Res. 5, 11–22. 10.5194/asr-5-11-2010.

72. Subbaraman, N. (2021). The US is boosting funding for research monkeys in the wake of COVID. Nature 595, 633–634. 10.1038/d41586-021-01894-z.

73. National Primate Research Center (2020). NPRCs Further Collaborations to Overcome Nonhuman Primate Shortage. https://nprc.org/research/nprcs-further-collaborations-to-overcome-nonhuman-primate-shortage/.

74. Kirby, J.E., Riedel, S., Dutta, S., Arnaout, R., Cheng, A., Ditelberg, S., Hamel, D.J., Chang, C.A., and Kanki, P.J. (2023). Sars-Cov-2 antigen tests predict infectivity based on viral culture: comparison of antigen, PCR viral load, and viral culture testing on a large sample cohort. Clin. Microbiol. Infect. 29, 94–100. 10.1016/j.cmi.2022.07.010.

75. Zhang, C., Cui, H., Guo, Z., Chen, Z., Yan, F., Li, Y., Liu, J., Gao, Y., and Zhang, C. (2022). SARS-CoV-2 Virus Culture, Genomic and Subgenomic RNA Load, and Rapid Antigen Test in Experimentally Infected Syrian Hamsters. J. Virol. 96, e01034–22. 10.1128/jvi.01034-22.

76. Ke, R., Martinez, P.P., Smith, R.L., Gibson, L.L., Mirza, A., Conte, M., Gallagher, N., Luo, C.H., Jarrett, J., Zhou, R., et al. (2022). Daily longitudinal sampling of SARS-CoV-2 infection reveals substantial heterogeneity in infectiousness. Nat. Microbiol. 7, 640–652. 10.1038/s41564-022-01105-z.

77. Killingley, B., Mann, A.J., Kalinova, M., Boyers, A., Goonawardane, N., Zhou, J., Lindsell, K., Hare, S.S., Brown, J., Frise, R., et al. (2022). Safety, tolerability and viral kinetics during SARS-CoV-2 human challenge in young adults. Nat. Med., 1–11. 10.1038/s41591-022-01780-9.

