## Supplementary Material for "PCR data accurately predict infectious virus: a characterization of SARS-CoV-2 in non-human primates"

#### Supplementary Methods

##### *Comprehensive literature search*

To construct our database, we conducted comprehensive literature searches on 11 March 2021, which was the chosen cutoff date for inclusion in the database. One author (CS) screened the Web of Science (Core Collection) and PubMed for articles published in English with the following search string: (SARS-CoV-2 OR COVID-19) AND (primate* OR macaque* OR monkey* OR "macaca" OR "chlorocebus"). On the same day, CS also jointly screened bioRxiv and medRxiv, with two separate search strings: (i) (SARS-CoV-2 OR COVID-19) AND (primate* OR macaque* OR monkey*), and (ii) (SARS-CoV-2 OR COVID-19) AND ("macaca" OR "chlorocebus"). These searches returned 163 records from Web of Science, 761 from PubMed, and 259 from bioRxiv and medRxiv. An additional 60 records were obtained from Google Scholar searches and citation trackers (before 11 March 2021) using the following search terms: SARS-CoV-2, COVID-19, macaque, monkey, non-human primate (**Figure S1**). In total, this returned 1,243 results, with 866 unique records after removing duplicates. Of those, the following were immediately excluded: (i) articles published before 2020, (ii) article types that do not generate primary data (e.g., opinions, reviews), and (iii) articles with clearly irrelevant titles based on our predefined eligibility/inclusion criteria (described in the **Methods**). CS inspected the abstracts of the remaining 275 studies and the full texts of 122 records, according to the same eligibility criteria.

##### *Data collection process*

For each included article and every infected individual, the following study design details were obtained from the text, figures, supplementary files, or the corresponding authors, when available: primate species, rhesus macaque origin (Chinese or Indian), ID, sex, age (years or months), age class, treatment group, viral inoculum strain, inoculation route(s), and route specific inoculation dose(s). For every sample, the following information was recorded when available: sample value (converted to log10, when quantitative and reported otherwise), sample time (day post infection, starting at 0), sample type (e.g., swab, tissue), sample location (e.g., liver), sample units (e.g., viral RNA copies/mL), method of quantification (e.g., RT-qPCR, plaque assay), target gene (for PCR), cell line (e.g., VeroE6, for viral culture), and limit of detection and/or quantification. We standardized all ID names as follows: [the first and last initial of the study's first author] _ [the ID name as assigned in the original study, if available].

When raw data was not published or methodological/biological details were missing or inconsistently reported, we contacted the corresponding author(s) using a standardized email template (paired with study-specific questions) to resolve discrepancies and/or request raw data. When clarification and/or raw data were obtained, the details were updated accordingly. In instances where clarification was not obtained but conflicting information existed in the article, we recorded the information most consistent with the methods section. Unclarified information that was missing entirely from the article materials is considered unknown.

***Standardizing age class and inoculation doses***

We developed a consistent method to assign age classes for all studies, following methods used in prior studies.[1–5] When age was reported in months and years, juvenile rhesus and cynomolgus macaques include individuals with ages <5 years, adults include 5-19 years, and geriatrics include ≥20 years. Juvenile African green monkeys include ages <5 years, adults include 5-14 years, and geriatrics include ≥15 years. When reported age ranges spanned multiple assignments, we used the reported age class if available, otherwise we considered it unknown.

Since inoculation doses are frequently reported as either plaque forming units (pfu) or tissue culture infectious dose 50 (TCID50), we converted all inoculation doses reported in TCID50 to pfu using a standard conversion factor (1 TCID50=0.69 pfu).[6]

***Confirming RNA types***

To confirm whether articles quantified full-length genomic RNA, subgenomic RNA, or total RNA (since reporting practices varied across studies), all relevant primer and probe sequences were extracted from each study or from referenced articles. We used SnapGene software (from Insightful Science; available at snapgene.com) to map all sequences to the SARS-CoV-2/human/CHN/Wu-1 reference genome. Any assays with both forward and reverse primers located within the ORF1ab gene were considered to amplify full-length genomic RNA, as this gene is not located on any canonical subgenomic RNAs. Assays with both forward and reverse primers located within any individual gene downstream of ORF1ab were considered to amplify total RNA, as these sequences can be found on both subgenomic and full-length genomic RNA. Assays where the forward primer was located in the 5' UTR but the reverse primer was located within a gene downstream of ORF1ab were classified as amplifying subgenomic RNA.

***Justification and prior description for candidate predictors***

Below, we provide further justification for the selection of all candidate predictors, including descriptions of all assigned priors. For the sgRNA models (that predict sgRNA results from totRNA), we use $\gamma/\delta$ to signify parameters for the logistic component and $\alpha/\beta$ for the linear component. For the culture model (that predicts culture positivity from totRNA quantities), we use $\gamma/\psi$ to signify the relevant parameters.

<u>*Primary predictors:*</u> We required all sgRNA models to include total RNA copy number (T) as a predictor, given this is the primary effect of interest. We expect the probability of detecting sgRNA to increase as total RNA copy numbers increase, which is supported by studies that find more sgRNA positive samples for those with large quantities of total RNA.[7,8] Given that we expect this relationship to be positive, we assign the following prior: $\delta_T \sim N(2,1)$. We also expect sgRNA copy numbers to increase with total RNA copy numbers, as observed in other studies that find positive linear relationships between them.[8–10] We assign the following prior: $\beta_T \sim Gamma(2,0.5)$. We evaluated culture models including total RNA, sgRNA (SG), or both as primary predictors. We expect the likelihood of detecting infectious virus to increase with increasing quantities of either RNA type, so we set the priors to be: $\Psi_T, \Psi_{SG} \sim N(2, 1)$.

<u>*Age, sex, and non-human primate species:*</u> As with other viruses, age (AGE) and sex (SEX) are hypothesized to affect individual responses to SARS-CoV-2 infection, including within-host infection kinetics and disease severity. Inter-species variability in infection dynamics has also been noted among non-human primate species (SP) and other animal models.[11] These differences may affect observed relationships between assays. However, given the complexity of these biological interactions, we do not assign *a priori* expectations about the direction of these effects, and we set all associated priors to be $N(0,1)$.

<u>*Inoculation dose:*</u> Since sgRNA is not typically packaged into virions[12] and thus should not meaningfully exist in viral stocks, we expect to find detectable levels of total RNA earlier than sgRNA after inoculation (i.e., lower probabilities of sgRNA detection per total RNA quantity for higher doses; $\delta_{DOSE} \sim N(-1,0.5)$). Larger inoculum doses could also result in (at least initially) higher levels of total RNA relative to sgRNA ($\beta_{DOSE} \sim N(-0.25,1)$. Exposure dose affects virion quantity and hence also total RNA quantity, which in turn may affect the probability of culture positivity through the presence of

residual inoculum or other effects on infection kinetics. We make no *a priori* predictions on the direction of this effect ($\Psi_{DOSE} \sim N(0,1)$).

<u>Day post infection</u>: sgRNA levels likely remain low for some time in recently infected tissues. They may rise as virions infect local cells and replicate, and the ratio between total RNA and sgRNA may stabilize.[8] These dynamic changes may be especially pronounced when large quantities of virions (and thus also genomic RNA) are introduced simultaneously, as in animal infection experiments. Consequently, we expect newly infected tissues to contain small, likely undetectable quantities of sgRNA, whereas tissues infected one or more days prior, which have experienced sufficient replication, likely contain more sgRNA copies. For infectious virus, residual inoculum-derived virus may actually increase the probability of culture positivity in infected tissues relative to non-inoculated tissues (where virions must be produced). We derived a categorical predictor with three levels: DPI[1], DPI[2], and DPI[3]. For inoculated tissues, we distinguish between the first day (DPI[1]) and all other days post infection (DPI[2]). Since infection timing is unknown for non-inoculated tissues, we group all days into one level (DPI[3]). **Table S3** lists which tissues were considered inoculated for each exposure procedure. We used the following priors: $\delta_{DPI[1]} \sim N(-1,1)$, $\delta_{DPI[2]} \sim N(1,1)$, $\delta_{DPI[3]} \sim N(0,1)$, $\beta_{DPI[1]} \sim N(-0.5,1)$ $\beta_{DPI[2]} \sim N(0.5,1)$, $\beta_{DPI[3]} \sim N(0,1)$, $\Psi_{DPI[1]} \sim N(1,1)$, $\Psi_{DPI[2]} \sim N(0.5,1)$, $\Psi_{DPI[3]} \sim N(0,1)$.

<u>Sample type:</u> Differences in the processing and content of non-invasive (e.g., swabs, biofluids) and invasive (e.g., whole tissues obtained at necropsy) samples may affect assay readouts, and so we include sample type as a candidate predictor. We make no *a priori* predictions on their differences, and instead we assign non-informative priors ($\delta_{ST},\beta_{ST},\Psi_{ST} \sim N(0,1)$).

<u>Target gene:</u> The RT-qPCR target gene affects quantification of SARS-CoV-2 copy numbers,[8,10,13,14] which stems from the nature of PCR in addition to the organizational structure and transcription mechanisms of the coronavirus genome. The production and abundance of sgRNA species varies by gene (e.g., sgRNA N > sgRNA E[15]), and sgRNA assays only amplify gene-specific sgRNA transcripts. In contrast, total RNA assays can amplify not only full-length genomic RNA and gene-specific sgRNA, but also other, larger sgRNAs that contain the target sequence. Given the many possible combinations of target gene pairs for total RNA and sgRNA assays, in our pooled dataset, we derived a generalizable predictor based on the number of transcripts available for amplification for each protocol. We distinguished between totRNA assays that amplify most (i.e., targets the Nucleocapsid gene; termed 'totRNA-high') or few sgRNA species (i.e., targets the Envelope gene; 'totRNA-low'). We also distinguish between sgRNA assays that target highly expressed (i.e., sgN; 'sgRNA-high') or less

expressed sgRNA species (i.e., sgE, sg7; 'sgRNA-low'). This predictor thus has the following four levels: (1) totRNA-high/sgRNA-high, (2) totRNA-low/sgRNA-high, (3) totRNA-high/sgRNA-low, and (4) totRNA-low/sgRNA-low. We expect sgRNA-high assays to have higher probabilities of sgRNA detection and higher quantities of sgRNA per totRNA quantity ($\delta_{TG[1]}$, $\delta_{TG[2]} \sim N(1,1)$; $\beta_{TG[1]}$, $\beta_{TG[2]} \sim N(0.5,1)$) than sgRNA-low assays ($\delta_{TG[3]}$, $\delta_{TG[4]} \sim N(-1,1)$; $\beta_{TG[3]}$, $\beta_{TG[4]} \sim N(-0.5,1)$). For all culture models containing total RNA, we simply used the total RNA target gene as the predictor. When predicting culture from sgRNA, we used the sgRNA target gene as the predictor. Given SARS-CoV-2's genomic structure, we expect decreasing per-sample viral loads for totRNA protocols targeting the Nucleocapsid ($\Psi_{TG[1]} \sim N(-1,1)$), Envelope ($\Psi_{TG[2]} \sim N(0,1)$), and Spike ($\Psi_{TG[3]} \sim N(1,1)$) genes, respectively. Since these stoichiometric ratios apply in any infected cell, we expect the relative probability of culture positivity to be higher for target genes with lower expression, such that the probability is highest for Spike ($\Psi_{TG[3]} \sim N(1,1)$), intermediate for Envelope ($\Psi_{TG[2]} \sim N(0,1)$), and lowest for Nucleocapsid ($\Psi_{TG[1]} \sim N(-1,1)$).

_Cell line:_ Viral infectivity can vary by cell line, which can influence the likelihood of detecting infectious virus in a sample. Our data includes three distinct cell lines (VeroE6, Vero76, VeroE6-TMPRSS2) which we group into a three-level categorical predictor (CELL) for the culture model only. Given evidence of increased SARS-CoV-2 entry in cells expressing TMPRSS2,[16,17] we expect the probability of detection to be highest for VeroE6-TMPRSS2 cells ($\Psi_{CELL[3]} \sim N(1,1)$). We use the same non-informative prior for the other two ($\Psi_{CELL[1]} \sim N(0,1)$, $\Psi_{CELL[2]} \sim N(0,1)$).

_Culture assay:_ The sensitivity of endpoint dilution (TCID50) and plaque assays can also vary, with plaque assays typically having lower sensitivity.[18] We distinguish between these protocols in a binary predictor (ASSAY), with endpoint dilution treated as the reference. We assign the following prior ($\Psi_{ASSAY} \sim N(-0.5,1)$).

##### *Approximate leave-one-out cross-validation*

Beyond standard 10-fold cross-validation used in our main analyses, we also evaluated model performance and conducted model selection using Pareto-Smoothed Importance Sampling Approximate Leave-One-Out cross-validation (PSIS-LOO) via the 'loo' R package.[19] We ran this method for the linear component of the sgRNA model using RStan version 2.21.0 to use additional functionality of the LOO software.[20] All other models were run using CmdStan, as described in the main **Methods**.

Pareto-k values indicate the accuracy of PSIS-LOO approximations, where values below 0.7 indicate sufficiently reliable estimates. We use the moment-matching functionality offered by the 'loo' package to correct Pareto-k values exceeding 0.7, after which all PSIS-LOO estimates generated in this study were reliable (adjusted Pareto-k < 0.7).

##### *Cross-validation methods and model evaluation metrics*

For 10-fold cross-validation, we assigned folds randomly, although we required each fold to contain similar quantities of data from each article. This allowed us to distribute protocols, demographics, and sampling types more evenly across folds. All statistics for this method were calculated separately for training and test sets to evaluate performance on out-of-sample data and assess potential bias or overfitting, with the exception of ELPD and MCC which were only calculated for test sets.

As outlined in the model description of the main **Methods**, the linear component of the sgRNA models contained article-specific hierarchical error rates. We used the estimates of each article's specific error distribution to generate ELPD values. However, when we calculated median absolute error and the percent of samples falling within given prediction intervals, we did not incorporate article-specific errors. Instead, for each iteration, we sampled the full range of estimated errors by sampling the error distribution of a random included article (uniformly distributed), so these statistics better correspond to performance on new data (e.g., new studies) where prior information on error distributions is unavailable.

##### *Selection procedure and results for the best sgRNA model*

In the paragraphs below, we describe our model selection procedure for the sgRNA model, which relied on various performance metrics for each component. Overall, this procedure clearly identified the best model for both the logistic and linear components of the sgRNA model (**Figure 2**). Parallel analyses conducted with Pareto-Smoothed Importance Sampling approximate leave-one-out cross-validation resulted in qualitatively similar outcomes and selection of the same model (**Tables S2, S3**). Model selection also did not vary when predictors were or were not centered and scaled. Sensitivity analyses confirmed qualitatively similar results between informative and non-informative priors for the model selection procedure and for the parameter estimates of the best model (**Figure S12**). Prediction accuracy and median absolute error (MAE) showed minimal differences between training and test datasets, for all PCR models considered, offering confidence in the models' generalizability (**Tables S2, S3**).

For the logistic component, the best model performed substantially better than the simple model for all three statistics considered. The best model had considerably higher ELPD (difference: 67.3; **Figure 2, Table S2**). Prediction accuracy on test data increased overall by 3.4 percent, and MCC increased from 0.75 to 0.82 (**Figure 2, Figure 3C, Table S2**). The best model also correctly predicted sgRNA detectability with higher probability for more samples and had a substantially higher ELPD, both of which reflect higher certainty for true classifications (**Figure 2, Figure 3C**). Prediction accuracy and Matthews correlation coefficient (MCC) indicate nearly identical performance of the best and full models (**Figure 2**, **Table S2**). The difference between the estimated log pointwise predictive density (ELPD) for the best and full models falls within two standard errors of the difference (difference: 1.60; standard error: 3.50), again indicating similar model performance.[21]

When predicting copy numbers for all sgRNA positive samples with the linear component, the best model showed better performance than the full and simple models on all three statistics considered. Predictive performance of the best model is substantially better than the simple model, with 55.0 versus 48.0 percent of samples falling within the 50 percent prediction interval (**Figure 2, Table S3**) and a decrease in median prediction error from 0.58 to 0.44 (**Figure 3F**). The ELPD difference is also substantial (-186.2). Posterior predictive checks show a strong correlation between observed and median predicted sgRNA values for the best model (adjusted $R^2$=0.77), which further supports superior performance of the multiple regression model compared to the simple single regression model (adjusted $R^2$=0.68). Relative to the full model, the best model has higher ELPD (difference: -4.44) and similar prediction error on test data (0.44 vs. 0.45) (**Figure 2, Table S3**). A higher percent of (test) samples fall within the model-generated 50 percent prediction interval for the best model (55.0 vs. 54.5).

##### *Selection procedure and results for the best culture model*

In the paragraphs below, we describe our model selection procedure for the culture model, which relied on three primary performance metrics. This procedure clearly identified the best culture model. Model selection did not vary between cross-validation methods or when centering and scaling predictors. Prior choice also did not alter model selection nor did it qualitatively affect parameter estimates (**Figure S12**). Prediction accuracy between training and test sets was comparable (**Table S4**), again supporting model generalizability.

The evaluation procedure identified the best model as the one containing all but two candidate predictors (**Figure 2**). For all three statistics considered, the best model performed better than the simple model. The best model had considerably higher ELPD, for which the difference was larger than two standard errors (difference: 55.70; standard error: 13.96; **Figure 2, Table S4**). For totRNA-positive samples, prediction accuracy on test data increased overall by 3.1 percent, but notably the best model correctly classified an additional 7.0 percent of culture positive samples (**Figure 4C**), both of which are reflected in the improvement of MCC from 0.48 to 0.57 (**Table S4**). The difference between the ELPD for the best and full models is small (difference: 0.76; standard error: 3.42), and their overall prediction accuracy and MCC are nearly identical (**Figure 2**, **Table S4**).

***Mathematical form of the best sgRNA model***

The mathematical form of the best sgRNA model is outlined below, where we use γ/δ to signify parameters for the logistic component and α/β for the linear component. Acronyms are as follows: SG: sgRNA, T: total RNA, DOSE: inoculation dose (log10 pfu), SP: species, TG: target gene (standardized to four levels), and DPI: day post infection.

$$SG_{>LOD,i} \sim Bernoulli(p_i)$$

$$logit(p_i) = \gamma + \delta_T T_i + \delta_{DOSE} DOSE_i + \delta_{TG_i} + \delta_{SP_i}$$

$$SG_{value,i} \sim N(y_i, \sigma_{article_i})$$

$$y_i = \alpha + \beta_T T_i + \beta_{DOSE} DOSE_i + \beta_{TG_i} + \beta_{DPI_i} + \beta_{SP_i}$$

$$\sigma_{article_i} \sim N(\bar{\sigma}, \sigma_{sd})$$

***Mathematical form of the best culture model***

The mathematical form of the best culture model is outlined below, where we use γ/ψ to signify the relevant parameters. Acronyms are as follows: C: culture, T: total RNA, DOSE: inoculation dose (log10 pfu), DPI: day post infection, AGE: age class, SP: species, CELL: culture cell line, ASSAY: culture assay, and TG: target gene (standardized to three levels).

$$C_{>LOD,i} \sim Bernoulli(p_i)$$

$$logit(p_i) = \gamma + \psi_T T_i + \psi_{DOSE} DOSE_i + \psi_{DPI_i} + \psi_{AGE_i} + \psi_{SP_i} +$$

$$\psi_{CELL_i} + \psi_{ASSAY} ASSAY_i + \psi_{TG_i}$$

***Prediction intervals and parameter estimates***

To generate prediction intervals for various statistics, including median probabilities of sgRNA or culture positivity and median predicted sgRNA copy numbers, we used all available post-warmup parameter samples from the best model. We used the same procedure to generate the fit lines shown in Figures 3 and 5. Each prediction is generated using grouped parameter samples (e.g., samples from the same chain and iteration) to preserve correlation structure.

***Estimating differences between predicted and known outcomes***

To compare median predicted sgRNA copies to observed sgRNA or totRNA copies, we modeled the distribution of each of these quantities as normally distributed random variables with unique means and variances. We also compared totRNA quantities for culture samples with false negative, true positive, and true negative predictions by estimating the population means for each group (captured as a normally distributed variable). For both of these investigations, we calculated the median and quantiles of the distribution of differences between these populations means, where we subtracted estimates only from the same chain and iteration. We fit these Bayesian models using the same procedures as described in the **Methods.**

#### References


1. Cramer, P.E., Gentzel, R.C., Tanis, K.Q., Vardigan, J., Wang, Y., Connolly, B., Manfre, P., Lodge, K., Renger, J.J., Zerbinatti, C., et al. (2018). Aging African green monkeys manifest transcriptional, pathological, and cognitive hallmarks of human Alzheimer's disease. Neurobiol. Aging *64*, 92–106. 10.1016/j.neurobiolaging.2017.12.011.

2. Lee, J.-R., Choe, S.-H., Kim, Y.-H., Cho, H.-M., Park, H.-R., Lee, H.-E., Jin, Y.B., Kim, J.-S., Jeong, K.J., Park, S.-J., et al. (2020). Longitudinal profiling of the blood transcriptome in an African green monkey aging model. Aging *13*, 846–864. 10.18632/aging.202190.

3. Simmons, H.A. (2016). Age-Associated Pathology in Rhesus Macaques (Macaca mulatta). Vet. Pathol. *53*, 399–416. 10.1177/0300985815620628.

4. Darusman, H.S., Call, J., Sajuthi, D., Schapiro, S.J., Gjedde, A., Kalliokoski, O., and Hau, J. (2014). Delayed response task performance as a function of age in cynomolgus monkeys (Macaca fascicularis). Primates *55*, 259–267. 10.1007/s10329-013-0397-8.

5. Koo, B.-S., Lee, D.-H., Kang, P., Jeong, K.-J., Lee, S., Kim, K., Lee, Y., Huh, J.-W., Kim, Y.-H., Park, S.-J., et al. (2019). Reference values of hematological and biochemical parameters in young-adult cynomolgus monkey (Macaca fascicularis) and rhesus monkey (Macaca mulatta) anesthetized with ketamine hydrochloride. Lab. Anim. Res. *35*, 1–6. 10.1186/s42826-019-0006-0.

6. Carter, J., and Saunders, V.A. (2007). Virology: Principles and Applications (John Wiley & Sons).

7. Perera, R.A.P.M., Tso, E., Tsang, O.T.Y., Tsang, D.N.C., Fung, K., Leung, Y.W.Y., Chin, A.W.H., Chu, D.K.W., Cheng, S.M.S., Poon, L.L.M., et al. (2020). SARS-CoV-2 Virus Culture and Subgenomic RNA for Respiratory Specimens from Patients with Mild Coronavirus Disease - Volume 26, Number 11—November 2020 - Emerging Infectious Diseases journal - CDC. *26*, 2701–2704. 10.3201/eid2611.203219.

8. Dimcheff, D.E., Valesano, A.L., Rumfelt, K.E., Fitzsimmons, W.J., Blair, C., Mirabelli, C., Petrie, J.G., Martin, E.T., Bhambhani, C., Tewari, M., et al. (2021). Severe Acute Respiratory Syndrome Coronavirus 2 Total and Subgenomic RNA Viral Load in Hospitalized Patients. J. Infect. Dis. *224*, 1287–1293. 10.1093/infdis/jiab215.

9. van Kampen, J.J.A., van de Vijver, D.A.M.C., Fraaij, P.L.A., Haagmans, B.L., Lamers, M.M., Okba, N., van den Akker, J.P.C., Endeman, H., Gommers, D.A.M.P.J., Cornelissen, J.J., et al. (2021). Duration and key determinants of infectious virus shedding in hospitalized patients with coronavirus disease-2019 (COVID-19). Nat. Commun. *12*, 267. 10.1038/s41467-020-20568-4.

10. Verma, R., Kim, E., Martínez-Colón, G.J., Jagannathan, P., Rustagi, A., Parsonnet, J., Bonilla, H., Khosla, C., Holubar, M., Subramanian, A., et al. (2021). SARS-CoV-2 Subgenomic RNA Kinetics in Longitudinal Clinical Samples. Open Forum Infect. Dis. *8*, ofab310. 10.1093/ofid/ofab310.

11. Chu, H., Chan, J.F.-W., and Yuen, K.-Y. (2022). Animal models in SARS-CoV-2 research. Nat. Methods *19*, 392–394. 10.1038/s41592-022-01447-w.

12. Escors, D., Izeta, A., Capiscol, C., and Enjuanes, L. (2003). Transmissible Gastroenteritis Coronavirus Packaging Signal Is Located at the 5′ End of the Virus Genome. J. Virol. *77*, 7890–7902. 10.1128/JVI.77.14.7890-7902.2003.

13. Lieberman, J.A., Pepper, G., Naccache, S.N., Huang, M.-L., Jerome, K.R., and Greninger, A.L. (2020). Comparison of Commercially Available and Laboratory-Developed Assays for In Vitro Detection of SARS-CoV-2 in Clinical Laboratories. J. Clin. Microbiol. *58*, e00821-20. 10.1128/JCM.00821-20.

14. Moreira, L.V.L., Luna, L.K. de S., Barbosa, G.R., Perosa, A.H., Chaves, A.P.C., Conte, D.D., Carvalho, J.M.A., and Bellei, N. (2021). Test on stool samples improves the diagnosis of hospitalized patients: Detection of SARS-CoV-2 genomic and subgenomic RNA. J. Infect. *82*, 186–230. 10.1016/j.jinf.2020.11.034.

15. Kim, D., Lee, J.-Y., Yang, J.-S., Kim, J.W., Kim, V.N., and Chang, H. (2020). The Architecture of SARS-CoV-2 Transcriptome. Cell *181*, 914-921.e10. 10.1016/j.cell.2020.04.011.

16. Hoffmann, M., Kleine-Weber, H., Schroeder, S., Krüger, N., Herrler, T., Erichsen, S., Schiergens, T.S., Herrler, G., Wu, N.H., Nitsche, A., et al. (2020). SARS-CoV-2 Cell Entry Depends on ACE2 and TMPRSS2 and Is Blocked by a Clinically Proven Protease Inhibitor. Cell *181*, 271–280. 10.1016/j.cell.2020.02.052.

17. Matsuyama, S., Nao, N., Shirato, K., Kawase, M., Saito, S., Takayama, I., Nagata, N., Sekizuka, T., Katoh, H., Kato, F., et al. (2020). Enhanced isolation of SARS-CoV-2 by TMPRSS2-expressing cells. Proc. Natl. Acad. Sci. *117*, 7001–7003. 10.1073/pnas.2002589117.

18. Smither, S.J., Lear-Rooney, C., Biggins, J., Pettitt, J., Lever, M.S., and Olinger, G.G. (2013). Comparison of the plaque assay and 50% tissue culture infectious dose assay as methods for measuring filovirus infectivity. J. Virol. Methods *193*, 565–571. 10.1016/j.jviromet.2013.05.015.

19. Vehtari, A., Gelman, A., and Gabry, J. (2017). Practical Bayesian model evaluation using leave-one-out cross-validation and WAIC. Stat. Comput. *27*, 1413–1432. 10.1007/s11222-016-9696-4.

20. Vehtari, A., Gabry, J., Magnusson, M., Yuling, Y., Bürkner, P.-C., Paananen, T., and Gelman, A. (2022). loo: Efficient leave-one-out cross-validation and WAIC for Bayesian models.

21. Sivula, T., Magnusson, M., Matamoros, A.A., and Vehtari, A. (2022). Uncertainty in Bayesian Leave-One-Out Cross-Validation Based Model Comparison. 10.48550/arXiv.2008.10296.

### Supplementary Figures

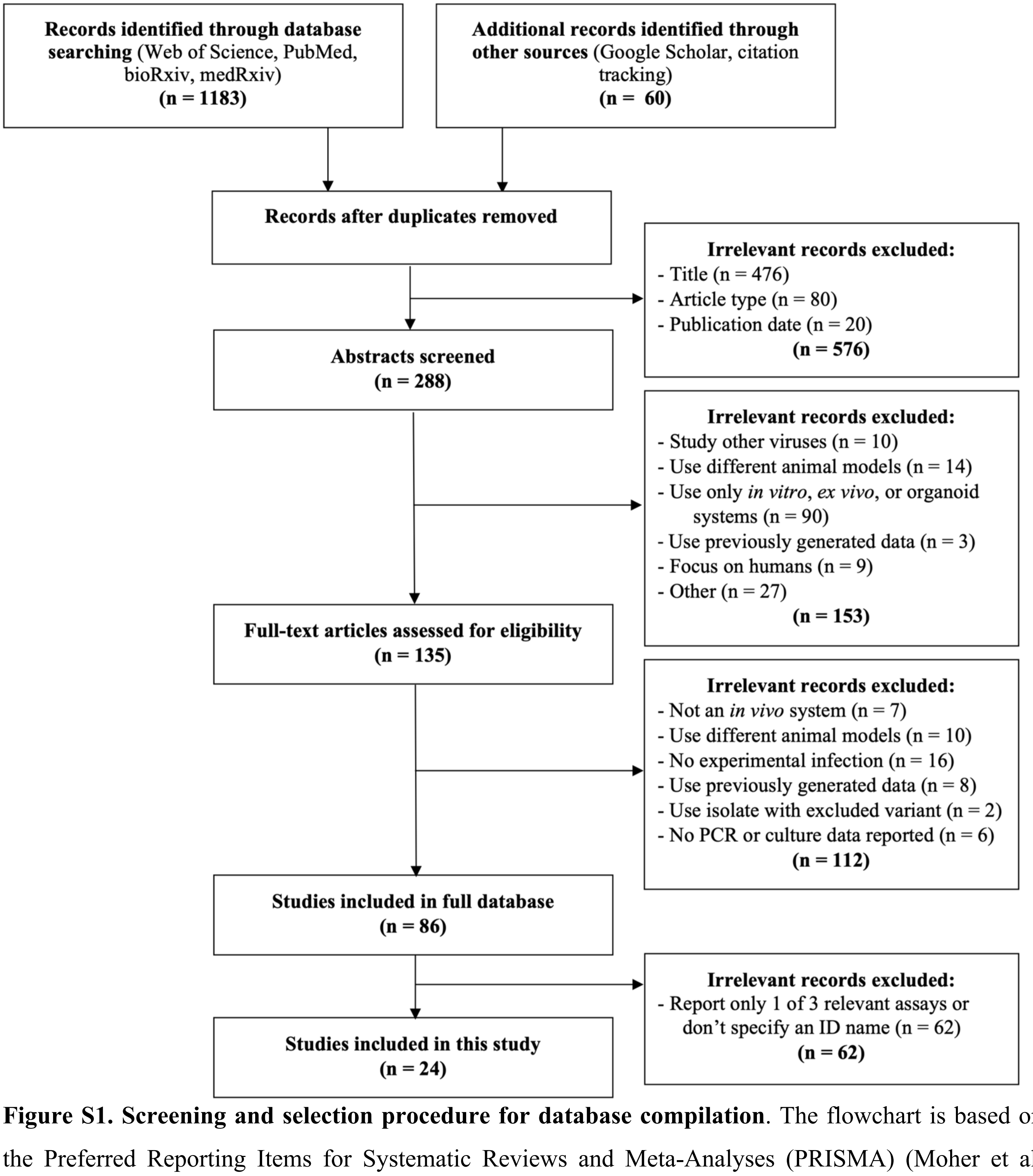


**Figure S1. Screening and selection procedure for database compilation**. The flowchart is based on the Preferred Reporting Items for Systematic Reviews and Meta-Analyses (PRISMA) (Moher et al. 2009). Additional information on the screening procedure is provided in the **Supplementary Methods**.

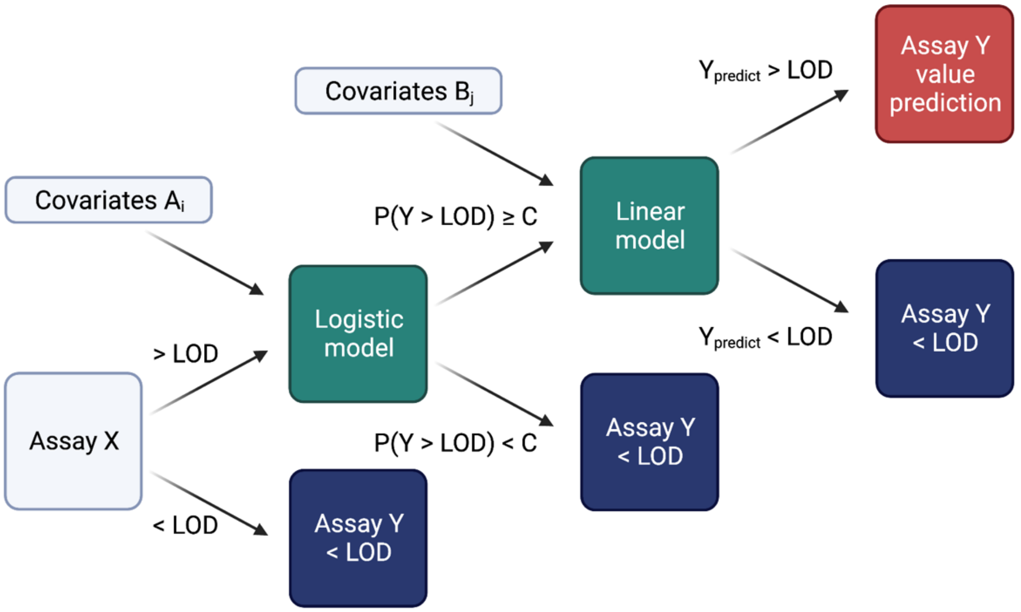


**Figure S2. Schematic diagram of generalizable hurdle model predicting assay Y from a more sensitive assay X**. Predictors are grey, model components are green, and predictions are red (positive) or blue (negative). If assay X falls below the limit of detection (< LOD), assay Y is also predicted to fall below the limit of detection. (Note that this particular assumption may not hold for all assay relationships, and modeling adjustments may need to be made in these scenarios.) If assay X falls above the limit of detection (> LOD), then the value of assay X is passed as a predictor to the logistic component of the hurdle model, which uses a set of additional covariates $A_i$ to predict whether assay Y falls above or below the LOD. If the posterior probability of assay Y falling above the limit of detection is less than some assigned threshold C (P(Y > LOD) < C), then the model predicts assay Y falls below the LOD. Otherwise, the model predicts assay Y falls above the LOD. Note that the probability cut-off value C should be selected to balance false positive and false negative rates as appropriate to investigator aims. In this study, we used a standard value of C=0.5. For samples predicted to fall above the LOD, the linear model component will generate a predicted value of assay Y ($Y_{predict}$) based on another set of covariates ($B_j$). If $Y_{predict}$ is larger than the reported LOD for assay Y, the model will return the predicted value. If $Y_{predict}$ is less than the reported LOD for assay Y, the model will predict assay Y falls below the LOD. This figure was created with BioRender.com.

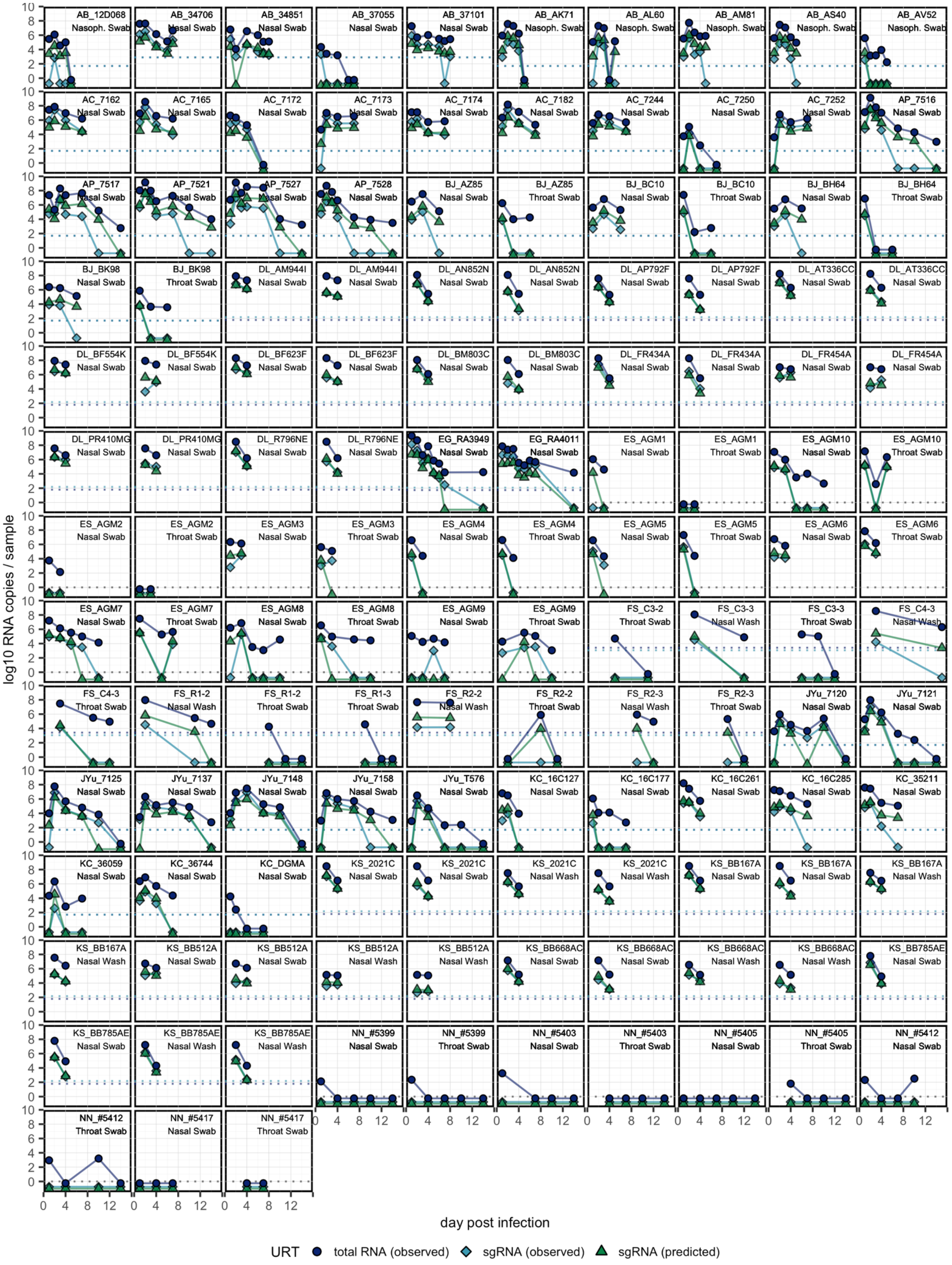
AB_12D068 Nasoph. Swab
AB_34706 Nasal Swab
AB_34851 Nasal Swab
AB_37055 Nasal Swab
AB_37101 Nasal Swab
AB_AK71 Nasoph. Swab
AB_AL60 Nasoph. Swab
AB_AM81 Nasoph. Swab
AB_AS40 Nasoph. Swab
AB_AV52 Nasoph. Swab
AC_7162 Nasal Swab
AC_7165 Nasal Swab
AC_7172 Nasal Swab
AC_7173 Nasal Swab
AC_7174 Nasal Swab
AC_7182 Nasal Swab
AC_7244 Nasal Swab
AC_7250 Nasal Swab
AC_7252 Nasal Swab
AP_7516 Nasal Swab
AP_7517 Nasal Swab
AP_7521 Nasal Swab
AP_7527 Nasal Swab
AP_7528 Nasal Swab
BJ_AZ85 Nasal Swab
BJ_AZ85 Throat Swab
BJ_BC10 Nasal Swab
BJ_BC10 Throat Swab
BJ_BH64 Nasal Swab
BJ_BH64 Throat Swab
BJ_BK98 Nasal Swab
BJ_BK98 Throat Swab
DL_AM944I Nasal Swab
DL_AM944I Nasal Swab
DL_AN852N Nasal Swab
DL_AN852N Nasal Swab
DL_AP792F Nasal Swab
DL_AP792F Nasal Swab
DL_AT336CC Nasal Swab
DL_AT336CC Nasal Swab
DL_BF554K Nasal Swab
DL_BF554K Nasal Swab
DL_BF623F Nasal Swab
DL_BF623F Nasal Swab
DL_BM803C Nasal Swab
DL_BM803C Nasal Swab
DL_FR434A Nasal Swab
DL_FR434A Nasal Swab
DL_FR454A Nasal Swab
DL_FR454A Nasal Swab
DL_PR410MG Nasal Swab
DL_PR410MG Nasal Swab
DL_R796NE Nasal Swab
DL_R796NE Nasal Swab
EG_RA3949 Nasal Swab
EG_RA4011 Nasal Swab
ES_AGM1 Nasal Swab
ES_AGM1 Throat Swab
ES_AGM10 Nasal Swab
ES_AGM10 Throat Swab
ES_AGM2 Nasal Swab
ES_AGM2 Throat Swab
ES_AGM3 Nasal Swab
ES_AGM3 Throat Swab
ES_AGM4 Nasal Swab
ES_AGM4 Throat Swab
ES_AGM5 Nasal Swab
ES_AGM5 Throat Swab
ES_AGM6 Nasal Swab
ES_AGM6 Throat Swab
ES_AGM7 Nasal Swab
ES_AGM7 Throat Swab
ES_AGM8 Nasal Swab
ES_AGM8 Throat Swab
ES_AGM9 Nasal Swab
ES_AGM9 Throat Swab
FS_C3-2 Throat Swab
FS_C3-3 Nasal Wash
FS_C3-3 Throat Swab
FS_C4-3 Nasal Wash
FS_C4-3 Throat Swab
FS_R1-2 Nasal Wash
FS_R1-2 Throat Swab
FS_R1-3 Throat Swab
FS_R2-2 Nasal Wash
FS_R2-2 Throat Swab
FS_R2-3 Nasal Wash
FS_R2-3 Throat Swab
JYu_7120 Nasal Swab
JYu_7121 Nasal Swab
JYu_7125 Nasal Swab
JYu_7137 Nasal Swab
JYu_7148 Nasal Swab
JYu_7158 Nasal Swab
JYu_T576 Nasal Swab
KC_16C127 Nasal Swab
KC_16C177 Nasal Swab
KC_16C261 Nasal Swab
KC_16C285 Nasal Swab
KC_35211 Nasal Swab
KC_36059 Nasal Swab
KC_36744 Nasal Swab
KC_DGMA Nasal Swab
KS_2021C Nasal Swab
KS_2021C Nasal Swab
KS_2021C Nasal Wash
KS_2021C Nasal Wash
KS_BB167A Nasal Swab
KS_BB167A Nasal Swab
KS_BB167A Nasal Wash
KS_BB167A Nasal Wash
KS_BB512A Nasal Swab
KS_BB512A Nasal Swab
KS_BB512A Nasal Wash
KS_BB512A Nasal Wash
KS_BB668AC Nasal Swab
KS_BB668AC Nasal Swab
KS_BB668AC Nasal Wash
KS_BB668AC Nasal Wash
KS_BB785AE Nasal Swab
KS_BB785AE Nasal Swab
KS_BB785AE Nasal Wash
KS_BB785AE Nasal Wash
NN_#5399 Nasal Swab
NN_#5399 Throat Swab
NN_#5403 Nasal Swab
NN_#5403 Throat Swab
NN_#5405 Nasal Swab
NN_#5405 Throat Swab
NN_#5412 Nasal Swab
NN_#5412 Throat Swab
NN_#5417 Nasal Swab
NN_#5417 Throat Swab
log10 RNA copies / sample
day post infection
URT
total RNA (observed)
sgRNA (observed)
sgRNA (predicted)

**Figure S3. Individual viral load trajectories in the upper respiratory tract, including sgRNA predictions generated by the best sgRNA model.** Each panel corresponds with one individual and one non-invasive sample type, indicated in the top right of each panel. Only individuals with both total RNA and sgRNA results for at least two days post infection are plotted. Some individuals were sampled from multiple locations in the upper respiratory tract, in which case they are plotted as neighboring panels. Each line and the accompanying points track the individual's total RNA (dark blue, circle), observed sgRNA (light blue, diamond), and median predicted sgRNA (green, triangle) trajectories. For some individuals (e.g., KS_2021C), multiple RT-qPCR assays targeting different genes were run on the same sample, which are plotted as distinct panels. All samples observed or predicted to fall below the limit of detection are plotted below 0 at set values for visual clarity (totRNA: -0.5, observed sgRNA: -0.75, predicted sgRNA: -1). When available, the limits of detection (LOD) or quantification (LOQ) for PCR assays are plotted as dotted lines in the assay-specific color. When both the LOD and LOQ were available, only the LOD is plotted. In instances where the total RNA and sgRNA assay LOD are equal, only the sgRNA line is visible. No instances exist in this dataset where the LOD or LOQ is only available for one RNA type.

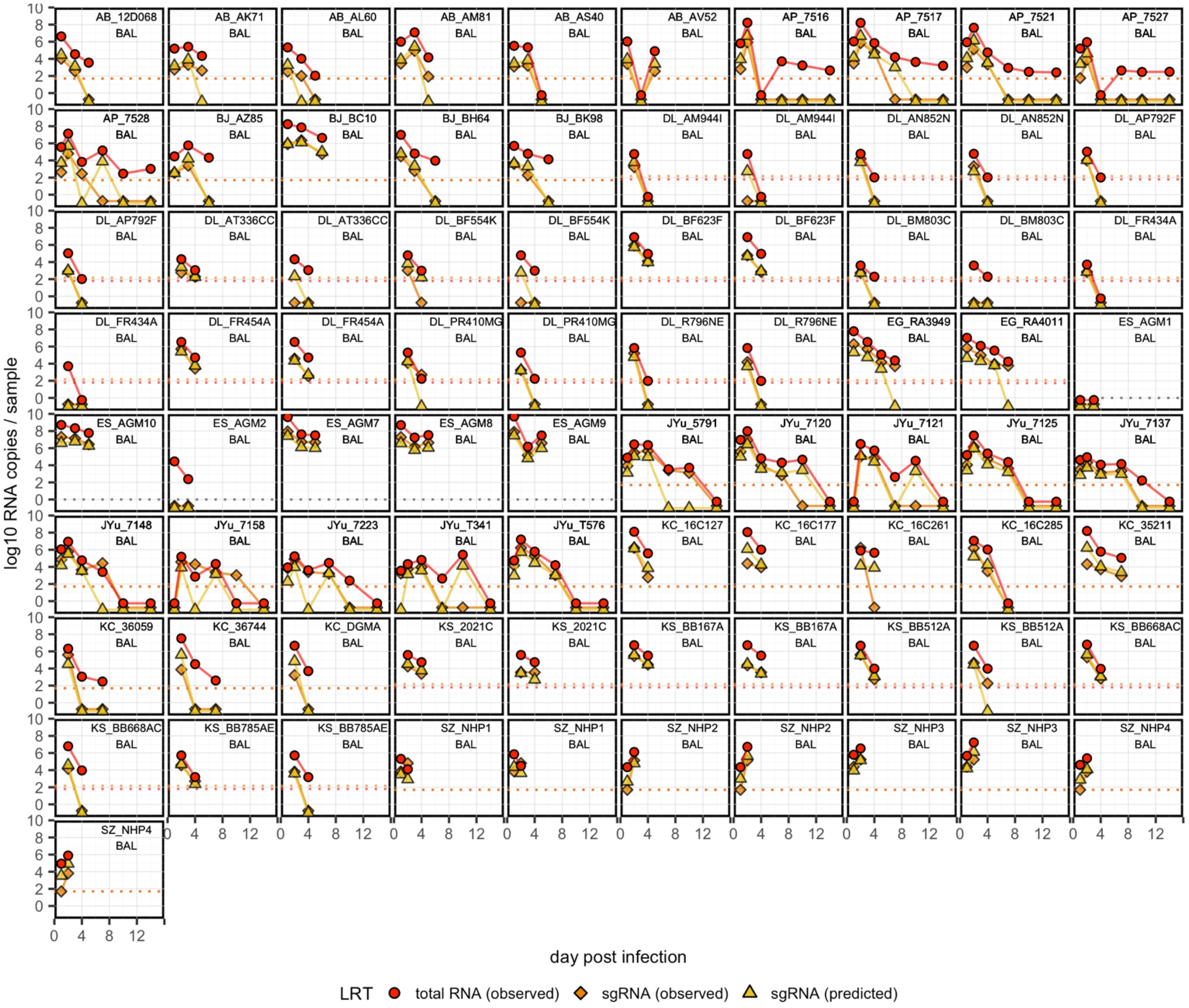


**Figure S4. Individual viral load trajectories in the lower respiratory tract, including sgRNA predictions generated by the best sgRNA model.** Each panel corresponds with one individual and one non-invasive sample type, indicated in the top right of each panel ('BAL': bronchoalveolar lavage). Only individuals with both total RNA and sgRNA results for at least two days post infection are plotted. Each line and the accompanying points track the individual's total RNA (dark red, circle), observed sgRNA (orange, diamond), and median predicted sgRNA (yellow, triangle) trajectories. For some individuals (e.g., KS_2021C), multiple RT-qPCR assays targeting different genes were run on the same sample, which are plotted as distinct panels. All samples observed or predicted to fall below the limit of detection are plotted below 0 at set values for visual clarity (totRNA: -0.5, observed sgRNA: -0.75, predicted sgRNA: -1). When available, the limits of detection (LOD) or quantification (LOQ) for PCR assays are plotted as dotted lines in the assay-specific color. When both the LOD and LOQ were available, only the LOD is plotted. In instances where the total RNA and sgRNA assay

LOD are equal, only the sgRNA line is visible. No instances exist in this dataset where the LOD or LOQ is only available for one RNA type.

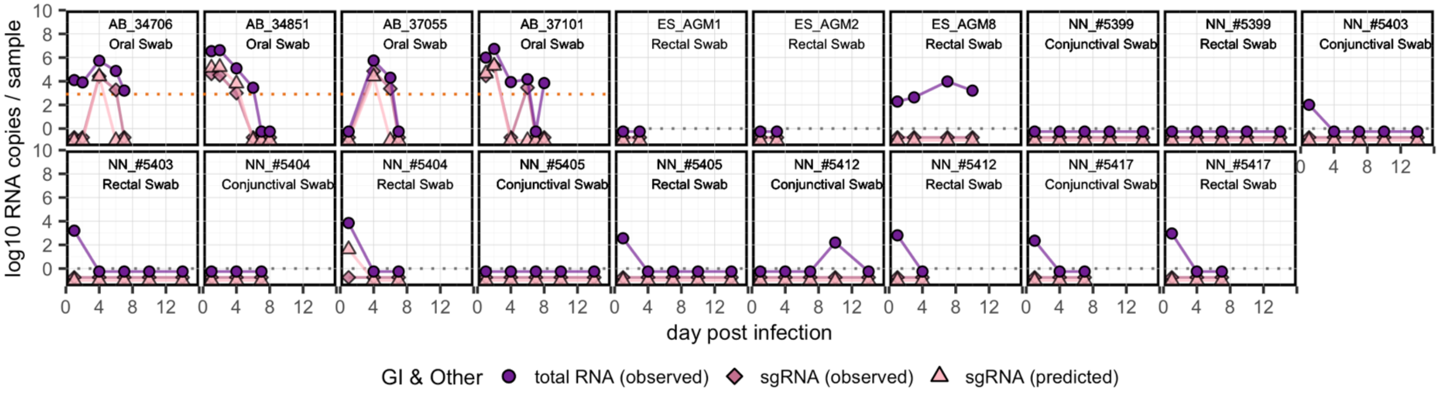


**Figure S5. Individual viral load trajectories in the gastrointestinal and other systems, including sgRNA predictions generated by the best sgRNA model.** Each panel corresponds with one individual and one non-invasive sample type, indicated in the top right of each panel. Only individuals with both total RNA and sgRNA results for at least two days post infection are plotted. Each line and the accompanying points track the individual's total RNA (dark purple, circle), observed sgRNA (dark pink, diamond), and median predicted sgRNA (light pink, triangle) trajectories. All samples observed or predicted to fall below the limit of detection are plotted below 0 at set values for visual clarity (totRNA: -0.5, observed sgRNA: -0.75, predicted sgRNA: -1). When available, the limits of detection (LOD) or quantification (LOQ) for PCR assays are plotted as dotted lines in the assay-specific color. When both the LOD and LOQ were available, only the LOD is plotted. In instances where the total RNA and sgRNA assay LOD are equal, only the sgRNA line is visible. No instances exist in this dataset where the LOD or LOQ is only available for one RNA type.

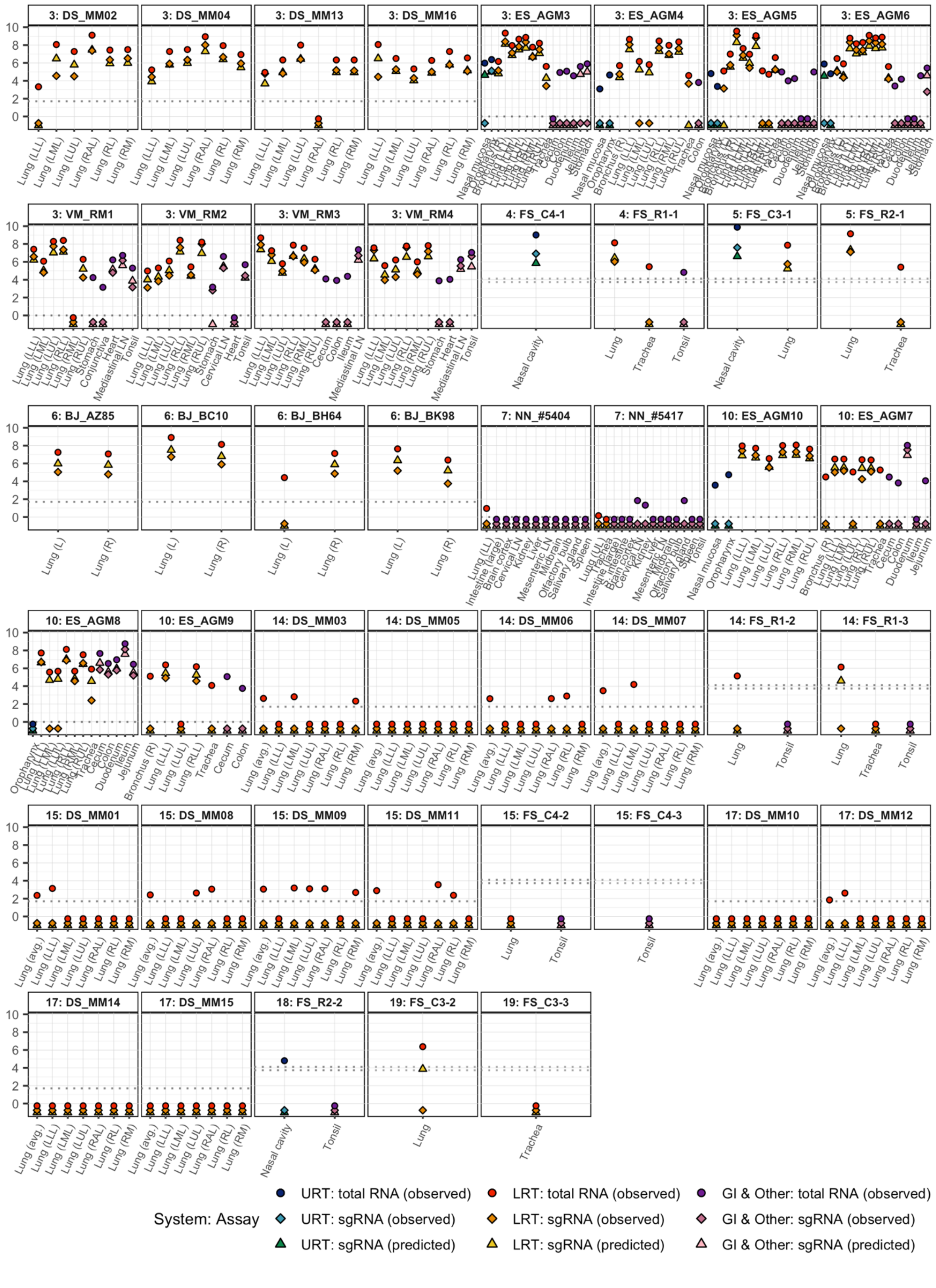


**Figure S6. Individual viral loads for invasive samples, including sgRNA predictions generated by the best sgRNA model.** Each panel corresponds with one individual, indicated with text in the panel (day post infection: individual). Each point presents the total RNA (circle), observed sgRNA (diamond), and predicted sgRNA (triangle) values. All samples observed or predicted to fall below the limit of detection are plotted below 0 at set values for visual clarity (totRNA: -0.5, observed sgRNA: -0.75, predicted sgRNA: -1). When available, the limits of detection (LOD) or quantification (LOQ) for PCR assays are plotted as dotted lines in the assay-specific color. When both the LOD and LOQ were available, only the LOD is plotted. In instances where the total RNA and sgRNA assay LOD are equal, only the sgRNA line is visible. No instances exist in this dataset where the LOD or LOQ is only available for one RNA type.

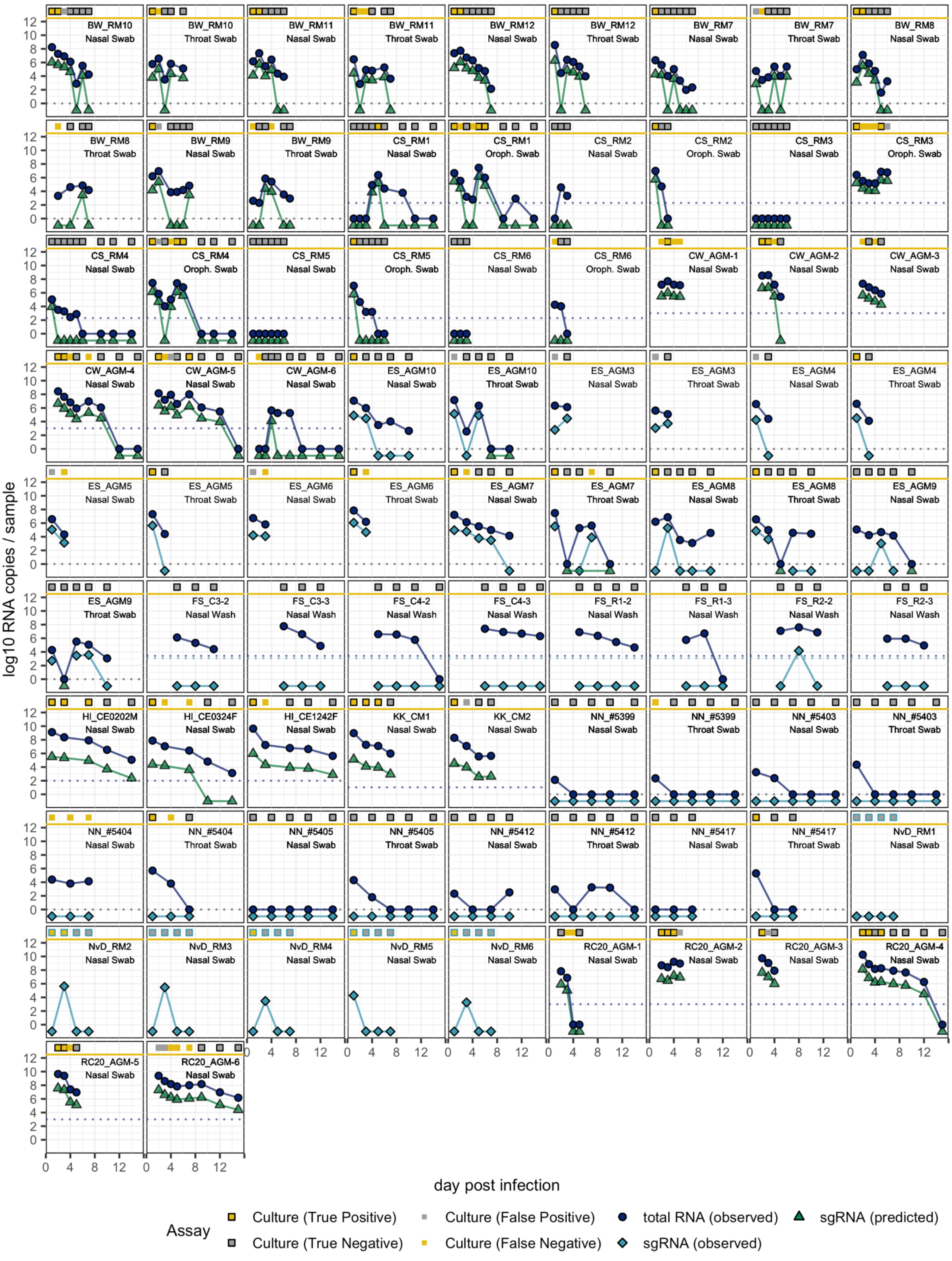

BW_RM10 Nasal Swab
BW_RM10 Throat Swab
BW_RM11 Nasal Swab
BW_RM11 Throat Swab
BW_RM12 Nasal Swab
BW_RM12 Throat Swab
BW_RM7 Nasal Swab
BW_RM7 Throat Swab
BW_RM8 Nasal Swab
BW_RM8 Throat Swab
BW_RM9 Nasal Swab
BW_RM9 Throat Swab
CS_RM1 Nasal Swab
CS_RM1 Oroph. Swab
CS_RM2 Nasal Swab
CS_RM2 Oroph. Swab
CS_RM3 Nasal Swab
CS_RM3 Oroph. Swab
CS_RM4 Nasal Swab
CS_RM4 Oroph. Swab
CS_RM5 Nasal Swab
CS_RM5 Oroph. Swab
CS_RM6 Nasal Swab
CS_RM6 Oroph. Swab
CW_AGM-1 Nasal Swab
CW_AGM-2 Nasal Swab
CW_AGM-3 Nasal Swab
CW_AGM-4 Nasal Swab
CW_AGM-5 Nasal Swab
CW_AGM-6 Nasal Swab
ES_AGM10 Nasal Swab
ES_AGM10 Throat Swab
ES_AGM3 Nasal Swab
ES_AGM3 Throat Swab
ES_AGM4 Nasal Swab
ES_AGM4 Throat Swab
ES_AGM5 Nasal Swab
ES_AGM5 Throat Swab
ES_AGM6 Nasal Swab
ES_AGM6 Throat Swab
ES_AGM7 Nasal Swab
ES_AGM7 Throat Swab
ES_AGM8 Nasal Swab
ES_AGM8 Throat Swab
ES_AGM9 Nasal Swab
ES_AGM9 Throat Swab
FS_C3-2 Nasal Wash
FS_C3-3 Nasal Wash
FS_C4-2 Nasal Wash
FS_C4-3 Nasal Wash
FS_R1-2 Nasal Wash
FS_R1-3 Nasal Wash
FS_R2-2 Nasal Wash
FS_R2-3 Nasal Wash
HI_CE0202M Nasal Swab
HI_CE0324F Nasal Swab
HI_CE1242F Nasal Swab
KK_CM1 Nasal Swab
KK_CM2 Nasal Swab
NN_#5399 Nasal Swab
NN_#5399 Throat Swab
NN_#5403 Nasal Swab
NN_#5403 Throat Swab
NN_#5404 Nasal Swab
NN_#5404 Throat Swab
NN_#5405 Nasal Swab
NN_#5405 Throat Swab
NN_#5412 Nasal Swab
NN_#5412 Throat Swab
NN_#5417 Nasal Swab
NN_#5417 Throat Swab
NvD_RM1 Nasal Swab
NvD_RM2 Nasal Swab
NvD_RM3 Nasal Swab
NvD_RM4 Nasal Swab
NvD_RM5 Nasal Swab
NvD_RM6 Nasal Swab
RC20_AGM-1 Nasal Swab
RC20_AGM-2 Nasal Swab
RC20_AGM-3 Nasal Swab
RC20_AGM-4 Nasal Swab
RC20_AGM-5 Nasal Swab
RC20_AGM-6 Nasal Swab
log10 RNA copies / sample
day post infection
Assay
Culture (True Positive)
Culture (False Positive)
total RNA (observed)
sgRNA (predicted)
Culture (True Negative)
Culture (False Negative)
sgRNA (observed)

**Figure S7. Individual culture trajectories in the upper respiratory tract.** Each panel corresponds with one individual and one non-invasive sample type, indicated in the top right of each panel. Only individuals with culture results for at least two days post infection are plotted. Culture data are plotted as squares above the yellow line at 10 log10 copies. Yellow squares are culture positive samples, while grey squares are culture negative. Squares outlined in black are correct predictions, squares with no outline are incorrect predictions. We did not generate predictions for the culture samples outlined in blue, as they do not have available totRNA results. We also plot observed total RNA values (circle) and observed sgRNA values (diamond), otherwise we plot predicted median sgRNA values generated by our best sgRNA model (triangle). Some individuals were sampled from multiple locations in the upper respiratory tract, in which case they are plotted as neighboring panels. All samples observed or predicted to fall below the limit of detection are plotted below 0 at set values for visual clarity (totRNA: 0, sgRNA: -1). When available, the limits of detection (LOD) or quantification (LOQ) for PCR assays are plotted as dotted lines in the assay-specific color. When both the LOD and LOQ were available, only the LOD is plotted. In instances where the total RNA and sgRNA assay LOD are equal, only the sgRNA line is visible. No instances exist in this dataset where the LOD or LOQ is only available for one RNA type. Individuals from one study currently cannot be included in this figure, though they will likely be incorporated in future versions.

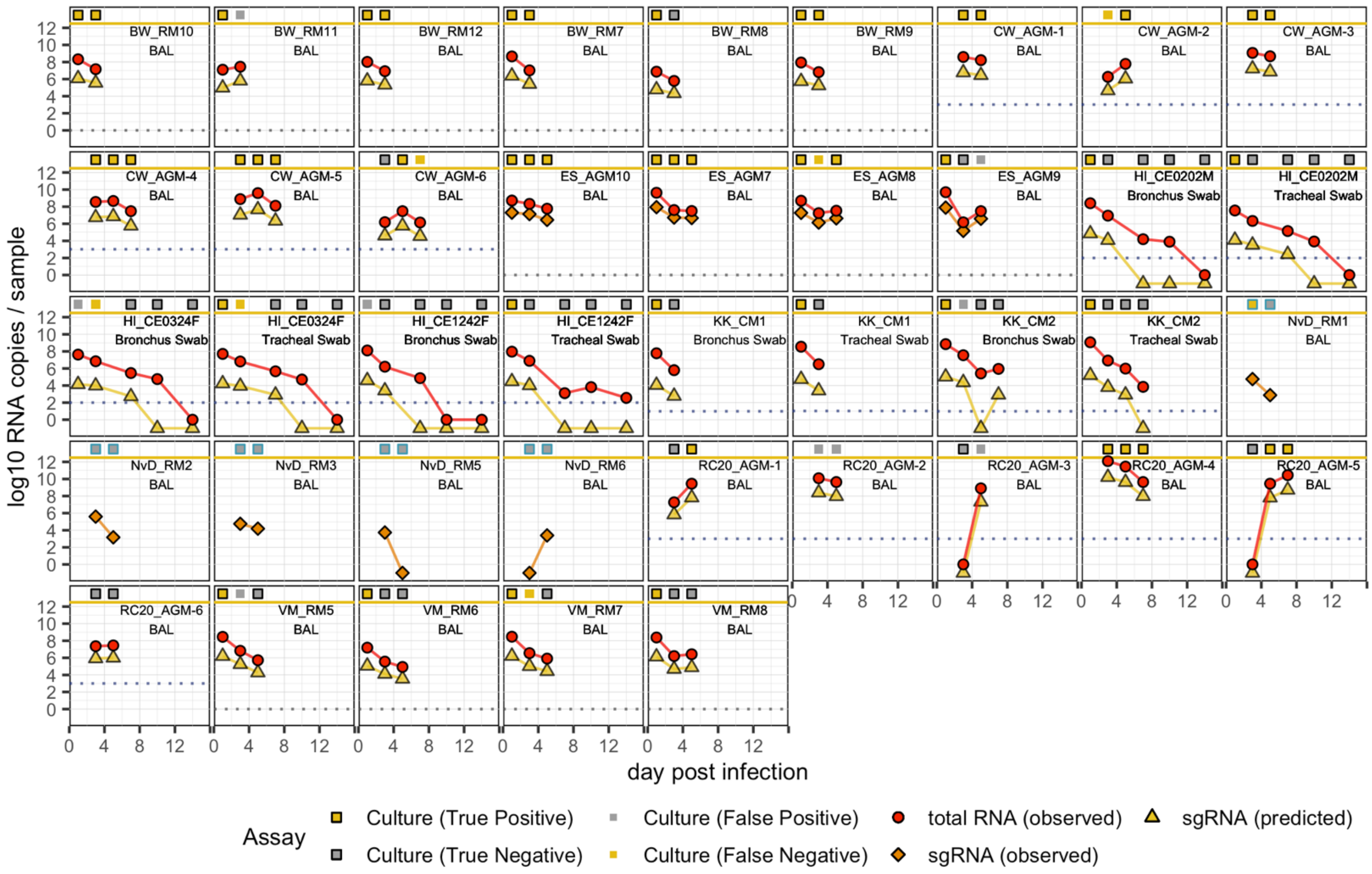


**Figure S8. Individual culture trajectories in the lower respiratory tract.** Each panel corresponds with one individual and one non-invasive sample type, indicated in the top right of each panel. Only individuals with culture results for at least two days post infection are plotted. Culture data are plotted as squares above the yellow line at 10 log10 copies. Yellow squares are culture positive samples, while grey squares are culture negative. Squares outlined in black are correct predictions, squares with no outline are incorrect predictions. We did not generate predictions for the culture samples outlined in blue, as they do not have available totRNA results. We also plot observed total RNA values (circle) and observed sgRNA values (diamond) when available, otherwise we plot predicted median sgRNA values generated by our best sgRNA model (triangle). Some individuals were sampled from multiple locations in the lower respiratory tract, in which case they are plotted as neighboring panels. All samples observed or predicted to fall below the limit of detection are plotted below 0 at set values for visual clarity (totRNA: 0, sgRNA: -1). When available, the limits of detection (LOD) or quantification (LOQ) for PCR assays are plotted as dotted lines in the assay-specific color. When both the LOD and LOQ were available, only the LOD is plotted. In instances where the total RNA and sgRNA assay LOD are equal, only the sgRNA line is visible. No instances exist in this dataset where the LOD or LOQ is only available for one RNA type.

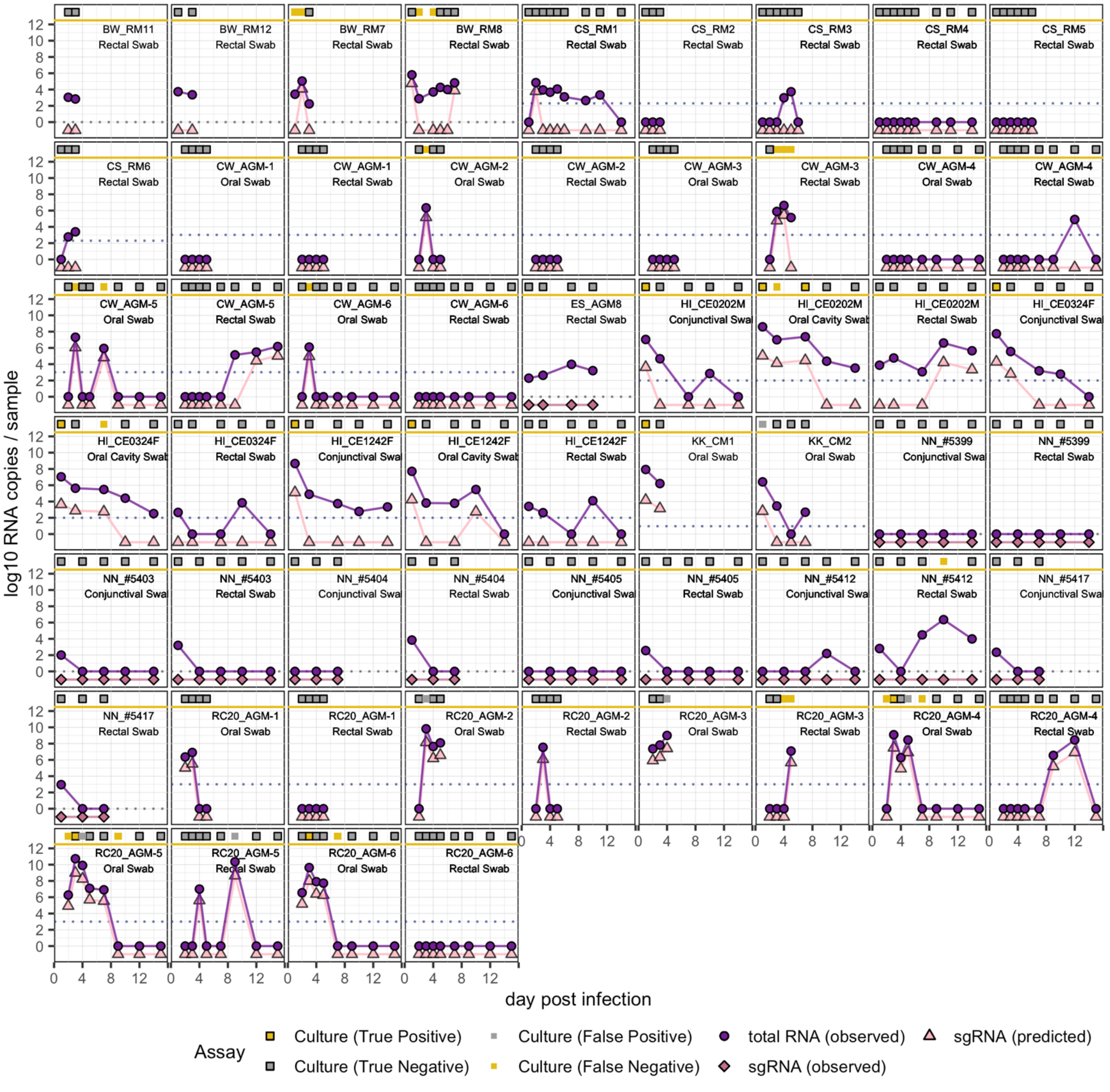


**Figure S9. Individual culture trajectories in the gastrointestinal and other systems.** Each panel corresponds with one individual and one non-invasive sample type, indicated in the top right of each panel. Only individuals with culture results for at least two days post infection are plotted. Culture data are plotted as squares above the yellow line at 10 log10 copies. Yellow squares are culture positive samples, while grey squares are culture negative. Squares outlined in black are correct predictions, squares with no outline are incorrect predictions. We also plot observed total RNA values (circle) and observed sgRNA values (diamond) when available, otherwise we plot predicted median sgRNA values generated by our best sgRNA model (triangle). Some individuals were sampled from multiple locations, in which case they are plotted as neighboring panels. All samples observed or predicted to fall below the limit of detection are plotted below 0 at set values for visual clarity (totRNA: 0, sgRNA: -1). When available, the limits of detection (LOD) or quantification (LOQ) for PCR assays are plotted as dotted lines in the assay-specific color. When both the LOD and LOQ were available, only the LOD is plotted. In instances where the total RNA and sgRNA assay LOD are equal, only the sgRNA line is visible. No instances exist in this dataset where the LOD or LOQ is only available for one RNA type. Individuals from one study currently cannot be included in this figure, though they will likely be incorporated in future versions.

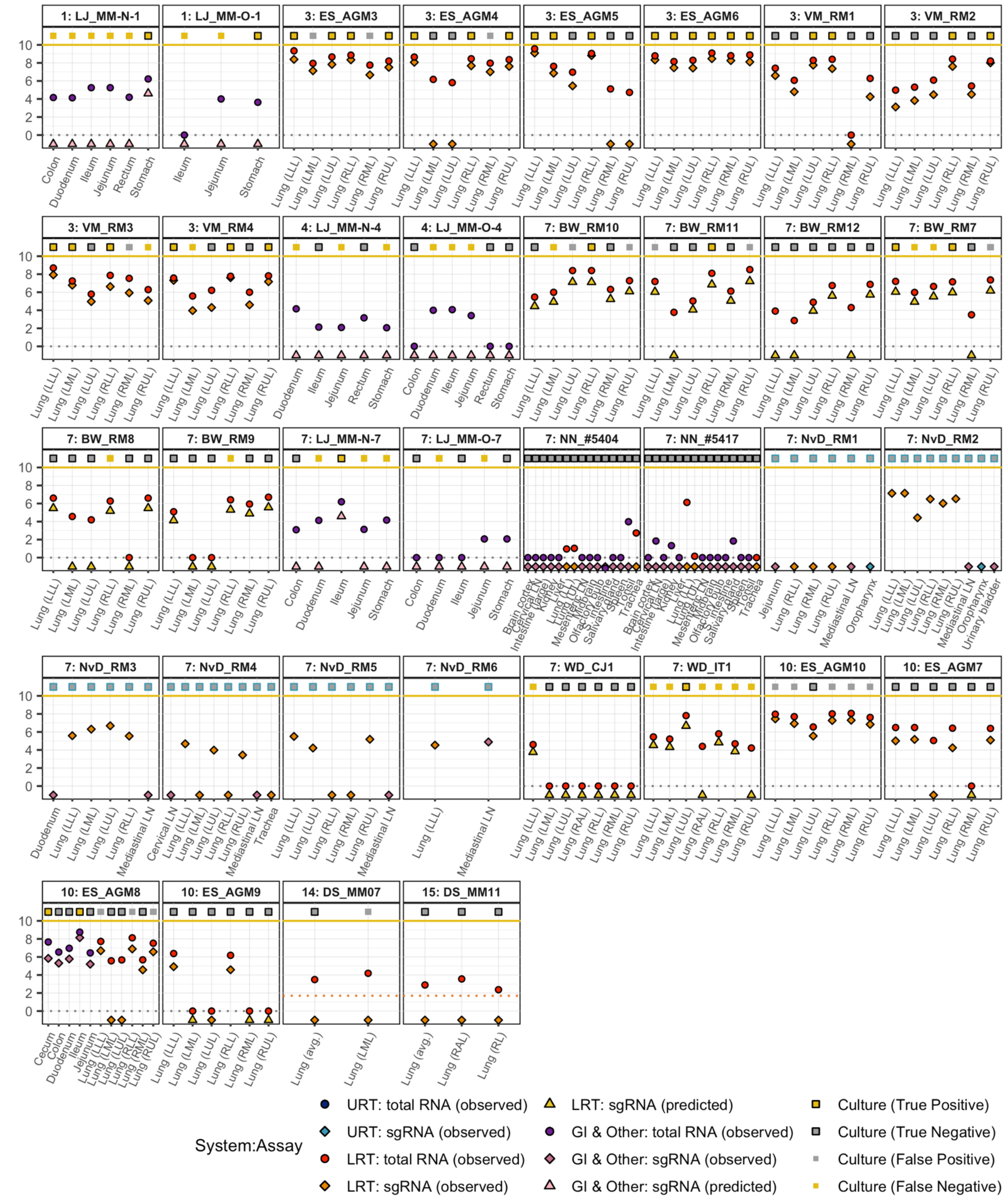


**Figure S10. Individual culture data for invasive samples.** Each panel corresponds with one individual, indicated with text in the panel (day post infection: individual). Culture data are plotted as squares above the yellow line at 10 log10 copies. Yellow squares are culture positive samples, while grey squares are culture negative. Squares outlined in black are correct predictions, squares with no outline are incorrect predictions. We did not generate predictions for the culture samples outlined in blue, as they do not have available totRNA results. We also plot the observed total RNA (circle) and observed sgRNA (diamond) values when available, otherwise we plot predicted median sgRNA values generated by our best sgRNA model (triangle). Color corresponds to the organ system from which the tissue was obtained (URT, upper respiratory tract; LRT, lower respiratory tract; GI & Other, gastrointestinal and other systems). All samples observed or predicted to fall below the limit of detection are plotted below 0 at set values for visual clarity (totRNA: 0, sgRNA: -1). When available, the limits of detection (LOD) or quantification (LOQ) for PCR assays are plotted as dotted lines in the assay-specific color. When both the LOD and LOQ were available, only the LOD is plotted. In instances where the total RNA and sgRNA assay LOD are equal, only the sgRNA line is visible. No instances exist in this dataset where the LOD or LOQ is only available for one RNA type.

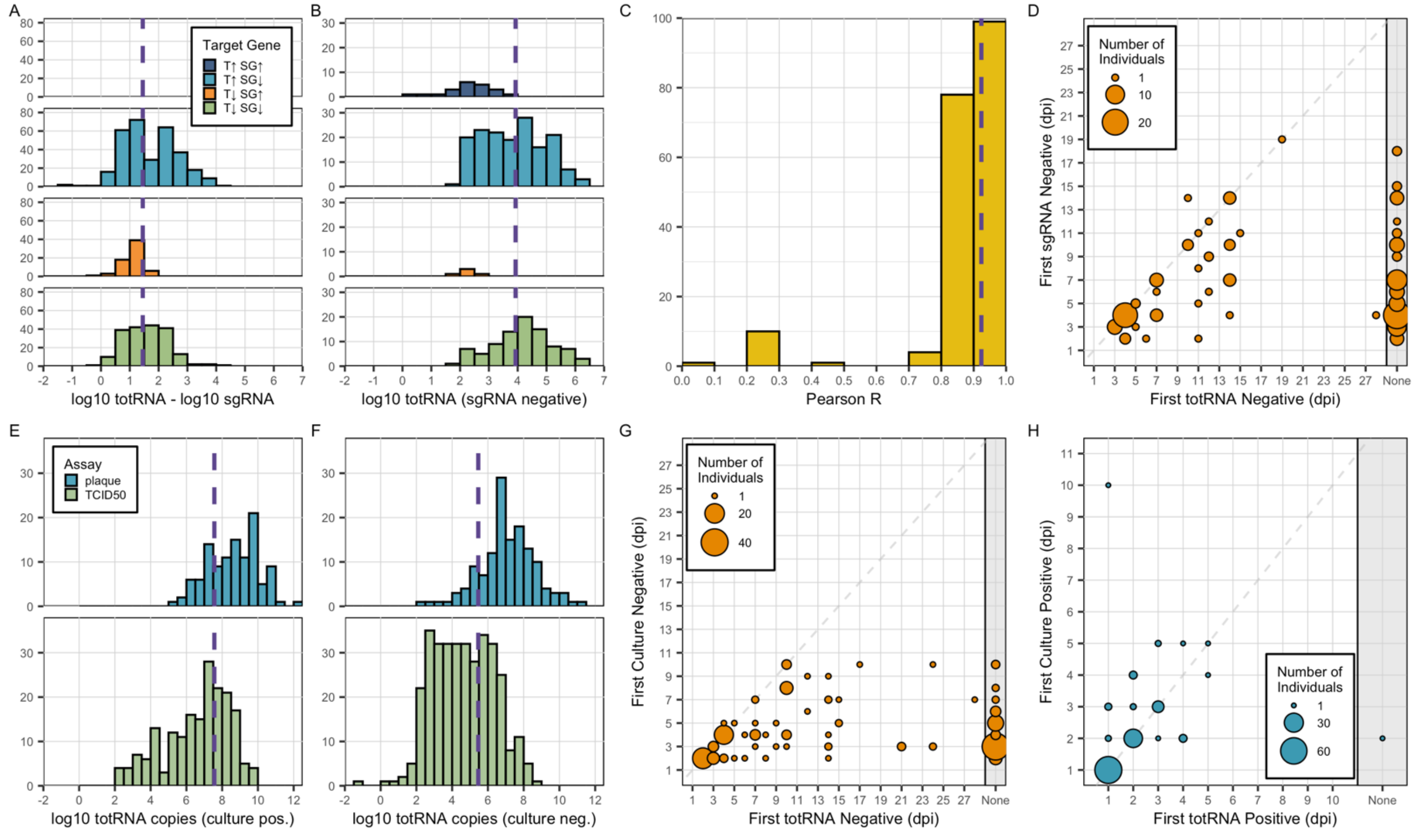


**Figure S11. Statistics relating PCR and culture results**. (A) Difference between total RNA and sgRNA copy numbers when both are detectable, stratified by target gene predictor with the following acronyms: "T↑SG↑": totRNA-high/sgRNA-high; "T↓SG↑": totRNA-low/sgRNA-high; "T↑SG↓": totRNA-high/sgRNA-low; "T↓SG↓": totRNA-high/sgRNA-low. No totRNA-high/sgRNA-high data was available for this investigation. (B) Total RNA copy numbers for all sgRNA negative samples, stratified by target gene as in (A). (C) Pearson correlation coefficients between total RNA and sgRNA copy numbers when both are detectable, for all individual-sample trajectories with at least three sampling days where both were positive. (D) Comparison of the timing of the first negative results from total RNA and sgRNA assays for each available individual-sample trajectory. (E) Total RNA copy numbers (when detectable) for all culture positive samples, stratified by culture assay type. (F) Total RNA copy numbers (when detectable) for all culture negative samples, stratified by culture assay type as in (E). (G) Comparison of the timing of the first negative results from total RNA and culture assays for each available individual-sample trajectory. (H) Comparison of the timing of the first positive results from total RNA and culture assays for each individual-sample trajectory. For panels (A), (B), (C), (E), and (F), the purple dashed line indicates the median for the full distribution (i.e., not stratified by assay or target gene). For panels (D), (G), and (H), the size of each circle indicates the number of individuals with the indicated observation. Individuals in the 'None' column were never negative (D, G) or positive (H) for total RNA. Individuals that were never sgRNA negative (D), culture negative (G), or culture positive (H) are not plotted.

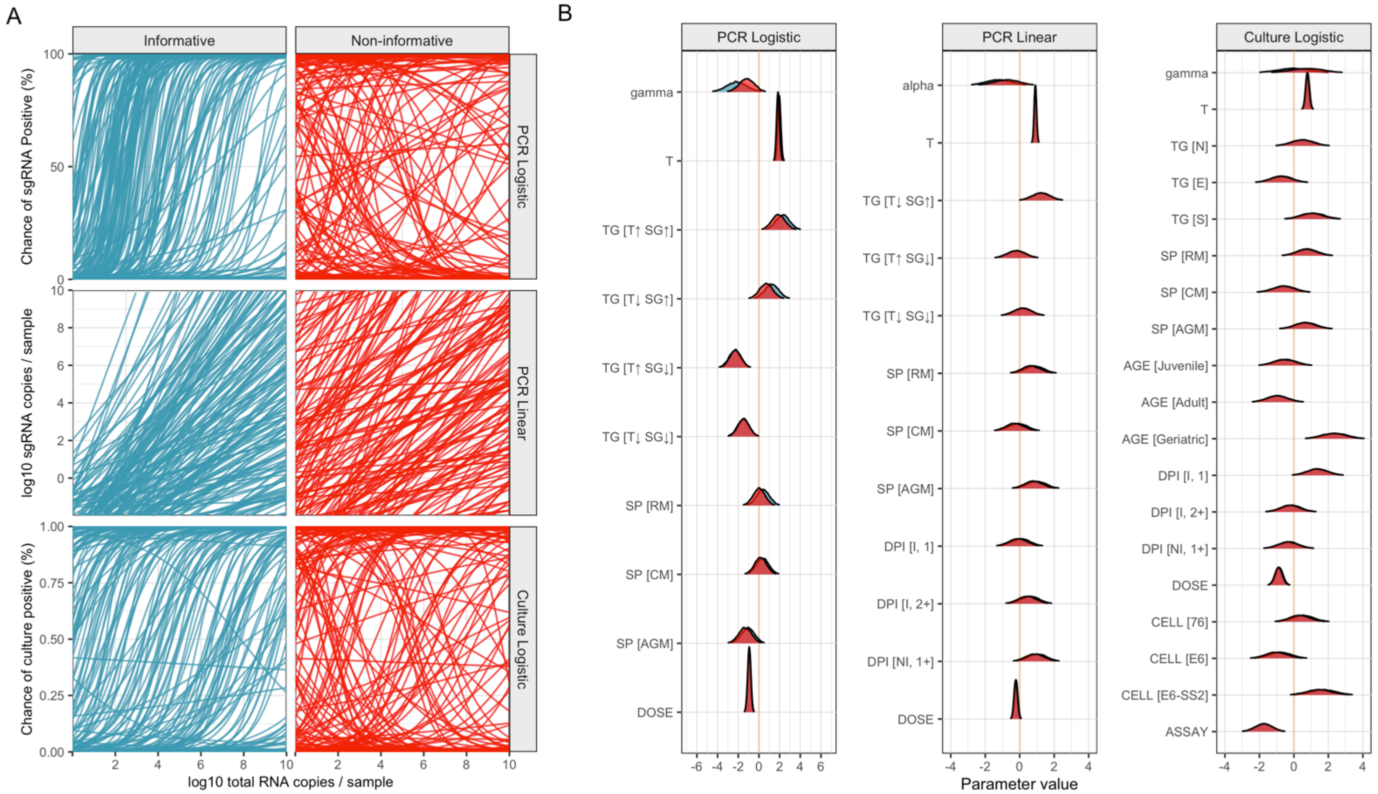


**Figure S12. Sensitivity analyses comparing informative (blue) and non-informative (red) priors.** (A) Each line presents an expected model fit generated by sampling the indicated prior distributions. Informative priors are outlined in the **Methods** and **Supplementary Methods**. All parameters were given a $N(0,1)$ prior for all non-informative investigations. Informative priors much better represent *a priori* understanding of the relationships between total RNA copy numbers and both sgRNA and culture outcomes. (B) Each panel compares the final parameter estimates obtained for the corresponding best model using the different prior types (red: non-informative; blue: informative), where each row is a distinct parameter. Acronyms are as described in **Figures 3, 4, and 5**. Note that in many instances parameters estimates are almost perfectly overlapping, so only the non-informative (red) priors are visible.

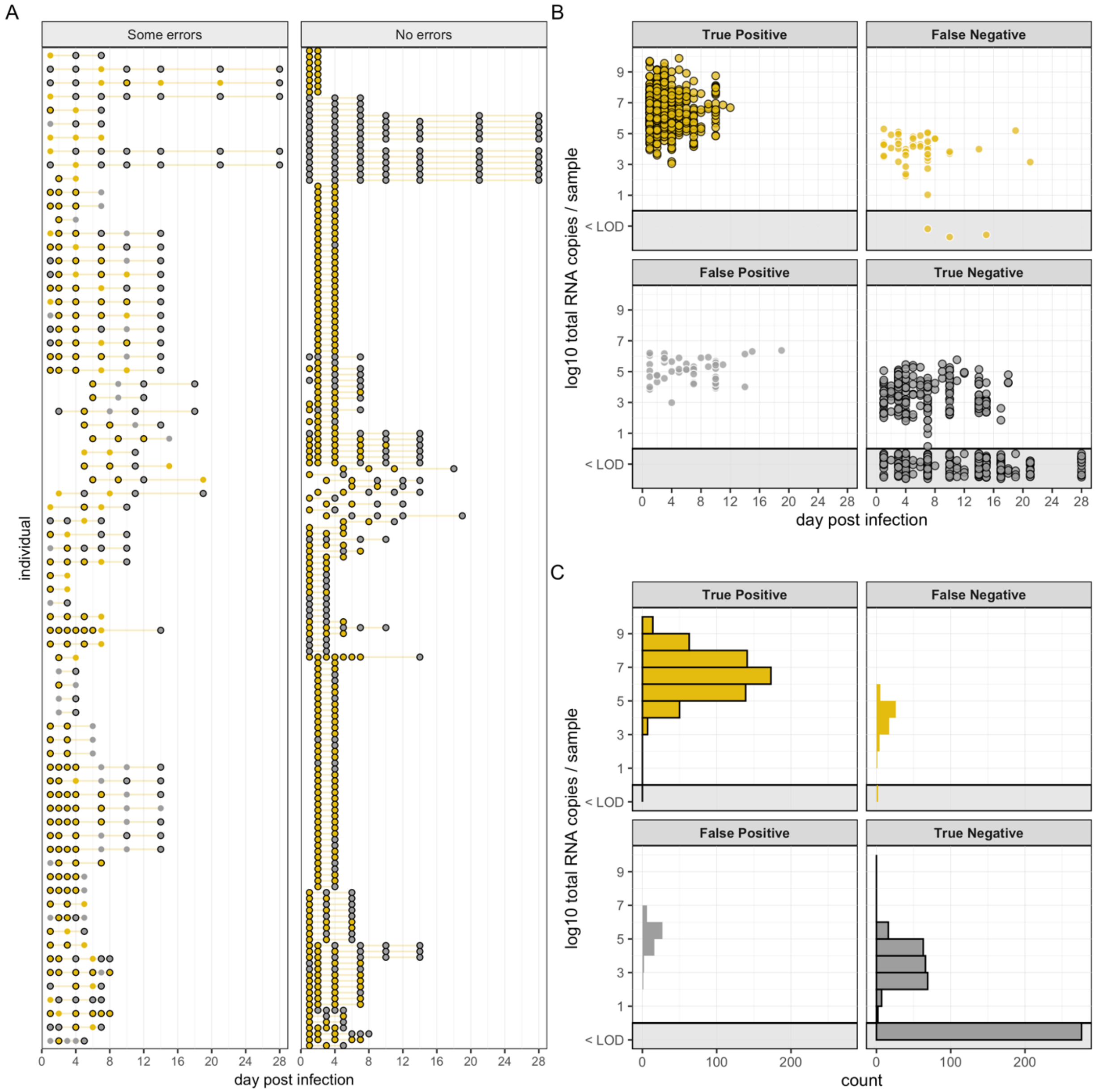


**Figure S13. Error analysis for the best sgRNA model.** (A) Individual-specific sgRNA trajectories, where each row presents one individual. These are stratified by whether the model misclassifies any samples for that individual ("Some errors") or whether the model makes no misclassifications ("No errors"). In both (A) and (B), yellow circles indicate positive samples and grey indicates negative samples. Circles with a black outline correspond with correctly classified samples, while no outline indicates incorrectly classified samples. (B) Scatterplot of all samples with sgRNA results, stratified by the elements of a confusion matrix and colored as in (A). The x-axis tracks the day post infection and the y-axis plots log10 total RNA copy numbers. Samples in the grey shaded region along the bottom present all samples where total RNA was undetectable. (C) Histograms of all samples grouped by the elements of a confusion matrix, where log10 total RNA copy numbers per sample is plotted on the y-axis. Bins located in the grey shaded region along the bottom (labelled "<LOD") include all totRNA-negative samples.

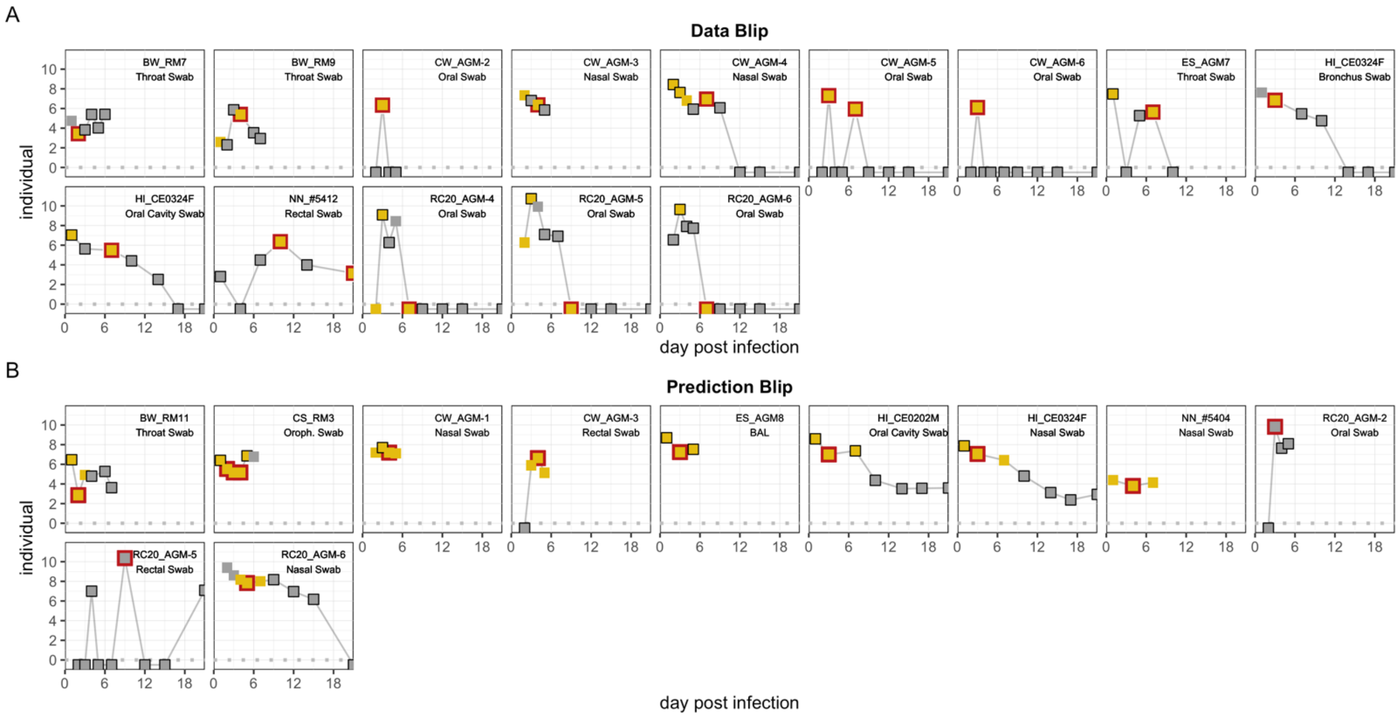


**Figure S14. Viral load and culture trajectories for individuals with data blip (A) or prediction blip (B) error types.** Panel-specific errors are indicated with red outlines. All other samples with prediction errors have no outline. Yellow squares indicate known culture positive samples, while grey squares indicate known culture negative samples. Text in the upper right corner of each panel indicates the ID name and sample type of the individual from whom the data was derived. All totRNA-negative samples are plotted below the grey dashed line at zero. Note that individual NN_#5412 has an additional (true negative) sample available on a later day post infection, which is not shown for visual clarity. Six trajectories from one study currently cannot be included in this figure, though they will likely be incorporated in future versions.

#### Supplementary Tables

| Article | N | Species | Sex | Age class | Exposure route | Exposure dose | Viral isolate | Sample type | Sample location | Sample time | PCR target gene *totRNA* | PCR target gene *sgRNA* | Culture cell line |
|---|---|---|---|---|---|---|---|---|---|---|---|---|---|
| Baum et al. 2020 | 43 (4) | RM | F, M | U | IT, IN | 6.02 | USA/WA1/2020 | NI | 1, 2, 3 | URT, Other | N | E (3) | -- |
| | 48 (6) | RM | F, M | A | IT, IN | 5.04 | USA/WA1/2020 | NI | 1, 2 | URT, LRT | N | E (3) | -- |
| Chandrashekar et al. 2020 | 12 (3) | RM | U | A | IT, IN | 6.04 | USA/WA1/2020 | NI | 1, 2 | URT | N | E (3) | -- |
| | 12 (3) | RM | U | A | IT, IN | 5.04 | USA/WA1/2020 | NI | 1, 2 | URT | N | E (3) | -- |
| | 12 (3) | RM | U | A | IT, IN | 4.04 | USA/WA1/2020 | NI | 1, 2 | URT | N | E (3) | -- |
| Corbett et al. 2020 | 50 (8) | RM | F, M | J, A | IT, IN | 5.88 | USA/WA1/2020 | NI | 1, 2 | URT, LRT | N | E (3) | -- |
| Cross et al. 2020 | 124 (6) | AGM | F | A | IN | 6.45 | ITA/INMI1/2020 | NI | 2, 3 | URT, LRT, GI, Other | N | -- | Vero E6 |
| Dagotto et al. 2021 | 16 (4) | RM | U | A | IT, IN | 4.04 | USA/WA1/2020 | NI | 1, 2 | LRT | N, E | E (3, 4) | -- |
| Deng et al. 2020 | 7 (1) | RM | M | J | IT | 5.85* | CHN/WH-09/2020 | I | 3 | LRT | E | -- | Vero E6† |
| | 7 (1) | RM | M | J | OC | 5.85* | CHN/WH-09/2020 | I | 3 | LRT | E | -- | Vero E6† |
| Gabitzsch et al. 2021 | 24 (2) | RM | F, M | J | IT, IN | 5.85* | USA/WA1/2020 | NI | 1, 2 | URT, LRT | N | E (3) | -- |
| Ishigaki et al. 2021 | 144 (3) | CM | F, M | A | IT, IN, OR, OC | 6.19* | JPN/WK-521/2020 | NI | 1, 2, 3 | URT, LRT, GI, Other | N | -- | Vero E6† |
| Jiao et al. 2021 | 16 (3) | RM | M | U | IN | 7 | CHN/U | I | 3 | GI | N | -- | Vero E6† |
| | 14 (3) | RM | M | U | IG | 7 | CHN/U | I | 1, 2, 3 | GI | N | -- | Vero E6† |
| Johnston et al. 2020 | 60 (4) | RM | F, M | A | AE | 4.46 | USA/WA1/2020 | NI | 2, 3 | URT, GI | N | -- | Vero 76 |
| | 45 (3) | AGM | F, M | A | AE | 4.58 | USA/WA1/2020 | NI | 2, 3 | URT, GI | N | -- | Vero 76 |
| | 60 (4) | CM | F, M | J, A | AE | 4.69 | USA/WA1/2020 | NI | 2, 3 | URT, GI | N | -- | Vero 76 |
| Jones et al. 2021 | 44 (4) | RM | F | A | IT, IN | 5.04 | USA/WA1/2020 | I, NI | 1, 2, 3 | URT, LRT | N | E (3) | -- |
| Kobiyama et al. 2021 | 26 (2) | CM | F | A | IT, IN, OR, OC | 7.3 | U/U | NI | 1, 2 | URT, LRT, Other | N | -- | Vero E6/TMPRSS2† |
| Li et al. 2021 | 148 (16) | CM | F, M | J, A | IT, IN | 5 | USA/WA1/2020 | NI | 2 | URT, LRT | E | E, N (4, 2) | -- |
| Munster et al. 2020 | 53 (8) | RM | F, M | J, A | IT, IN, OR, OC | 6.26* | USA/WA1/2020 | I, NI | 1, 2, 3 | LRT, GI, Other | E | ORF7 (4) | Vero E6 |
| Nagata et al. 2021 | 165 (6) | CM | F | A | IT, IN, OC | 7.42* | JPN/WK-521/2020 | I, NI | 1, 2, 3 | URT, LRT, GI, Other | N | N (1) | Vero E6/TMPRSS2† |
| Patel et al. 2021 | 65 (5) | RM | F, M | J | IT, IN | 4.04 | USA/WA1/2020 | NI | 1, 2 | URT, LRT | N | E (3) | -- |
| Salguero et al. 2021 | 63 (6) | RM | F, M | J | IT, IN | 6.7 | AUS/VIC01/2020 | I, NI | 1, 2, 3 | URT, LRT, Other | N | E (3) | Vero E6 |
| | 58 (6) | CM | F, M | J | IT, IN | 6.7 | AUS/VIC01/2020 | I, NI | 1, 2, 3 | URT, LRT, Other | N | E (3) | Vero E6 |
| Shan et al. 2020 | 108 (6) | RM | F, M | A | IT | 6.69* | CHN/WIV04/2019 | NI | 3 | URT, GI | S | -- | Vero E6† |
| Singh et al. 2020 | 108 (16) | RM | F, M | G, J | IT, IN, OC | 6.02 | USA/WA1/2020 | I | 3 | LRT | N | E (3) | Vero E6 |
| Speranza et al. 2020 | 194 (10) | AGM | F, M | A | IT, IN, OR, OC | 6.26* | USA/WA1/2020 | I, NI | 1, 2, 3 | URT, LRT, GI | E | E (4) | Vero E6 |
| van Doremalen et al. 2020 | 72 (6) | RM | U | J | IT, IN, OR, OC | 6.26* | USA/WA1/2020 | I, NI | 1, 2, 3 | URT, LRT, GI, Other | -- | E | Vero E6† |
| Williamson et al. 2020 | 135 (6) | RM | F, M | J | IT, IN, OR, OC | 6.26* | USA/WA1/2020 | I, NI | 1, 2, 3 | URT, LRT, GI | E | -- | Vero E6† |
| Woolsey et al. 2020 | 132 (6) | AGM | F, M | A | IT, IN | 5.66 | ITA/INMI1/2020 | NI | 2, 3 | URT, LRT, GI, Other | N | -- | Vero E6 |

| | | | | | | | | | | | | |
|---|---|---|---|---|---|---|---|---|---|---|---|---|
| Yu et al. 2020 | 102 (10) | RM | U | A | IT, IN | 4.04 | U/U | NI | 1, 2 | URT, LRT | N | E (3) | -- |

**Table S1. Summary of articles included in the dataset.** Multiple rows for an individual article are included when the study involved multiple species and/or multiple exposure doses. In all columns, U indicates the detail is unknown. Sample sizes (N) are presented in the following format: number of available datapoints (number of individuals). Species abbreviations are as follows: RM, rhesus macaque; CM, cynomolgus macaque; AGM, African green monkey. Age class presents the standardized assignments according to our protocol (**Supplementary Methods**), and the abbreviations are: J, juvenile; A, adult; G, geriatric. Individuals inoculated via multiple routes are indicated by exposure routes joined by commas, where the abbreviations are: AE, aerosol; IT, intratracheal; IN, intranasal; IG, intragastric; OC, ocular; OR, oral. Exposure dose is presented as log10 plaque forming units, and an adjoining * indicates the dose was originally reported as TCID50, so those values were converted using the standard method described in the **Supplementary Methods**. NI indicates non-invasive sample types (i.e., swabs, biofluids, BAL), while I indicates invasive tissue samples obtained at necropsy. Sample location distinguishes between the following systems: URT, upper respiratory tract; LRT, lower respiratory tract; GI, gastrointestinal tract; and Other, all other locations. Sample time presents the days post infection with available samples according to our DPI predictor, where 1: 1 dpi, inoculated tissues, 2: 2+ dpi, inoculated tissues, 3: any dpi, non-inoculated tissue (further categorization information is in **Table S9**). PCR target genes are stratified by total RNA (totRNA) and sgRNA. The level of the target gene predictor for the sgRNA model follows the sgRNA gene in parentheses: (1) totRNA-high/sgRNA-high, (2) totRNA-low/sgRNA-high, (3) totRNA-high/sgRNA-low, and (4) totRNA-low/sgRNA-low. The cell lines used for culture are indicated when available, with SS2 as an abbreviation for TMPRSS2. An adjoining † indicates the use of a TCID50 assay, while no symbol indicates a plaque assay.

| Model | Predictors | *Cross-validation* ELPD Difference (SE) | *Cross-validation* ELPD (SE) | *PSIS-LOO Approximation* ELPD Difference (SE) | *PSIS-LOO Approximation* ELPD (SE) | *Prediction* MCC | *Prediction* % correctly predicted train | *Prediction* % correctly predicted test |
|---|---|---|---|---|---|---|---|---|
| **l1** | | **-73.31 (10.63)** | **-311.56 (17.36)** | **-71.13 (10.76)** | **-311.33 (17.29)** | **0.75** | **87.75** | **87.69** |
| l2.1 | DOSE | -60.78 (10.02) | -299.03 (19.7) | -58.16 (10.15) | -298.36 (19.46) | 0.78 | 89.35 | 89.28 |
| l2.2 | ST | -58.37 (9.44) | -296.62 (18.51) | -56.4 (9.65) | -296.6 (18.45) | 0.79 | 89.31 | 89.45 |
| l2.3 | SP | -45.45 (9.79) | -283.71 (17.24) | -43.66 (9.94) | -283.86 (17.26) | 0.79 | 89.93 | 89.78 |
| l2.4 | AGE | -71.06 (10.53) | -309.31 (17.43) | -69.38 (10.65) | -309.58 (17.39) | 0.76 | 88.63 | 88.27 |
| l2.5 | SEX | -72.66 (10.58) | -310.91 (17.57) | -73.37 (10.75) | -313.57 (17.64) | 0.75 | 87.89 | 87.52 |
| l2.6 | DPI | -67.27 (10.05) | -305.52 (18.06) | -64.8 (10.2) | -305 (17.98) | 0.76 | 88.28 | 88.19 |
| l2.7 | TG | -40.19 (9.13) | -278.44 (16.42) | -37.97 (9.2) | -278.17 (16.39) | 0.79 | 89.61 | 89.61 |
| l3.1 | TG + DOSE | -10.98 (4.22) | -249.23 (17.36) | -8.53 (4.25) | -248.73 (17.25) | 0.81 | 91.02 | 90.79 |
| l3.2 | TG + ST | -32.17 (7.81) | -270.42 (17.23) | -30.17 (7.96) | -270.37 (17.15) | 0.8 | 90.27 | 89.95 |
| l3.3 | TG + SP | -24.62 (7.97) | -262.87 (16.06) | -22.69 (8.03) | -262.89 (16.07) | 0.79 | 89.86 | 89.7 |
| l3.4 | TG + AGE | -33.67 (8.07) | -271.92 (16.34) | -32.3 (8.09) | -272.5 (16.33) | 0.78 | 89.61 | 89.2 |
| l3.5 | TG + SEX | -43.33 (9.22) | -281.58 (16.7) | -43.32 (9.48) | -283.52 (16.88) | 0.78 | 89.87 | 88.86 |
| l3.6 | TG + DPI | -37.05 (8.46) | -275.3 (16.91) | -34.24 (8.52) | -274.44 (16.84) | 0.79 | 89.86 | 89.7 |
| l4.1 | TG + DOSE + ST | -9.37 (3.78) | -247.62 (17.99) | -7.31 (3.84) | -247.51 (17.84) | 0.82 | 91.4 | 90.95 |
| **l4.2** | **TG + DOSE + SP** | **-5.98 (3.58)** | **-244.23 (17.2)** | **-4.19 (3.61)** | **-244.39 (17.16)** | **0.82** | **91.41** | **91.12** |
| l4.3 | TG + DOSE + AGE | -10.94 (3.87) | -249.19 (17.45) | -9.11 (3.9) | -249.31 (17.38) | 0.81 | 91.1 | 90.79 |
| l4.4 | TG + DOSE + SEX | -12.48 (4.26) | -250.73 (17.42) | -10.3 (4.24) | -250.5 (17.35) | 0.81 | 90.91 | 90.7 |
| l4.5 | TG + DOSE + DPI | -12.29 (4.2) | -250.54 (17.7) | -9.73 (4.21) | -249.93 (17.58) | 0.82 | 91.09 | 91.04 |
| l5.1 | TG + DOSE + SP + ST | -4.58 (3) | -242.83 (17.74) | -3.05 (3.08) | -243.25 (17.64) | 0.82 | 91.3 | 91.12 |
| l5.2 | TG + DOSE + SP + AGE | -1.52 (2.02) | -239.77 (17.25) | -1.15 (1.95) | -241.35 (17.28) | 0.83 | 91.62 | 91.46 |
| l5.3 | TG + DOSE + SP + SEX | -7.43 (3.62) | -245.68 (17.28) | -5.92 (3.62) | -246.12 (17.24) | 0.82 | 91.36 | 91.21 |
| l5.4 | TG + DOSE + SP + DPI | -6.66 (3.42) | -244.91 (17.51) | -4.74 (3.42) | -244.94 (17.41) | 0.82 | 91.14 | 91.12 |
| l6.1 | TG + DOSE + SP + AGE + ST | 0 (0) | -238.25 (17.79) | 0 (0) | -240.2 (17.78) | 0.83 | 91.69 | 91.46 |
| l6.2 | TG + DOSE + SP + AGE + SEX | -2.89 (2.02) | -241.14 (17.31) | -2.32 (1.94) | -242.52 (17.36) | 0.83 | 91.69 | 91.46 |
| l6.3 | TG + DOSE + SP + AGE + DPI | -1.51 (1.88) | -239.76 (17.61) | -1.86 (1.83) | -242.05 (17.71) | 0.82 | 91.35 | 91.04 |
| l7.1 | TG + DOSE + SP + AGE + ST + SEX | -1.41 (0.56) | -239.66 (17.87) | -1.28 (0.41) | -241.48 (17.85) | 0.83 | 91.71 | 91.46 |
| l7.2 | TG + DOSE + SP + AGE + ST + DPI | -1.23 (0.99) | -239.48 (17.96) | -1.7 (1) | -241.9 (17.98) | 0.82 | 91.62 | 91.21 |
| **l8.1** | **TG + DOSE + SP + AGE + ST + DPI + SEX** | **-2.77 (1.25)** | **-241.02 (18.06)** | **-2.87 (1.13)** | **-243.07 (18.07)** | **0.82** | **91.68** | **91.12** |

**Table S2. Extended sgRNA logistic model performance comparisons.** Models are ordered by increasing number of predictors, with the simplest (l1), best (l4.2), and full (l8.1) models noted in bold. We report expected log pointwise predictive density (ELPD) generated by 10-fold cross validation (cross-validation columns), where larger ELPD indicates better performance. ELPD difference indicates the difference between ELPDs of the given model and the model with the largest ELPD (in this case model l6.1, though this is not our 'best model'). The PSIS-LOO approximation columns present statistics generated by running Pareto-Smoothed Importance Sampling approximate leave-one-out cross validation, including ELPD and ELPD difference as above. The prediction columns indicate the percent of samples (stratified by training and test sets) for which posterior predictions generated by 10-fold cross validation correctly classified them as below or above the limit of detection (i.e., where the per-sample posterior predictive distributions exhibited at least a probability of 0.5 for the true, observed classification). MCC is the Matthews correlation coefficient. Note that all models included total RNA as a predictor, even though it is not specified in the predictor column. Standard error (SE) is shown in parentheses following all relevant statistics.

| | | *Cross-validation* | | *PSIS-LOO Approximation* | | *Prediction* | | | | | |
|---|---|---|---|---|---|---|---|---|---|---|---|
| **Model** | **Predictors** | **ELPD Difference (SE)** | **ELPD (SE)** | **ELPD Difference (SE)** | **ELPD (SE)** | **MAE (scaled)** | | **% within 50% PI** | | **% within 95% PI** | |
| | | | | | | **train** | **test** | **train** | **test** | **train** | **test** |
| **f1** | | **-186.24 (17.05)** | **-1017.32 (29.41)** | **-196.49 (21.07)** | **-1017.12 (29.35)** | **0.58 (0.7)** | **0.58 (0.7)** | **47.5** | **48** | **94.9** | **94.8** |
| f2.1 | DOSE | -114.04 (11.7) | -945.12 (32.24) | -111.91 (15.27) | -932.54 (30.49) | 0.6 (0.65) | 0.6 (0.65) | 46.9 | 46.6 | 95.9 | 95.6 |
| f2.2 | ST | -98.38 (10.49) | -929.46 (32.38) | -95.87 (13.7) | -916.5 (30.74) | 0.56 (0.63) | 0.56 (0.63) | 49.9 | 50.3 | 96.5 | 96 |
| f2.3 | SP | -109.47 (11.8) | -940.55 (32.74) | -106.94 (15.43) | -927.56 (31.08) | 0.56 (0.61) | 0.57 (0.62) | 50.7 | 49.4 | 96.2 | 95.8 |
| f2.4 | AGE | -107.02 (12.6) | -938.1 (32.65) | -104.67 (16.34) | -925.29 (30.91) | 0.57 (0.62) | 0.57 (0.64) | 50.7 | 49.6 | 96.9 | 96.5 |
| f2.5 | SEX | -120.9 (12.58) | -951.98 (32.98) | -129.78 (17.11) | -950.41 (32.6) | 0.56 (0.62) | 0.58 (0.64) | 50.6 | 48.5 | 96.2 | 95.8 |
| f2.6 | DPI | -82.23 (9.7) | -913.31 (31.99) | -79.37 (12.66) | -900 (30.31) | 0.56 (0.65) | 0.55 (0.64) | 47.9 | 47.5 | 96.1 | 96 |
| f2.7 | TG | -94.6 (11.02) | -925.68 (33.96) | -93.29 (14.53) | -913.91 (32.33) | 0.55 (0.61) | 0.57 (0.63) | 50.7 | 50.4 | 97.8 | 97.7 |
| f3.1 | DPI + DOSE | -83.66 (9.65) | -914.74 (32.05) | -81.44 (12.63) | -902.06 (30.39) | 0.56 (0.65) | 0.56 (0.65) | 48 | 48.5 | 96.4 | 96 |
| f3.2 | DPI + ST | -82.78 (9.59) | -913.86 (32.07) | -80.27 (12.56) | -900.9 (30.46) | 0.55 (0.65) | 0.55 (0.65) | 47.6 | 47.8 | 96.1 | 96 |
| f3.3 | DPI + SP | -78.12 (9.53) | -909.2 (32.18) | -76.32 (12.54) | -896.95 (30.58) | 0.55 (0.65) | 0.55 (0.65) | 48.1 | 48.5 | 96 | 95.8 |
| f3.4 | DPI + AGE | -74.78 (10.35) | -905.86 (32.11) | -72.99 (13.5) | -893.62 (30.54) | 0.54 (0.64) | 0.54 (0.65) | 49.6 | 48.9 | 96.3 | 96.2 |
| f3.5 | DPI + SEX | -87.71 (10.23) | -918.79 (32.22) | -91.38 (13.37) | -912 (30.95) | 0.54 (0.64) | 0.57 (0.67) | 48.9 | 46.6 | 95.9 | 95.5 |
| f3.6 | DPI + TG | -48.36 (8.3) | -879.44 (34.01) | -46.14 (10.88) | -866.77 (32.38) | 0.51 (0.61) | 0.52 (0.6) | 50.8 | 51 | 97.7 | 97.7 |
| f4.1 | DPI + TG + DOSE | -40.6 (8.79) | -871.68 (35.31) | -38.35 (11.32) | -858.98 (33.91) | 0.49 (0.54) | 0.49 (0.53) | 52.1 | 52.2 | 96.6 | 96.7 |
| f4.2 | DPI + TG + ST | -44.28 (7.39) | -875.36 (34.38) | -42.03 (9.66) | -862.65 (32.86) | 0.51 (0.6) | 0.51 (0.59) | 51.3 | 51.7 | 97.7 | 97.6 |
| f4.3 | DPI + TG + SP | -2.82 (3.66) | -833.9 (34.41) | -1.33 (4.81) | -821.95 (33.02) | 0.46 (0.57) | 0.47 (0.57) | 53.1 | 52.7 | 96.5 | 97 |
| f4.4 | DPI + TG + AGE | -36.03 (7.45) | -867.11 (33.05) | -34.98 (9.47) | -855.6 (31.27) | 0.48 (0.56) | 0.49 (0.59) | 53.8 | 52.9 | 97 | 96.7 |
| f4.5 | DPI + TG + SEX | -52.59 (8.54) | -883.67 (34.03) | -53.68 (11.25) | -874.3 (32.66) | 0.5 (0.6) | 0.52 (0.61) | 51.6 | 50.8 | 97.6 | 97.4 |
| **f5.1** | **DPI + TG + SP + DOSE** | **0 (0)** | **-831.08 (34.5)** | **0 (0)** | **-820.63 (33.24)** | **0.43 (0.53)** | **0.44 (0.54)** | **56** | **55** | **97** | **96.9** |
| f5.2 | DPI + TG + SP + ST | -3.41 (3.65) | -834.49 (34.35) | -1.83 (4.82) | -822.45 (32.99) | 0.46 (0.57) | 0.48 (0.58) | 52.8 | 52.7 | 96.4 | 96.7 |
| f5.3 | DPI + TG + SP + AGE | -4.88 (3.34) | -835.96 (34.24) | -4.13 (4.51) | -824.76 (32.79) | 0.46 (0.57) | 0.48 (0.58) | 53.5 | 53.1 | 96.6 | 96.7 |
| f5.4 | DPI + TG + SP + SEX | -4.64 (3.6) | -835.72 (34.36) | -7.22 (4.65) | -827.85 (33.13) | 0.45 (0.57) | 0.48 (0.59) | 53.4 | 52.4 | 96.5 | 96 |
| f6.1 | DPI + TG + SP + DOSE + ST | -1.11 (0.32) | -832.19 (34.53) | -0.68 (0.31) | -821.3 (33.14) | 0.43 (0.53) | 0.44 (0.54) | 56 | 54.5 | 97 | 96.7 |
| f6.2 | DPI + TG + SP + DOSE + AGE | -1.51 (1.17) | -832.59 (34.4) | -1.01 (1.9) | -821.64 (32.86) | 0.43 (0.53) | 0.43 (0.53) | 56.5 | 55.3 | 97 | 96.7 |
| f6.3 | DPI + TG + SP + DOSE + SEX | -1.5 (0.69) | -832.58 (34.48) | -5.76 (0.85) | -826.39 (33.37) | 0.43 (0.53) | 0.44 (0.54) | 56.6 | 54.5 | 96.9 | 96.7 |
| f7.1 | DPI + TG + SP + DOSE + ST + AGE | -2.64 (1.22) | -833.72 (34.4) | -2.56 (1.85) | -823.19 (33.01) | 0.43 (0.52) | 0.43 (0.53) | 56.4 | 55.1 | 97 | 96.7 |
| f7.2 | DPI + TG + SP + DOSE + ST + SEX | -2.72 (0.76) | -833.8 (34.54) | -6.76 (0.88) | -827.39 (33.4) | 0.43 (0.53) | 0.44 (0.54) | 56.4 | 54.1 | 96.9 | 96.9 |
| **f8.1** | **DPI + TG + SP + DOSE + ST + AGE + SEX** | **-4.44 (2.39)** | **-835.52 (34.2)** | **-9.76 (4.12)** | **-830.38 (33.19)** | **0.43 (0.53)** | **0.45 (0.56)** | **57.2** | **54.5** | **96.9** | **96.7** |

**Table S3. Extended sgRNA linear model performance comparisons.** Models are ordered by increasing number of predictors, with the simplest (f1), best (f5.1), and full (f8.1) models noted in bold. We report expected log pointwise predictive density (ELPD) generated by 10-fold cross validation (cross-validation columns), where larger ELPD indicates better performance. The top logistic model was run in tandem with all tested linear components, so the ELPD reported here reflects the sum of the ELPD for the top logistic and the considered linear components. ELPD difference indicates the difference between ELPDs of the given model and the model with the largest ELPD (in this case model l5.1, the 'best model'). The PSIS-LOO approximation columns present statistics generated by running Pareto-Smoothed Importance Sampling approximate leave-one-out cross validation, including ELPD and ELPD difference. Standard error (SE) is shown in parentheses following all relevant statistics. We also used multiple metrics to assess model predictions, which are all stratified by performance on training versus test data sets and were generated by 10-fold cross validation. MAE is the median difference between the observed value and the posterior predictive median (i.e., median absolute error around the median) for all samples with sgRNA above the LOD, and this metric was also scaled by one standard deviation (Scaled). '% within 50% PI' and '% within 95% PI' columns indicate the percent of sgRNA positive samples where the true, observed value fell within the sample-specific 50% and 95% prediction intervals, respectively. Note that all models included total RNA as a predictor, even though it is not specified in the predictor column.

| Model | Predictors | Cross-validation ELPD Difference (SE) | Cross-validation ELPD (SE) | PSIS-LOO Approximation ELPD Difference (SE) | PSIS-LOO Approximation ELPD (SE) | Prediction MCC | Prediction % correctly predicted train | Prediction % correctly predicted test |
|---|---|---|---|---|---|---|---|---|
| **c1** | **T** | **-57.32 (14.47)** | **-434.28 (12.5)** | **-44.26 (8.66)** | **-434.2 (12.56)** | **0.48** | **81.9** | **81.7** |
| c2.1 | T + CELL | -56.61 (14.45) | -433.58 (12.83) | -42.96 (8.91) | -432.9 (12.88) | 0.51 | 82.6 | 82.5 |
| c2.2 | T + ASSAY | -45 (12.78) | -421.96 (12.88) | -32.05 (7.62) | -421.99 (12.95) | 0.53 | 83.3 | 83.1 |
| c2.3 | T + DOSE | -58.54 (14.58) | -435.5 (12.63) | -45.22 (8.68) | -435.16 (12.65) | 0.48 | 81.8 | 81.7 |
| c2.4 | T + ST | -56.44 (13.99) | -433.41 (12.34) | -42.62 (8.53) | -432.56 (12.38) | 0.45 | 81 | 80.5 |
| c2.5 | T + SP | -50.39 (13.4) | -427.35 (12.63) | -37.65 (8.02) | -427.59 (12.69) | 0.47 | 81.2 | 81 |
| c2.6 | T + AGE | -43.81 (12.94) | -420.77 (13.36) | -44.83 (8.31) | -434.77 (14.43) | 0.48 | 81.5 | 81.6 |
| c2.7 | T + SEX | -53.21 (13.65) | -430.17 (12.63) | -39.79 (8.2) | -429.73 (12.69) | 0.48 | 81.6 | 81.7 |
| c2.8 | T + DPI | -38.17 (12.08) | -415.14 (13.29) | -25.16 (6.99) | -415.1 (13.32) | 0.49 | 82.3 | 82.1 |
| c2.9 | T + TG | -54.07 (14.1) | -431.03 (13.27) | -41.53 (8.26) | -431.47 (13.32) | 0.52 | 83 | 82.9 |
| c3.1 | T + DPI + CELL | -39.93 (11.97) | -416.89 (13.43) | -26.62 (6.94) | -416.56 (13.48) | 0.49 | 82.3 | 81.9 |
| c3.2 | T + DPI + ASSAY | -34.81 (11.29) | -411.77 (13.46) | -21.83 (6.45) | -411.77 (13.5) | 0.53 | 83.2 | 83.3 |
| c3.3 | T + DPI + DOSE | -39.4 (12.33) | -416.36 (13.58) | -26.1 (6.98) | -416.04 (13.58) | 0.49 | 82.1 | 82 |
| c3.4 | T + DPI + ST | -36.89 (11.47) | -413.85 (13.25) | -23.7 (6.75) | -413.65 (13.3) | 0.48 | 82 | 81.7 |
| c3.5 | T + DPI + SP | -30.04 (10.37) | -407 (13.34) | -17.39 (5.84) | -407.33 (13.41) | 0.52 | 82.9 | 82.7 |
| c3.6 | T + DPI + AGE | -26.08 (9.99) | -403.04 (13.99) | -26.6 (6.39) | -416.54 (15.01) | 0.5 | 82.2 | 82.3 |
| c3.7 | T + DPI + SEX | -33.16 (10.83) | -410.12 (13.38) | -19.74 (6.24) | -409.68 (13.4) | 0.5 | 82.3 | 82.1 |
| c3.8 | T + DPI + TG | -37.17 (11.6) | -414.13 (13.88) | -24.37 (6.52) | -414.32 (13.93) | 0.51 | 82.9 | 82.8 |
| c4.1 | T + DPI + AGE + CELL | -27.26 (9.94) | -404.22 (14.09) | -28.6 (6.74) | -418.54 (15.27) | 0.5 | 82.4 | 82.2 |
| c4.2 | T + DPI + AGE + ASSAY | -20.97 (8.63) | -397.93 (14.24) | -23.2 (5.8) | -413.14 (15.44) | 0.54 | 84.1 | 83.7 |
| c4.3 | T + DPI + AGE + DOSE | -27.09 (10.2) | -404.05 (14.18) | -27.87 (6.44) | -417.81 (15.24) | 0.5 | 82.4 | 82.4 |
| c4.4 | T + DPI + AGE + ST | -27.07 (10.02) | -404.03 (14.04) | -27.71 (6.41) | -417.65 (15.07) | 0.5 | 82.2 | 82.3 |
| c4.5 | T + DPI + AGE + SP | -20.32 (8.28) | -397.28 (14.21) | -19.06 (5.05) | -409 (15.05) | 0.5 | 82.9 | 82.3 |
| c4.6 | T + DPI + AGE + SEX | -23.74 (9.1) | -400.7 (14.11) | -21.94 (5.56) | -411.88 (14.92) | 0.5 | 82.9 | 82.3 |
| c4.7 | T + DPI + AGE + TG | -19.42 (7.68) | -396.38 (14.66) | -29.38 (6.66) | -419.32 (16.85) | 0.54 | 83.9 | 83.8 |
| c5.1 | T + DPI + AGE + TG + CELL | -20.13 (7.58) | -397.09 (14.63) | -32.11 (7.53) | -422.05 (17.21) | 0.54 | 84.3 | 83.6 |
| c5.2 | T + DPI + AGE + TG + ASSAY | -17.58 (7.13) | -394.54 (14.82) | -24.06 (5.78) | -414 (16.55) | 0.56 | 84.5 | 84.5 |
| c5.3 | T + DPI + AGE + TG + DOSE | -20.39 (7.9) | -397.35 (14.85) | -30.78 (6.8) | -420.73 (17.13) | 0.55 | 83.9 | 84 |
| c5.4 | T + DPI + AGE + TG + ST | -20.11 (7.52) | -397.07 (14.77) | -28.8 (6.36) | -418.74 (16.79) | 0.54 | 83.9 | 83.7 |
| c5.5 | T + DPI + AGE + TG + SP | -18.93 (7.2) | -395.89 (14.71) | -25.99 (5.7) | -415.93 (16.5) | 0.56 | 84 | 84.2 |
| c5.6 | T + DPI + AGE + TG + SEX | -18.29 (6.87) | -395.25 (14.79) | -25.23 (5.62) | -415.17 (16.56) | 0.55 | 84.1 | 84.1 |
| c6.1 | T + DPI + AGE + TG + ASSAY + CELL | -19.07 (7.23) | -396.03 (14.83) | -27.82 (6.48) | -417.76 (16.89) | 0.55 | 84.8 | 84.1 |
| c6.2 | T + DPI + AGE + TG + ASSAY + DOSE | -16.79 (7.13) | -393.75 (15.15) | -23.08 (5.5) | -413.02 (16.85) | 0.54 | 84.3 | 83.8 |

| | | | | | | | | |
|---|---|---|---|---|---|---|---|---|
| c6.3 | T + DPI + AGE + TG + ASSAY + ST | -18.54 (7.11) | -395.5 (14.9) | -24.69 (5.76) | -414.63 (16.6) | 0.56 | 84.6 | 84.2 |
| c6.4 | T + DPI + AGE + TG + ASSAY + SP | -7.88 (5.11) | -384.84 (14.74) | -5.17 (2.87) | -395.11 (15.4) | 0.57 | 84.9 | 84.6 |
| c6.5 | T + DPI + AGE + TG + ASSAY + SEX | -16.07 (6.08) | -393.03 (14.93) | -19.51 (4.6) | -409.45 (16.28) | 0.55 | 84.4 | 84.1 |
| c7.1 | T + DPI + AGE + TG + ASSAY + SP + CELL | -9.5 (5.15) | -386.46 (14.69) | -8.23 (3.39) | -398.18 (15.49) | 0.56 | 85 | 84.3 |
| c7.2 | T + DPI + AGE + TG + ASSAY + SP + DOSE | -5.18 (4.83) | -382.14 (15.06) | -2.2 (1.65) | -392.14 (15.68) | 0.58 | 85.1 | 85 |
| c7.3 | T + DPI + AGE + TG + ASSAY + SP + ST | -8.26 (5.12) | -385.22 (14.85) | -4.88 (2.83) | -394.82 (15.43) | 0.56 | 84.9 | 84.3 |
| c7.4 | T + DPI + AGE + TG + ASSAY + SP + SEX | -5.72 (3.81) | -382.68 (14.86) | -0.93 (1.58) | -390.87 (15.35) | 0.57 | 85.3 | 84.7 |
| **c8.1** | **T + DPI + AGE + TG + ASSAY + SP + DOSE + CELL** | **-1.62 (3.25)** | **-378.59 (15.12)** | **-4.38 (4.11)** | **-394.32 (16.45)** | **0.57** | **85.4** | **84.7** |
| c8.2 | T + DPI + AGE + TG + ASSAY + SP + DOSE + ST | -5.77 (4.88) | -382.73 (15.14) | -1.98 (1.62) | -391.93 (15.74) | 0.58 | 85.2 | 85 |
| c8.3 | T + DPI + AGE + TG + ASSAY + SP + DOSE + SEX | -4.57 (4.14) | -381.54 (15.14) | 0 (0) | -389.94 (15.63) | 0.57 | 85.1 | 84.8 |
| c9.1 | T + DPI + AGE + TG + ASSAY + SP + DOSE + CELL + ST | -2.33 (3.3) | -379.29 (15.2) | -4.74 (4.05) | -394.69 (16.51) | 0.58 | 85.5 | 84.9 |
| c9.2 | T + DPI + AGE + TG + ASSAY + SP + DOSE + CELL + SEX | 0 (0) | -376.96 (15.2) | -0.76 (3.27) | -390.7 (16.31) | 0.57 | 85.8 | 84.6 |
| **c10.1** | **T + DPI + AGE + TG + ASSAY + SP + DOSE + CELL + SEX + ST** | **-0.87 (0.81)** | **-377.83 (15.28)** | **-1.67 (3.29)** | **-391.61 (16.39)** | **0.57** | **85.8** | **84.7** |

**Table S4. Extended culture model performance comparisons with totRNA as the primary predictor.** Models are ordered by increasing number of predictors, with the simplest (c1), best (c8.1), and full (c10.1) models noted in bold. We report expected log pointwise predictive density (ELPD) generated by 10-fold cross validation (cross-validation columns), where larger ELPD indicates better performance. ELPD difference indicates the difference between ELPDs of the given model and the model with the largest ELPD (in this case model l9.2, though this is not our 'best model'). The PSIS-LOO approximation columns present statistics generated by running Pareto-Smoothed Importance Sampling approximate leave-one-out cross validation, including ELPD and ELPD difference. The prediction column indicates the percent of samples (stratified by training and test sets) for which posterior predictions generated by 10-fold cross validation correctly classified them as below or above the limit of detection (i.e., where the per-sample posterior predictive distributions exhibited at least a probability of 0.5 for the true, observed classification). Standard error (SE) is shown in parentheses following all relevant statistics.

| Model | Predictors | *Cross-validation* ELPD Difference (SE) | *Cross-validation* ELPD (SE) | *PSIS-LOO Approximation* ELPD Difference (SE) | *PSIS-LOO Approximation* ELPD (SE) | *Prediction* MCC | *Prediction* % correctly predicted train | *Prediction* % correctly predicted test |
|---|---|---|---|---|---|---|---|---|
| **c1** | **SG** | **-48.77 (17.87)** | **-350.19 (7.87)** | **-47.12 (9.22)** | **-350.38 (7.97)** | **0.43** | **80.8** | **80.8** |
| c2.1 | SG + CELL | -46.15 (17.66) | -347.58 (8.24) | -45.15 (9.11) | -348.41 (8.36) | 0.46 | 81.5 | 81.6 |
| c2.2 | SG + ASSAY | -43.94 (17.39) | -345.37 (8.14) | -42.73 (8.97) | -345.98 (8.26) | 0.44 | 81.1 | 80.9 |
| c2.3 | SG + DOSE | -49.75 (18.01) | -351.18 (8.13) | -48.24 (9.27) | -351.5 (8.2) | 0.42 | 80.6 | 80.5 |
| c2.4 | SG + ST | -44.12 (16.49) | -345.55 (8.51) | -42.39 (8.48) | -345.64 (8.65) | 0.47 | 82.1 | 81.9 |
| c2.5 | SG + SP | -47.13 (16.95) | -348.56 (8.21) | -45.76 (8.76) | -349.02 (8.31) | 0.43 | 81.1 | 80.9 |
| c2.6 | SG + AGE | -47.52 (17.64) | -348.94 (8.27) | -46.21 (9.12) | -349.47 (8.34) | 0.44 | 80.9 | 80.9 |
| c2.7 | SG + SEX | -48.93 (17.77) | -350.35 (8.13) | -47.77 (9.19) | -351.03 (8.24) | 0.44 | 81.3 | 81.1 |
| c2.8 | SG + DPI | -16.3 (10.84) | -317.73 (10.23) | -14.63 (5.69) | -317.89 (10.32) | 0.52 | 83.5 | 83.4 |
| c2.9 | SG + TG | -49.06 (17.9) | -350.49 (8.11) | -47.51 (9.23) | -350.77 (8.19) | 0.43 | 80.7 | 80.8 |
| c3.1 | SG + DPI + CELL | -15.02 (10.4) | -316.44 (10.4) | -13.52 (5.47) | -316.78 (10.51) | 0.54 | 84 | 84 |
| c3.2 | SG + DPI + ASSAY | -13.09 (9.9) | -314.52 (10.36) | -11.79 (5.18) | -315.05 (10.47) | 0.52 | 83.8 | 83.6 |
| c3.3 | SG + DPI + DOSE | -17.31 (11.11) | -318.74 (10.42) | -15.68 (5.81) | -318.93 (10.47) | 0.51 | 83.6 | 83.3 |
| c3.4 | SG + DPI + ST | -16.67 (10.63) | -318.1 (10.31) | -15.35 (5.58) | -318.61 (10.4) | 0.52 | 83.5 | 83.5 |
| c3.5 | SG + DPI + SP | -17.26 (10.42) | -318.68 (10.34) | -15.78 (5.46) | -319.04 (10.46) | 0.51 | 83.6 | 83 |
| c3.6 | SG + DPI + AGE | -16.06 (10.62) | -317.49 (10.56) | -14.41 (5.57) | -317.67 (10.6) | 0.52 | 83.4 | 83.6 |
| c3.7 | SG + DPI + SEX | -16.15 (10.48) | -317.57 (10.45) | -14.71 (5.53) | -317.97 (10.51) | 0.53 | 83.9 | 83.7 |
| c3.8 | SG + DPI + TG | -15.41 (10.6) | -316.83 (10.55) | -14 (5.57) | -317.26 (10.63) | 0.52 | 83.8 | 83.5 |
| c4.1 | SG + DPI + ASSAY + CELL | -13.64 (9.66) | -315.06 (10.51) | -12.25 (5.08) | -315.5 (10.63) | 0.52 | 84 | 83.6 |
| c4.2 | SG + DPI + ASSAY + DOSE | -14.07 (10.26) | -315.49 (10.75) | -12.81 (5.33) | -316.07 (10.82) | 0.53 | 83.8 | 83.7 |
| c4.3 | SG + DPI + ASSAY + ST | -10.94 (8.41) | -312.36 (10.56) | -9.72 (4.42) | -312.98 (10.7) | 0.53 | 84.4 | 83.8 |
| c4.4 | SG + DPI + ASSAY + SP | -14.16 (9.73) | -315.59 (10.53) | -13.13 (5.08) | -316.39 (10.68) | 0.52 | 83.9 | 83.5 |
| c4.5 | SG + DPI + ASSAY + AGE | -10.83 (8.96) | -312.26 (10.81) | -9.42 (4.68) | -312.67 (10.9) | 0.52 | 84.3 | 83.7 |
| c4.6 | SG + DPI + ASSAY + SEX | -13.32 (9.61) | -314.74 (10.58) | -12.4 (5.06) | -315.66 (10.68) | 0.52 | 83.9 | 83.4 |
| c4.7 | SG + DPI + ASSAY + TG | -12.51 (9.7) | -313.93 (10.66) | -11.52 (5.09) | -314.78 (10.77) | 0.53 | 83.9 | 83.8 |
| c5.1 | SG + DPI + ASSAY + AGE + CELL | -11.65 (8.76) | -313.07 (10.92) | -10.11 (4.62) | -313.36 (11) | 0.52 | 84.2 | 83.7 |
| c5.2 | SG + DPI + ASSAY + AGE + DOSE | -11.86 (9.18) | -313.28 (11.06) | -10.32 (4.78) | -313.58 (11.1) | 0.52 | 84.1 | 83.5 |
| c5.3 | SG + DPI + ASSAY + AGE + ST | -9.45 (7.96) | -310.87 (11.01) | -8.18 (4.19) | -311.43 (11.13) | 0.54 | 84.3 | 84.2 |
| c5.4 | SG + DPI + ASSAY + AGE + SP | -10.83 (8.15) | -312.25 (10.94) | -9.82 (4.29) | -313.08 (11.07) | 0.53 | 84.2 | 83.8 |
| c5.5 | SG + DPI + ASSAY + AGE + SEX | -10.02 (8.18) | -311.44 (10.99) | -9.01 (4.29) | -312.27 (11.07) | 0.52 | 84.5 | 83.7 |
| c5.6 | SG + DPI + ASSAY + AGE + TG | -10.65 (8.79) | -312.07 (11.01) | -9.56 (4.62) | -312.82 (11.13) | 0.54 | 84.3 | 84.1 |
| c6.1 | SG + DPI + ASSAY + AGE + ST + CELL | -10.58 (7.89) | -312.01 (11.14) | -9.22 (4.17) | -312.48 (11.25) | 0.53 | 84.2 | 83.8 |
| c6.2 | SG + DPI + ASSAY + AGE + ST + DOSE | -10.03 (7.98) | -311.46 (11.3) | -8.65 (4.17) | -311.91 (11.39) | 0.54 | 84.6 | 84.2 |

| | | | | | | | | |
|---|---|---|---|---|---|---|---|---|
| c6.3 | SG + DPI + ASSAY + AGE + ST + SP | -9.89 (7.24) | -311.31 (11.13) | -8.99 (3.84) | -312.25 (11.31) | 0.53 | 84.5 | 84 |
| c6.4 | SG + DPI + ASSAY + AGE + ST + SEX | -8.37 (6.83) | -309.8 (11.19) | -7.57 (3.62) | -310.83 (11.28) | 0.53 | 84.4 | 83.9 |
| c6.5 | SG + DPI + ASSAY + AGE + ST + TG | -8.33 (7.55) | -309.76 (11.33) | -7.31 (4.01) | -310.57 (11.48) | 0.54 | 84.5 | 84.1 |
| c7.1 | SG + DPI + ASSAY + AGE + ST + TG + CELL | -9.56 (7.57) | -310.99 (11.47) | -8.36 (4.02) | -311.62 (11.61) | 0.53 | 84.5 | 83.8 |
| c7.2 | SG + DPI + ASSAY + AGE + ST + TG + DOSE | -8.94 (7.49) | -310.37 (11.51) | -7.84 (3.95) | -311.1 (11.61) | 0.54 | 84.7 | 84.2 |
| c7.3 | SG + DPI + ASSAY + AGE + ST + TG + SP | -9.35 (7.08) | -310.77 (11.39) | -8.76 (3.78) | -312.02 (11.61) | 0.54 | 84.7 | 84.3 |
| c7.4 | SG + DPI + ASSAY + AGE + ST + TG + SEX | -7.14 (6.32) | -308.56 (11.46) | -6.8 (3.39) | -310.06 (11.63) | 0.54 | 84.7 | 84.3 |
| c8.1 | SG + DPI + ASSAY + AGE + ST + TG + SEX + CELL | -7.25 (5.87) | -308.67 (11.57) | -6.67 (3.13) | -309.93 (11.69) | 0.56 | 85 | 84.6 |
| c8.2 | SG + DPI + ASSAY + AGE + ST + TG + SEX + DOSE | -7.92 (6.52) | -309.35 (11.57) | -7.34 (3.45) | -310.6 (11.7) | 0.55 | 84.8 | 84.5 |
| c8.3 | SG + DPI + ASSAY + AGE + ST + TG + SEX + SP | -8.19 (5.88) | -309.62 (11.54) | -8.03 (3.19) | -311.29 (11.76) | 0.55 | 84.9 | 84.4 |
| c9.1 | SG + DPI + ASSAY + AGE + ST + TG + SEX + CELL + DOSE | -2.88 (4.95) | -304.3 (11.8) | -2.43 (2.57) | -305.69 (11.95) | 0.55 | 85.2 | 84.6 |
| c9.2 | SG + DPI + ASSAY + AGE + ST + TG + SEX + CELL + SP | -5.56 (3.71) | -306.99 (11.64) | -5.3 (2.01) | -308.56 (11.85) | 0.56 | 85.2 | 84.8 |
| **c10.1** | **SG + DPI + ASSAY + AGE + ST + TG + SEX + CELL + DOSE + SP** | **0 (0)** | **-301.42 (11.91)** | **0 (0)** | **-303.26 (12.15)** | **0.56** | **85.7** | **84.9** |

**Table S5. Extended culture model performance comparisons with sgRNA as the primary predictor.** Models are ordered by increasing number of predictors, with the simplest (c1) and best/full model (c10.1) models noted in bold. We report expected log pointwise predictive density (ELPD) generated by 10-fold cross validation (cross-validation columns), where larger ELPD indicates better performance. ELPD difference indicates the difference between ELPDs of the given model and the model with the largest ELPD (in this case model c10.1, our 'best model'). The PSIS-LOO approximation columns present statistics generated by running Pareto-Smoothed Importance Sampling approximate leave-one-out cross validation, including ELPD and ELPD difference. The prediction column indicates the percent of samples (stratified by training and test sets) for which posterior predictions generated by 10-fold cross validation correctly classified them as below or above the limit of detection (i.e., where the per-sample posterior predictive distributions exhibited at least a probability of 0.5 for the true, observed classification). Standard error (SE) is shown in parentheses following all relevant statistics.

| | | TotRNA | 0 | 1 | 2 | 3 | 4 | 5 | 6 | 7 | 8 | 9 | 10 |
|---|---|---|---|---|---|---|---|---|---|---|---|---|---|
| sgRNA Logistic | Dose | 4 | 0, 0 | 0, 1 | 1, 4 | 9, 19 | 43, 61 | 84, 92 | 97, 99 | 100, 100 | 100, 100 | 100, 100 | 100, 100 |
| | | 5.5 | 0, 0 | 0, 0 | 0, 1 | 2, 6 | 16, 27 | 6, 72 | 91, 95 | 98, 99 | 100, 100 | 100, 100 | 100, 100 |
| | | 7 | 0, 0 | 0, 0 | 0, 0 | 0, 2 | 4, 10 | 23, 43 | 69, 84 | 94, 98 | 99, 100 | 100, 100 | 100, 100 |
| | Species | RM | 0, 0 | 0, 0 | 0, 1 | 2, 6 | 16, 27 | 6, 72 | 91, 95 | 98, 99 | 100, 100 | 100, 100 | 100, 100 |
| | | CM | 0, 0 | 0, 0 | 0, 1 | 2, 7 | 12, 33 | 5, 77 | 87, 96 | 98, 100 | 100, 100 | 100, 100 | 100, 100 |
| | | AGM | 0, 0 | 0, 0 | 0, 0 | 0, 2 | 3, 12 | 17, 49 | 59, 88 | 91, 98 | 98, 100 | 100, 100 | 100, 100 |
| | Target Gene | T↑SG↑ | 0, 3 | 3, 17 | 18, 58 | 61, 91 | 91, 99 | 99, 100 | 100, 100 | 100, 100 | 100, 100 | 100, 100 | 100, 100 |
| | | T↓SG↑ | 0, 1 | 1, 8 | 6, 38 | 3, 81 | 75, 97 | 95, 100 | 99, 100 | 100, 100 | 100, 100 | 100, 100 | 100, 100 |
| | | T↑SG↓ | 0, 0 | 0, 0 | 0, 1 | 2, 6 | 16, 27 | 60, 72 | 91, 95 | 98, 99 | 100, 100 | 100, 100 | 100, 100 |
| | | T↓SG↓ | 0, 0 | 0, 0 | 0, 3 | 4, 15 | 24, 55 | 69, 90 | 94, 98 | 99, 100 | 100, 100 | 100, 100 | 100, 100 |
| sgRNA Linear | Dose | 4 | -1.16, -0.5 | -0.23, 0.39 | 0.7, 1.28 | 1.63, 2.17 | 2.55, 3.07 | 3.47, 3.97 | 4.38, 4.87 | 5.29, 5.79 | 6.2, 6.7 | 7.1, 7.62 | 8, 8.55 |
| | | 5.5 | -1.4, -0.88 | -0.46, 0 | 0.48, 0.89 | 1.41, 1.77 | 2.34, 2.66 | 3.27, 3.56 | 4.18, 4.46 | 5.09, 5.37 | 5.99, 6.29 | 6.88, 7.22 | 7.77, 8.15 |
| | | 7 | -1.76, -1.14 | -0.83, -0.25 | 0.1, 0.64 | 1.02, 1.53 | 1.94, 2.43 | 2.86, 3.33 | 3.77, 4.24 | 4.68, 5.15 | 5.58, 6.06 | 6.48, 6.98 | 7.38, 7.91 |
| | DPI | I, 1 | -1.96, -1.39 | -1.02, -0.5 | -0.09, 0.38 | 0.84, 1.28 | 1.77, 2.17 | 2.69, 3.06 | 3.61, 3.97 | 4.52, 4.88 | 5.42, 5.79 | 6.32, 6.71 | 7.2, 7.64 |
| | | I, 2+ | -1.4, -0.88 | -0.46, 0 | 0.48, 0.89 | 1.41, 1.77 | 2.34, 2.66 | 3.27, 3.56 | 4.18, 4.46 | 5.09, 5.37 | 5.99, 6.29 | 6.88, 7.22 | 7.77, 8.15 |
| | | NI, 1+ | -1.01, -0.41 | -0.08, 0.47 | 0.86, 1.35 | 1.79, 2.24 | 2.72, 3.13 | 3.65, 4.03 | 4.57, 4.92 | 5.48, 5.83 | 6.38, 6.74 | 7.28, 7.66 | 8.18, 8.58 |
| | Target Gene | T↓SG↑ | 0.01, 0.6 | 0.94, 1.49 | 1.87, 2.38 | 2.79, 3.28 | 3.71, 4.18 | 4.62, 5.09 | 5.53, 6 | 6.43, 6.91 | 7.33, 7.83 | 8.22, 8.76 | 9.11, 9.69 |
| | | T↑SG↓ | -1.4, -0.88 | -0.46, 0 | 0.48, 0.89 | 1.41, 1.77 | 2.34, 2.66 | 3.27, 3.56 | 4.18, 4.46 | 5.09, 5.37 | 5.99, 6.29 | 6.88, 7.22 | 7.77, 8.15 |
| | | T↓SG↓ | -1.05, -0.48 | -0.11, 0.4 | 0.81, 1.29 | 1.74, 2.19 | 2.66, 3.08 | 3.57, 3.99 | 4.48, 4.9 | 5.39, 5.82 | 6.28, 6.73 | 7.18, 7.66 | 8.07, 8.59 |
| | Species | RM | -1.4, -0.88 | -0.46, 0 | 0.48, 0.89 | 1.41, 1.77 | 2.34, 2.66 | 3.27, 3.56 | 4.18, 4.46 | 5.09, 5.37 | 5.99, 6.29 | 6.88, 7.22 | 7.77, 8.15 |
| | | CM | -2.42, -1.74 | -1.49, -0.86 | -0.55, 0.03 | 0.38, 0.92 | 1.3, 1.81 | 2.23, 2.71 | 3.14, 3.61 | 4.05, 4.52 | 4.96, 5.43 | 5.86, 6.34 | 6.76, 7.26 |
| | | AGM | -1.36, -0.65 | -0.43, 0.23 | 0.5, 1.12 | 1.43, 2.01 | 2.35, 2.9 | 3.28, 3.8 | 4.2, 4.7 | 5.11, 5.61 | 6.02, 6.51 | 6.92, 7.43 | 7.82, 8.35 |

**Table S6. 90% Credible Intervals for the best sgRNA model.** These credible intervals correspond with the predictions in Figure 3C and H.

|  | Predictors | Prediction Accuracy (%) |  |  | MCC |
|---|---|---|---|---|---|
|  |  | Overall | Positive | Negative |  |
| Data subset | T | 90.4 | 72.2 | 94.3 | 0.67 |
|  | SG | 90.0 | 70.4 | 94.3 | 0.66 |
|  | T + SG | 90.0 | 68.5 | 94.7 | 0.65 |
| All data | T | 81.9 | 53.7 | 91.4 | 0.49 |
|  | SG*† | 80.8 | 46.7 | 91.7 | 0.43 |
|  | T + SG* | 81.7 | 51.8 | 91.7 | 0.48 |
|  | Best T | 84.7 | 60.7 | 92.8 | 0.57 |
|  | Best SG*† | 84.9 | 57.7 | 93.6 | 0.56 |

**Table S7. Performance comparison of culture models using totRNA, sgRNA, or both as the primary predictor(s).** Statistics are stratified by predictor(s) and the dataset used for fitting, including the full dataset (based on sgRNA predictions; 'all data') and the subset containing only samples with known sgRNA and totRNA results ('data subset'). Prediction accuracy reflects aggregate performance on test data across the full 10 train-test folds, stratified by all available samples (Overall), only known positive samples, and only known negative samples. MCC corresponds to the Matthews correlation coefficient. Note that we do not report ELPD because these models were fit with different quantities of data and so ELPD is not comparable. * includes imputed data. † includes data with observed sgRNA outcomes but no observed totRNA outcomes.

| | TotRNA | 0 | 1 | 2 | 3 | 4 | 5 | 6 | 7 | 8 | 9 | 10 | 11 | 12 |
|---|---|---|---|---|---|---|---|---|---|---|---|---|---|---|
| **Dose** | 4 | 0, 4 | 1, 7 | 2, 14 | 4, 26 | 10, 44 | 19, 63 | 35, 79 | 55, 90 | 72, 95 | 85, 98 | 92, 99 | 96, 100 | 98, 100 |
| | 5.5 | 0, 1 | 0, 2 | 1, 4 | 2, 7 | 4, 15 | 8, 27 | 17, 44 | 31, 64 | 50, 80 | 68, 90 | 82, 96 | 90, 98 | 95, 99 |
| | 7 | 0, 0 | 0, 1 | 0, 1 | 0, 2 | 1, 5 | 2, 10 | 5, 19 | 11, 33 | 21, 52 | 37, 71 | 56, 85 | 73, 93 | 85, 97 |
| **DPI** | I, 1 | 0, 5 | 1, 9 | 3, 18 | 6, 31 | 13, 49 | 25, 67 | 43, 82 | 63, 91 | 79, 96 | 89, 98 | 95, 99 | 97, 100 | 99, 100 |
| | I, 2+ | 0, 1 | 0, 2 | 1, 4 | 2, 7 | 4, 15 | 8, 27 | 17, 44 | 31, 64 | 50, 80 | 68, 90 | 82, 96 | 90, 98 | 95, 99 |
| | NI, 1+ | 0, 1 | 0, 2 | 1, 3 | 1, 7 | 3, 13 | 7, 24 | 15, 41 | 28, 61 | 46, 78 | 65, 89 | 80, 95 | 89, 98 | 95, 99 |
| **Species** | RM | 0, 1 | 0, 2 | 1, 4 | 2, 7 | 4, 15 | 8, 27 | 17, 44 | 31, 64 | 50, 80 | 68, 90 | 82, 96 | 90, 98 | 95, 99 |
| | CM | 0, 0 | 0, 1 | 0, 1 | 0, 2 | 1, 5 | 2, 9 | 4, 19 | 9, 33 | 19, 53 | 34, 72 | 52, 85 | 70, 93 | 83, 97 |
| | AGM | 0, 1 | 0, 2 | 1, 3 | 2, 6 | 4, 12 | 9, 21 | 20, 36 | 36, 55 | 55, 73 | 73, 87 | 85, 94 | 92, 97 | 96, 99 |
| **Age Class** | Juvenile | 0, 1 | 1, 3 | 1, 5 | 3, 10 | 6, 19 | 14, 33 | 27, 52 | 45, 71 | 64, 85 | 79, 93 | 89, 97 | 94, 99 | 97, 99 |
| | Adult | 0, 1 | 0, 2 | 1, 4 | 2, 7 | 4, 15 | 8, 27 | 17, 44 | 31, 64 | 50, 80 | 68, 90 | 82, 96 | 90, 98 | 95, 99 |
| | Geriatric | 3, 24 | 6, 40 | 13, 59 | 25, 76 | 42, 87 | 62, 94 | 78, 97 | 89, 99 | 94, 99 | 97, 100 | 99, 100 | 99, 100 | 100, 100 |
| **Cell Line** | 76 | 0, 1 | 0, 2 | 1, 4 | 2, 7 | 4, 15 | 8, 27 | 17, 44 | 31, 64 | 50, 80 | 68, 90 | 82, 96 | 90, 98 | 95, 99 |
| | E6 | 0, 0 | 0, 1 | 0, 1 | 0, 2 | 1, 4 | 2, 9 | 5, 17 | 11, 31 | 23, 49 | 39, 68 | 59, 83 | 75, 92 | 87, 96 |
| | E6-SS2 | 0, 4 | 1, 7 | 2, 14 | 4, 26 | 8, 43 | 16, 62 | 31, 78 | 49, 89 | 68, 95 | 82, 98 | 91, 99 | 95, 100 | 98, 100 |
| **Assay** | TCID50 | 1, 5 | 2, 10 | 4, 18 | 8, 33 | 17, 51 | 32, 70 | 51, 84 | 69, 92 | 83, 96 | 91, 98 | 96, 99 | 98, 100 | 99, 100 |
| | Plaque | 0, 1 | 0, 2 | 1, 4 | 2, 7 | 4, 15 | 8, 27 | 17, 44 | 31, 64 | 50, 80 | 68, 90 | 82, 96 | 90, 98 | 95, 99 |
| **Target Gene** | N | 0, 1 | 0, 2 | 1, 4 | 2, 7 | 4, 15 | 8, 27 | 17, 44 | 31, 64 | 50, 80 | 68, 90 | 82, 96 | 90, 98 | 95, 99 |
| | E | 0, 0 | 0, 1 | 0, 2 | 0, 4 | 1, 7 | 2, 15 | 3, 28 | 8, 46 | 15, 65 | 28, 81 | 47, 91 | 65, 96 | 80, 98 |
| | S | 0, 2 | 0, 4 | 1, 9 | 2, 17 | 5, 31 | 10, 49 | 20, 68 | 36, 83 | 55, 92 | 72, 96 | 85, 98 | 92, 99 | 96, 100 |

**Table S8. 90% Credible Intervals for the best culture model.** These credible intervals correspond with the predictions in Figure 5C.

| Exposure Route | Inoculated Locations | Non-inoculated Locations |
|---|---|---|
| AE | Nose/Nasopharynx, Oropharynx | Anus/Rectum |
| IT | | Anus/Rectum, Lung, Nose/Nasopharynx, Oropharynx |
| IN | Nose/Nasopharynx | Anus/Rectum, BAL, Colon, Mouth, Small intestine, Stomach |
| IG | Stomach | Anus/Rectum, Colon, Small intestine |
| OC | | Lung |
| IT, IN | BAL, Nose/Nasopharynx, Oropharynx, Trachea | Anus/Rectum, Lung, Mouth, Tonsil |
| IT, IN, OC | Eye, Nose/Nasopharynx, Oropharynx, Trachea | Anus/Rectum, Brain, Cervical LN, Colon, Kidney, Liver, Lung, Mesenteric LN, Salivary gland, Small intestine, Spleen, Tonsil |
| IT, IN, OR, OC | BAL, Bronchus, Eye, Mouth, Nose/Nasopharynx, Oropharynx, Tonsil, Trachea | Anus/Rectum, Cervical LN, Colon, Heart, Lung, Mediastinal LN, Small intestine, Stomach |

**Table S9**. **Categorization of inoculated versus non-inoculated sample locations per exposure route**. Because fluid is administered in the trachea for intratracheal (IT) inoculations, which is connected directly to the bronchioles, we include bronchus as an exposure tissue for IT inoculations. We also consider BAL an inoculated tissue for IT exposures since this procedure collects fluid from similar areas where the inoculum is administered. Exposure route abbreviations are: AE, aerosol; IT, intratracheal; IN, intranasal; IG, intragastric; OC, ocular; OR, oral.
